# Demographic Model for Inheritable Cardiac Disease

**DOI:** 10.1101/614735

**Authors:** Thomas P. Burghardt

## Abstract

The cardiac muscle proteins, generating and regulating energy transduction during a heartbeat, assemble in the sarcomere into a cyclical machine repetitively translating actin relative to myosin filaments. Myosin is the motor transducing ATP free energy into actin movement against resisting force. Cardiac myosin binding protein C (mybpc3) regulates shortening velocity probably by transient N-terminus binding to actin while its C-terminus strongly binds the myosin filament. Inheritable heart disease associated mutants frequently modify these proteins involving them in disease mechanisms. Nonsynonymous single nucleotide polymorphisms (SNPs) cause single residue substitutions with independent characteristics (sequence location, residue substitution, human demographic, and allele frequency) hypothesized to decide dependent phenotype and pathogenicity characteristics in a feed-forward Neural network model. Trial models train and validate on a dynamic worldwide SNP database for cardiac muscle proteins then predict phenotype and pathogenicity for any single residue substitution in myosin, mybpc3, or actin. A separate Bayesian model formulates conditional probabilities for phenotype or pathogenicity given independent SNP characteristics. Neural/Bayes forecasting tests SNP pathogenicity vs (in)dependent SNP characteristics to assess individualized disease risk and in particular to elucidate gender and human subpopulation bias in disease. Evident subpopulation bias in myosin SNP pathogenicities imply myosin normally engages other sarcomere proteins functionally. Consistent with this observation, mybpc3 forms a third actomyosin interaction competing with myosin essential light chain N-terminus suggesting a novel strain-dependent mechanism adapting myosin force-velocity to load dynamics. The working models, and the integral myosin/mybpc3 motor concept, portends the wider considerations involved in understanding heart disease as a systemic maladaptation.

## INTRODUCTION

The cardiac muscle proteins generate and regulate energy transduction during a heartbeat. They assemble into a cyclical machine in the sarcomere that repetitively translates actin relative to myosin filaments. Myosin is the motor transducing ATP free energy to the work of moving actin against resisting force. Cardiac myosin binding protein C (mybpc3) regulates shortening velocity probably by binding transiently to actin while stably bound to the myosin filament.

Myosin (Fig 1) has a 140 kDa N-terminal globular head called subfragment 1 (S1) and an extended α-helical tail domain (LMM+S2). Tail domains form dimers that self-assemble into myosin thick filaments with S1’s projecting outward from the core in a helical array ^2^. Thick filaments interdigitate with actin thin filaments in the sarcomere with S1’s spanning the interfilament distance. S1 also has the ATP binding site and a lever arm whose rotary movement cyclically applies tension to strongly bound actin ^3^. The lever arm complex, stabilized by bound essential and regulatory light chains (ELC and RLC) ^4–7^, converts torque generated by the motor into linear displacement.

**Fig 1.**
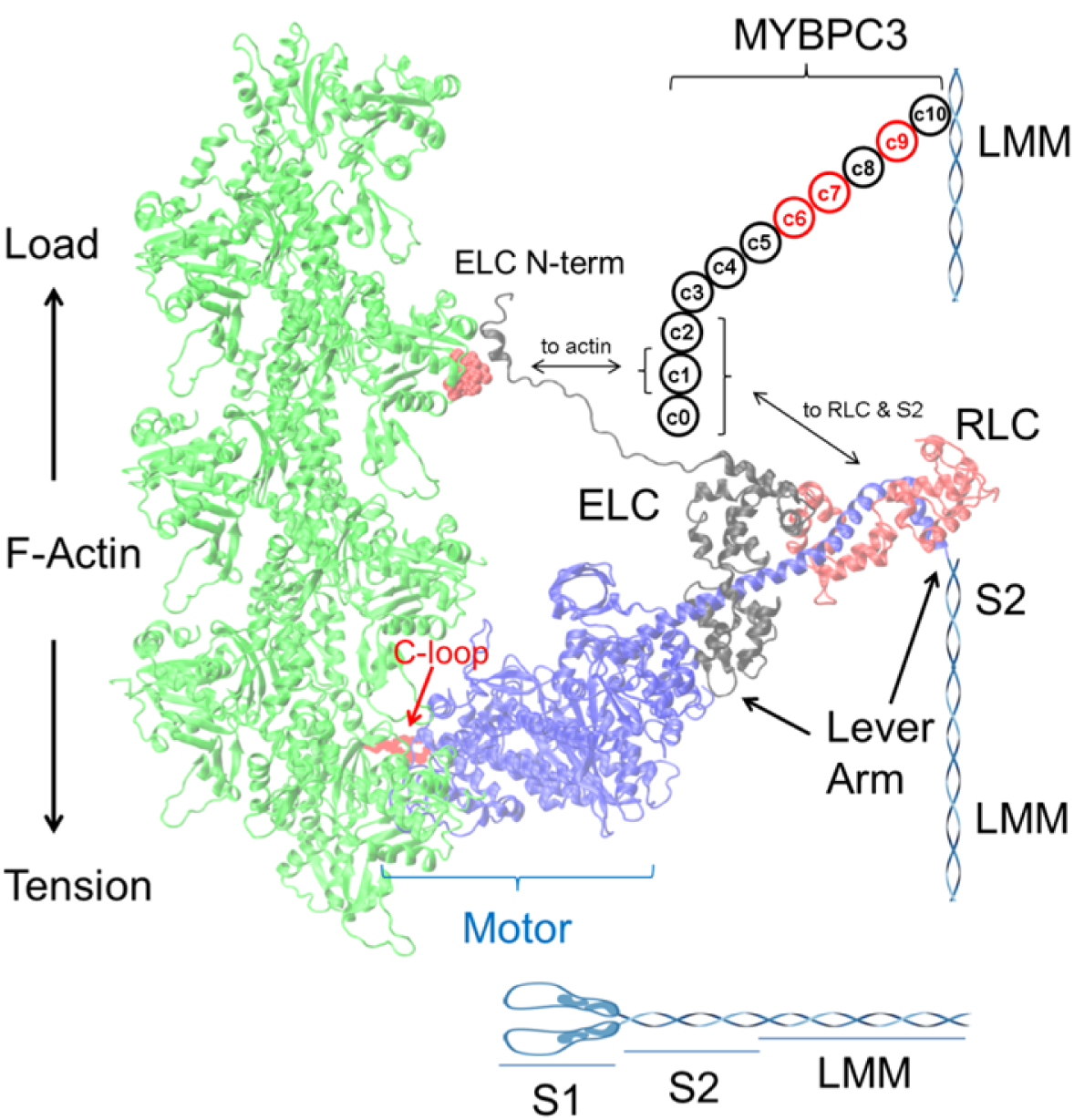
Myosin dimer proteolysis produces two subfragment 1 peptides (S1, blue), subfragment 2 (S2, blue), and light meromyosin (LMM, blue). S1 has a motor domain and lever arm with bound light chains ELC (black) and RLC (red). The motor binds to an actin filament and rotates the lever arm generating torque to apply tension on F-actin (green). The ELC N-terminus also binds actin to modulate myosin step-size. Mybpc3 has 11 domains with c10 binding myosin LMM and with c0-c2 maintaining transient interactions with actin, myosin S2, and RLC (mybpc3 domains in black or red). The actin binding site for mybpc3 (red space filling atoms) is proximal to the actin binding site for the ELC N-terminus.

Cardiac myosin binding protein C localizes to the C-zone of the muscle sarcomere and regulates the actomyosin sliding velocity ^8^. It has 11 immunoglobulin-like (Ig) or fibronectin-like (Fn) globular domains (c0-c10) resembling a pearl necklace with c0 near the N-terminus. The c0 Ig-like domain is unique to the cardiac isoform studied here ^9^. The protein contains several sites for serine, and one site for threonine, phosphorylation and involving phosphokinase A (PKA) and PKC in its regulation. The mybpc3 C-terminus associates with the myosin thick filament ^10^. The mybpc3 N-terminus associates with myosin S2 ^11^, RLC ^12^ and with F-actin in vitro ^13–15^ and in intact muscle ^16, 17^ (Fig 1). The role of mybpc3 in cardiac muscle regulation is extensively studied and associates with modulation of contractile force-velocity ^11, 18, 19^, myosin S2 stability ^11^, and myosin super-relaxation ^20^. As relates to modulation of force-velocity, we propose mybpc3 both resembles and competes with ELC because its transient actomyosin crosslink affects movement and it binds actin at the same site as the ELC N-terminus. We suggest it is a fourth subunit of the motor after myosin heavy chain (MHC), RLC, and ELC.

Actin has two major domains separated by a nucleotide-binding cleft that are further subdivided into subdomains 1 and 2, and, 3 and 4 ^21^. In their strong binding state, actin and cardiac myosin make multiple contacts over 3 actin monomers in the actin (thin) filament as show in Fig 1 ^22^. These contacts in actin include residues in all 4 subdomains ^23^. In myosin they included notable structured ^24, 25^ and unstructured surface loops ^26^ usually involving ionic interactions, hydrophobic regions on the myosin surface ^27^, and a unique contact between actin and the myosin ELC N-terminus ^28–30^. These contacts modulate actin activation of the myosin ATPase, actin affinity for myosin and, myosin step-size ^31^.

Human heart diseases link to variations in myosin, actin, and mybpc3 with diverse phenotypes reflecting modifications to cardiac muscle mechanochemistry. The depth and breadth of cardiac myosin ^32, 33^, actin ^1, 23^, and mybpc3 ^9, 17^ structural characterization is unique due to vigorous scientific interest driven by desires to positively affect human health and to understand a natural nanomotor design. Missense single nucleotide polymorphisms (SNPs) are clues for understanding cardiovascular disease mechanism quantitated here in an extensive database containing mutant location in a protein domain (domain), residue substitution (sidechain), human population group (demographic), and prevalence (frequency) as independent variables that imply the dependent disease phenotype and pathogenicity characteristics. The unknown disease model mechanism is surmised implicitly using a feed-forward neural network then interpreted explicitly with a discrete Bayes network for Neural/Bayes forecasting as already described ^34^. Involvement of the independent demographic and frequency characteristics is new to this application as is the application to actin. Neural/Bayes forecasting tests pathogenicity vs demographics of mutations in different protein domains. It provides a prognosis that assesses individual risk due to genetic background and gender, and, identifies protein domains and inter-protein interactions critical to disease mechanisms.

## 2. METHODS

### 2.1 SNP data retrieval

An automated search/fetch Perl script downloads the SNP reference numbers (rs#) from the National Center for Bioinformatics (NCBI) SNP database for the human cardiac myosin heavy and light chains, actin, and mybpc3 genes. Automated extraction of other information from the NCBI SNP database uses database text search/extract tools in Mathematica (Wolfram, Champaign, IL, USA) collecting location in the protein sequence, residue substitution, population demographic, allele frequency, and the clinical data set assigning pathogenicity and phenotype. Clinical phenotype and pathogenicity data is frequently unfulfilled (presumed unknown), inconclusive (presumed unknown), or contradictory among data submitters. We inspected the clinical data when results from data submitters conflicted and reached consensus by requiring two or more reports to agree. Otherwise, when there is no consensus, the result was presumed unknown.

Human ventricular βcardiac myosin (βmys) is encoded in the MYH7 (MHC), MYL3 (ELC), and MYL2 (RLC) genes. Human atrial αcardiac myosin (αmys) is encoded by the MYH6 (MHC), MYL4 (ELC), and MYL7 (RLC) genes. Human mybpc3 is encoded by the MYPBC3 gene. Human actin is encoded by the ACTC1 gene.

### 2.2 Network configuration

Trial network configurations associate mutant (mu) location in the protein domain (cd), residue substitution (re), population demographic (po), and SNP allele frequency (af) in a causal relationship with phenotype (ph) and pathogenicity (pa) in model configurations denoted in Fig 2.

**Fig 2.**
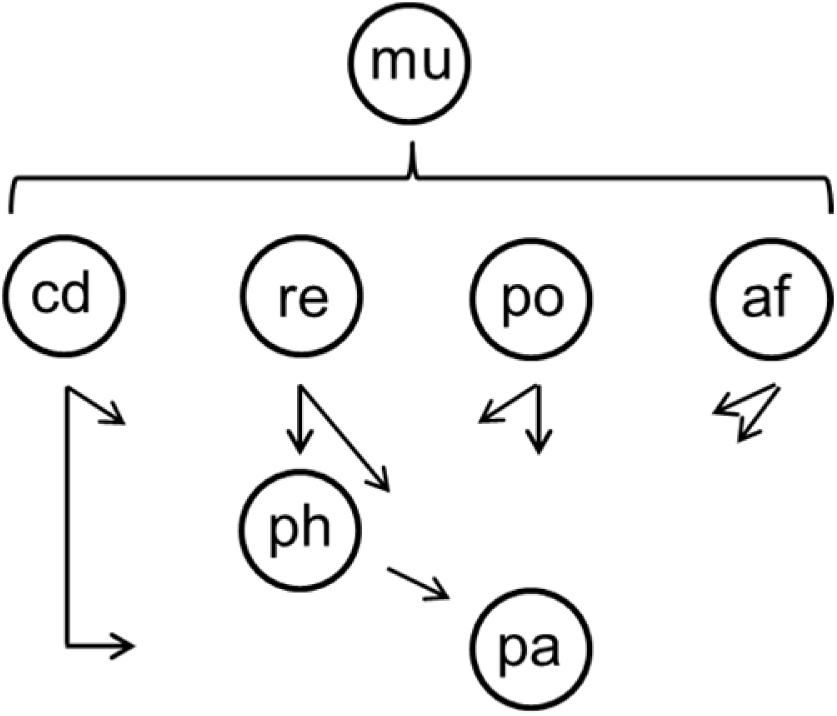
Configuration denoting relationships among mutant (mu) location (cd), residue substitution (re), population demographic (po), allele frequency (af), phenotype (ph), and pathogenicity (pa) that model the structure/function influence pathway in neural and Bayes networks.

All protein domains and their 2 letter abbreviations are indicated in Supplementary Information (**SI**) **Tables S1 and S2**. Myosin domains (cd) include 27 functional sites as described previously ^34^. βmys and αmys heavy chains have identical sequences except that αmys has single residue insertions at N211 and G634 (αmys numbering) that slightly alter the domain assignments. The myosin light chain domain designations are unaffected by sequence differences between βmys and αmys. Mybpc3 and cardiac actin domains include 26 and 28 sites. Every SNP in the database has an assigned domain. Fig 3 shows linear representations of myosin, actin, and mybpc3 indicating mutual binding sites and the locations of domains listed in **SI Tables S1 and S2**.

**Fig 3.**
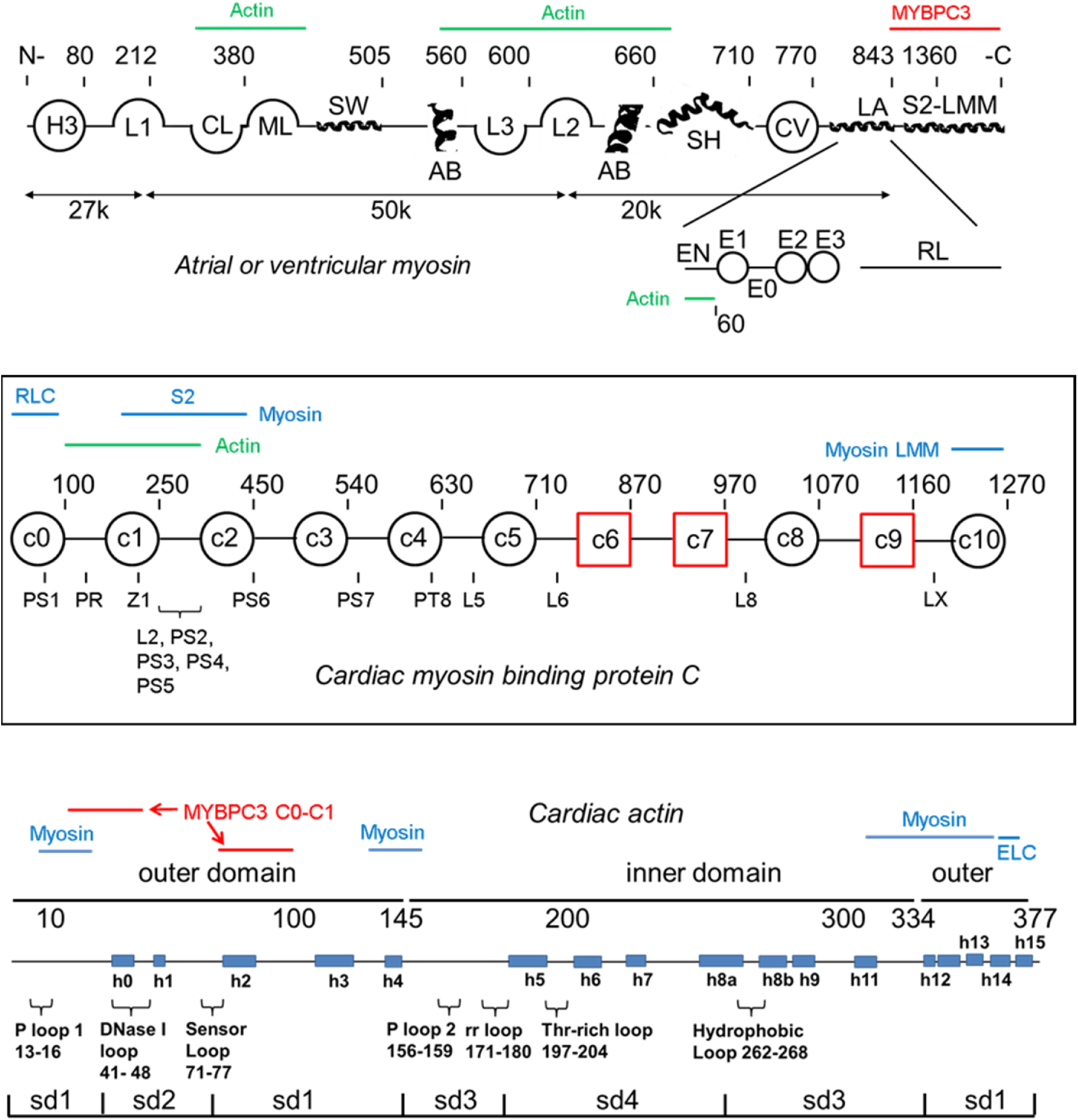
Linearized diagrams for cardiac myosins (top), mybpc3 (middle), and actin (bottom) identifying most domains defined in SI Tables S1 and S2. The myosin diagram does not indicate the active site (ac), OM binding site (om), and mesa (me) because they occupy multiple regions in the linearized representation. Myosin light chains appear below the heavy chain. Actin and MYBPC3 binding sites are indicated in green and red above the heavy chain and below the ELC. The mybpc3 diagram consists of 8 Ig-like domains (black circles) and 3 fibronectin-like domains (red squares). Phosphorylation sites (PS or PT) are indicated below the chain. Domain linkers of interest include the proline rich linker (PR) and L2 containing a regulatory site. Z1 is a zinc binding site. Myosin S2 and LMM and actin binding sites on mybpc3 are indicated above the linearized model. The cardiac actin diagram consists of α-helical segments (h0-h15) and looping regions indicated with brackets 1. Domain structure divides the molecule into outer and inner domains and four subdomains (sd1-4). Myosin and mybpc3 binding sites on actin are indicated above the linearized model in blue and red.

Residue substitution (re in Fig 2) refers to the reference and substituted residue (ref/sub) pair. Possible ref/sub combinations have 420 possibilities for 21 amino acids. A hydrophobicity index was derived from model peptide studies on the stability of amphipathic α-helices. It ranks residues as hydrophilic (p), neutral (n), hydrophobic (m), and very hydrophobic (h) ^35^. The descriptive index simplifies residue substitution to a more tractable 16 ref/sub pairs characterizing every mutation. Input residue substitution pairs and their 2 letter abbreviations indicated in **SI Table S3** summarizes input indicated by *re* in the Fig. 2 model. In addition, each residue is assigned an integer score whose difference for a ref/sub pair (the ref/sub score Δ) ranges from −3 for hydrophilic/very-hydrophobic pairs to +3 for very-hydrophobic/hydrophilic pairs. The ref/sub scores are also indicated in **SI Table S3**.

Residue substitution prevalence (allele frequency or af in Fig 2) in the human population group (demographic or po in Fig 2) fill out the independent parameters in the network. Demographic groups (po) and their 3 letter abbreviations are indicated in **SI Table S4**. We will use subsets of these demographic groups pertaining to ethnic identity or gender as indicated in **Table S4** to draw attention to interesting data tendencies. Allele frequency is a continuous variable in the database on the interval 0 ≤ af ≤ 1 for 1 meaning all alleles are substituted by the SNP. These data are subdivided into the three discrete categories indicated in **SI Table S5**.

Phenotype (ph) and pathology (pa) data have standardized classifications for cardiovascular disease. A total of 13 phenotypes for cardiac myosin, mybpc3, and cardiac actin from the NCBI SNP database include hypertrophic cardiomyopathy (hc), dilated cardiomyopathy (dc), restrictive cardiomyopathy (rc), left ventricle noncompaction cardiomyopathy (lv), cardiomyopathy (cm), congenital myopathy (gm), atrial fibrillation (af), ventricular fibrillation (vf), ventricular tachycardia (vt), cardiovascular phenotype (cp), atrial septal defect (ad), native (nv), and unknown (uk). Cardiomyopathy (cm) describes apparent heart conditions in which specific etiologies are not clearly identified. Cardiovascular phenotype (cp) describes conditions affecting the cardiovascular system but not directly the myocardium (e.g. valvulopaties, aortic coarctation) ^36, 37^. Congenital myopathy (gm) are muscular diseases of genetic etiology that rarely affects the heart. Left ventricle noncompaction (lv) and atrial septal defect (ad) are heart developmental defects sometimes leading to secondary cardiomyopathy due to hemodynamic abnormalities. This phenotype list is also indicated in **SI Table S6**.

Pathogenicity (pa) is likewise taken from the NCBI SNP database and includes pathogenic (pt), likely pathogenic (lp), benign (be), likely benign (lb), and unknown (uk). The pathogenicities and their 2 letter codes are summarized in Table 1.

**Table 1.** Pathogenicity (pa) with 2 letter codes

| pathogenicity | code |
| --- | --- |
| (p)a(t)hogenic | pt |
| (l)ikely (p)athogenic | lp |
| (be)nign | be |
| (l)ikely (b)enign | lb |
| (u)n(k)nown | uk |

### 2.2 SNP neural network

The neural network pathogenicity code modeling structure/function influences from disease follow from the model in Fig 2. They relate domain position (cd), residue substitution (re), population (po), and allele frequency (af) inputs to phenotype and pathogenicity outputs through 4 fully connected linear (hidden) and dropout layers with 304 or 252 nodes each, and, a softmax layer conditioning output for digital classification of phenotype and pathogenicity as indicated in Fig 4. Dropout layers mitigate overtraining. Training data contains 50% of the fulfilled 6ddps.

**Fig 4.**
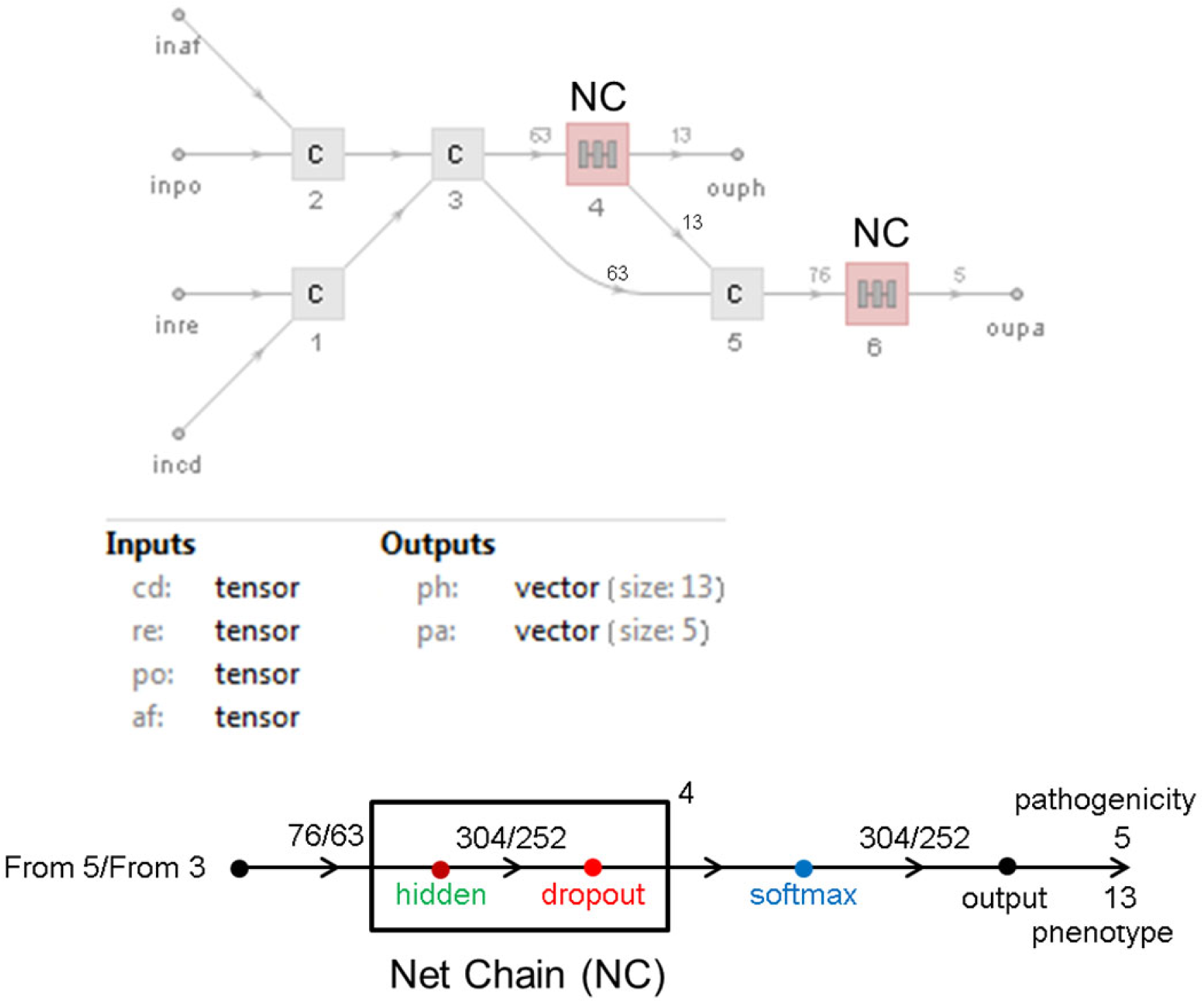
Feed forward neural nets relating inputs for the site of the modification (cd), residue substitutions (re), population (po), and allele frequency (af) with disease phenotype (ph) and pathogenicity (pa) in the models corresponding to those in Fig 2. The Net Chain (NC), depicted in the lower half of the figure, is a component in the model in the upper half. Numbers above the horizontal line are nodes. Four connected hidden layers are indicated by the superscripted 4.

Trial training data sets of fulfilled 6ddps are selected randomly from the validation data set but subject to the constraint that each phenotype and pathogenicity outcome must be represented in the set except when their representation in the fulfilled 6ddp’s is <2 occurrences. Learnable weights for the linear layers are randomly initialized. Weight initializations are normally distributed with zero mean and standard deviation of (1/n)^½^ (for n inputs). Bias is initialized to zero. A total of 1800 trials generate the 20 best implicit models. This process is repeated 5 times and the results (100 total models) are combined into 24-25 best-of-the-best implicit models. These 24-25 neural networks embody distinct implicit models for disease that are sufficiently diverse to cause normally distributed estimates for Bayes network probabilities (discussed below) implying that they randomly sample the set of good implicit disease models.

### 2.3 Neural network validation

The ability to correctly classify new SNPs (new-unknown data corresponding to new unfulfilled 6ddps) measures suitability of a neural network trial. We set aside fulfilled 6ddps to be the new-unknown dataset but from the part of the validation data pool that does not include training data since a real new-unknown could never be a part of training. Each new-unknown 6ddp has its position and substitution assignment evaluated by the neural network trial with the output phenotype and pathogenicity compared to the know value. This comparison is the new-unknown predictor metric that we use to rank neural network model suitability.

The best model neural networks (ranked by their new-unknown predictor metric) predict unfulfilled (ph, pa) outputs from their domain position (cd), residue substitution (re), population (po), and allele frequency (af) assignments. Fulfilled and predicted outputs combined are the database for Bayes network (Fig 2) tasked with formulating a statistics based myosin, mybpc3, or actin structure/function mechanism as described in the next section.

### 2.4 Bayes network modeling of myosin structure/function

Fig 2 shows the Bayes network model. Arrows imply a direction for influence hence the domain (cd), residue substitution (re), population (po), and allele frequency (af) assignment implies a probability for phenotype (ph) and pathogenicity (pa). Datasets 6ddpMYH7.xls, 6ddpMYH6.xls, 6ddpMYBPC3.xls, and 6ddpACTC1.xls in SI show the fulfilled and unfulfilled 6ddps for βmys, αmys, mybpc3, and actin containing 4523, 6649, 4003, and 170 variations in the database corresponding to 1877, 1798, 1181, and 131 distinct residue substitutions. Combined fulfilled and predicted 6ddp data sets are expressed as conditional probability tables (CPT’s, eq. 1) defining the joint probability density (left hand side of eq. 1) representing the networks in Fig 2 such that,

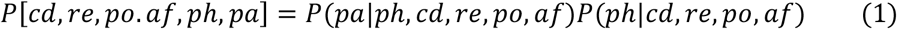

We use this statistical method to query pathogenicity (pa) or phenotype (ph) probability due to protein residue domain (cd), substitution (re), population (po), and allele frequency (af).

### 2.5 Significance testing

We tested reproducibility and significance of the cardiac disease models by generating a large pool of implicit neural network models in the Fig 2 configuration. Models are independent solutions to this highly constrained problem that are ranked for reliability by the new-unknown predictor metric. We used the ranked model solutions to estimate the collective quantities described subsequently in RESULTS related to demographics for each protein sequence. Best ranked model solutions formed a finite subset for each protein drawn from the larger pool of independent solutions. Model solution members in each subset were increased (best solutions by the new-unknown predictor metric used first) until the collective quantities were unchanged by further enlargement of the subset. This selection process favored model subsets best approximating the real disease mechanism by minimizing random error but is unlikely to address systematic model limitations. Each model solution in a subset exactly reproduces 80-92% of the known 6ddps in the target protein constraining potential systematic errors in the models to just 8-20%.of the dataset and implying the measure of their reliability, however, potential systematic errors will not affect relationships indicated within the target protein model solutions described in RESULTS.

One-way ANOVA with Bonferroni or Tukey-Kramer post-tests for the p< 0.01 or p<0.05 significance levels are used for all significance testing when a significance level (p) is mentioned.

## 3. RESULTS

We applied the Neural/Bayes network models for disease in Fig 2 to ventricular and atrial cardiac myosin, mybpc3, and cardiac actin to investigate inheritable cardiac disease demographics versus the mutant location in protein functional domains. Given human populations listed in **SI Table S4**, pathogenic or benign outcome probabilities were calculated for nonsynonymous SNPs falling into the functional protein domains represented schematically in Fig 3 and listed in **SI Tables S1-S2**. Results are from using the 24-25 independent best-of-the-best implicit models for each protein’s contribution to the contractile mechanism. Models selected satisfy the reproducibility and significance testing described in METHODS (section 2.5).

Pathogenicity is summarized as either pathogenic or benign by combining likely-pathogenic with pathogenic probabilities or likely-benign with benign probabilities (see **Table 1**). Demographic probabilities for each protein functional domain and pathogenicity category were computed with Bayesian statistics as described ^34^ and listed in **SI Tables S7-S10**. An example from that data for a single functional domain in βmys is shown in Fig 5. There are 26-28 domains for the proteins considered suggesting that a baseline contribution to pathogenicity probability for each functional domain is ∼3% of the total. We split this probability between pathogenic and benign categories suggesting domains contributing significantly and specifically to function will likely contribute >1% to each category. We refer to these domains as qualified functional domains (QFDs). SNPs in QFDs cause pathogenic and benign outcomes by selective residue substitutions suggesting their side chains are specifically involved in sarcomere function and implying that their demographic trends are reliable.

**Fig 5.**
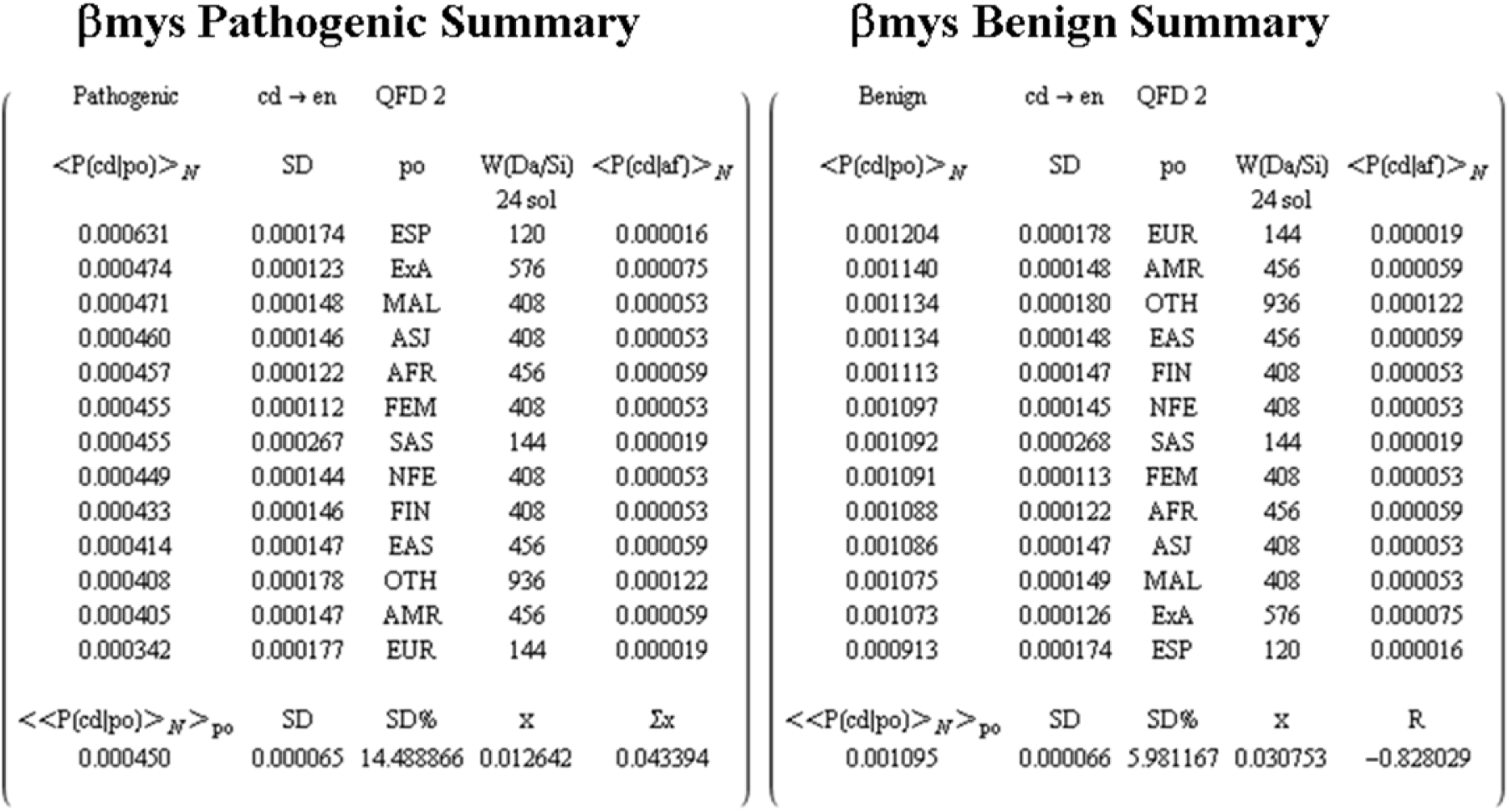
Demographic probabilities and statistics for populations (po) in columns ranking most to least likely from top to bottom. This example for the ELC N-terminus domain (en) from βmys is taken from SI Table S7. *En* domain is a QFD in βmys. Quantities are defined in the text.

Protein domains included in a result for a given protein are those represented in each of the 24-25 implicit models and for both pathogenic and benign outcomes. The example matrices in Fig 5 are for the ELC N-terminus domain (en). It is a QFD and drawn verbatim from the complete results in **SI Table S7**. In Fig 5 and **SI Table S7**, large side-by-side matrices are for pathogenic (left) or benign (right) pathogenicity while both matrices pertain to the same functional domain (cd, coded in the first row). Probabilities (first column) are an average over 24-25 best-of-the-best implicit models (under heading <P(cd|po)>*_N_*, where *N* is 24-25) for population demographic (po) in the third column. SD (standard deviation, second column) measures the spread. The number of 6ddps contributing to the probability, indicated in column 4, combine SNP data from NCBI when known (Da for data) with that from estimates by the neural network when unknown (Si for simulated) from 24 implicit models (sol). Mean allele frequency from the contributing 6ddps (under heading < P(cd|af) >*_N_*), appears in column 5. N-averaged probabilities in column 1 are themselves averaged over the populations to obtain the value in the bottom row under the heading << P(cd|po) >*_N_* >_po_ and with spread below the lower SD in column 2. The quantity indicated by *x* is the total protein domain probability for pathogenic (left matrix) and benign (right matrix) outcomes hence they are comparable horizontally (over pathogenicity) for a given protein domain and vertically among the different protein domains. A domain is a QFD (see Methods section 2.5) when x ≥ 0.01 for both pathogenic and benign categories (*en* is a QFD in Fig 5). All *x* quantities for a given protein (seen in **SI TablesS7-S10**) do not sum to 1 because *x* from some domains are not represented in both pathogenic and benign categories and are not included in the tables. The *Σx* appearing only on the pathogenic side indicates the sum of *x* for pathogenic and benign cases in the same domain (i.e., side-by-side matrices). Pearson’s *r* (R) spans the dynamic range −1 ≤ R ≤ 1 and measures linear correlation between the integer sequence counting three or more populations in column 3 and declining probabilities in column 1. It is assigned the largest magnitude value for pathogenic and benign cases in each protein domain and appears only on the benign side. It relates correlation strength for probability vs demographics over the protein functional domains. The R-correlation range over functional domains in βmys, αmys, mybpc3, or actc1 is listed in the last row of **SI Tables S7-S10**.

The detailed results in **Tables S7-S10** are broadly summarized by significance tested estimates for most and least overall pathogenic or most and least overall benign ethnic population or gender in Fig 6. The *summary ranking* estimates identify the demographic that consistently ranks highest or lowest in pathogenic outcomes over the entire protein as measured directly by pathogenicity or indirectly by the lowest or highest ranking in benign outcomes. Summary ranking is a qualitative measure for heart disease burden in a demographic and is done in two ways. The first using Bayes statistics that is appropriately weighted by normalized probabilities and a second simpler approach that permits easier comparison with quantitative results in **SI Tables S7-S10**. The second method does not preserve relative weighting of functional domains but requires that included functional domains have ≥ 1% of the total probability for pathogenic or benign outcomes. This is different from the QFD rule where both pathogenic and benign outcomes must qualify with ≥ 1% of the total probability. In the second method ranking 1^st^ or 2^nd^ at the high end of the demographic probability listing (column 1 in the qualified matrices from **SI Tables S7-S10**) earns a 3 or 1 score while ranking lowest or second from lowest in the probability listing earns a −3 or −1 score. These integers are replaced by real probabilities, but keeping the sign assignment, for the Bayes statistics method. Both methods give identical ranking of data used here. When gender is the population group, 2 quantities are involved and scores rank first or last positions with 1 or −1 or the appropriately signed probability for the Bayes statistics method. Smaller matrices in Fig 6 show summary rankings and numerical scores for ethnic population or gender subsets over protein domains that contribute >1% probability to pathogenic (left) or benign (right) outcomes. Ethnic (eg., AFR or EUR) and gender subsets are defined in **SI Table S4**.

**Fig 6.**
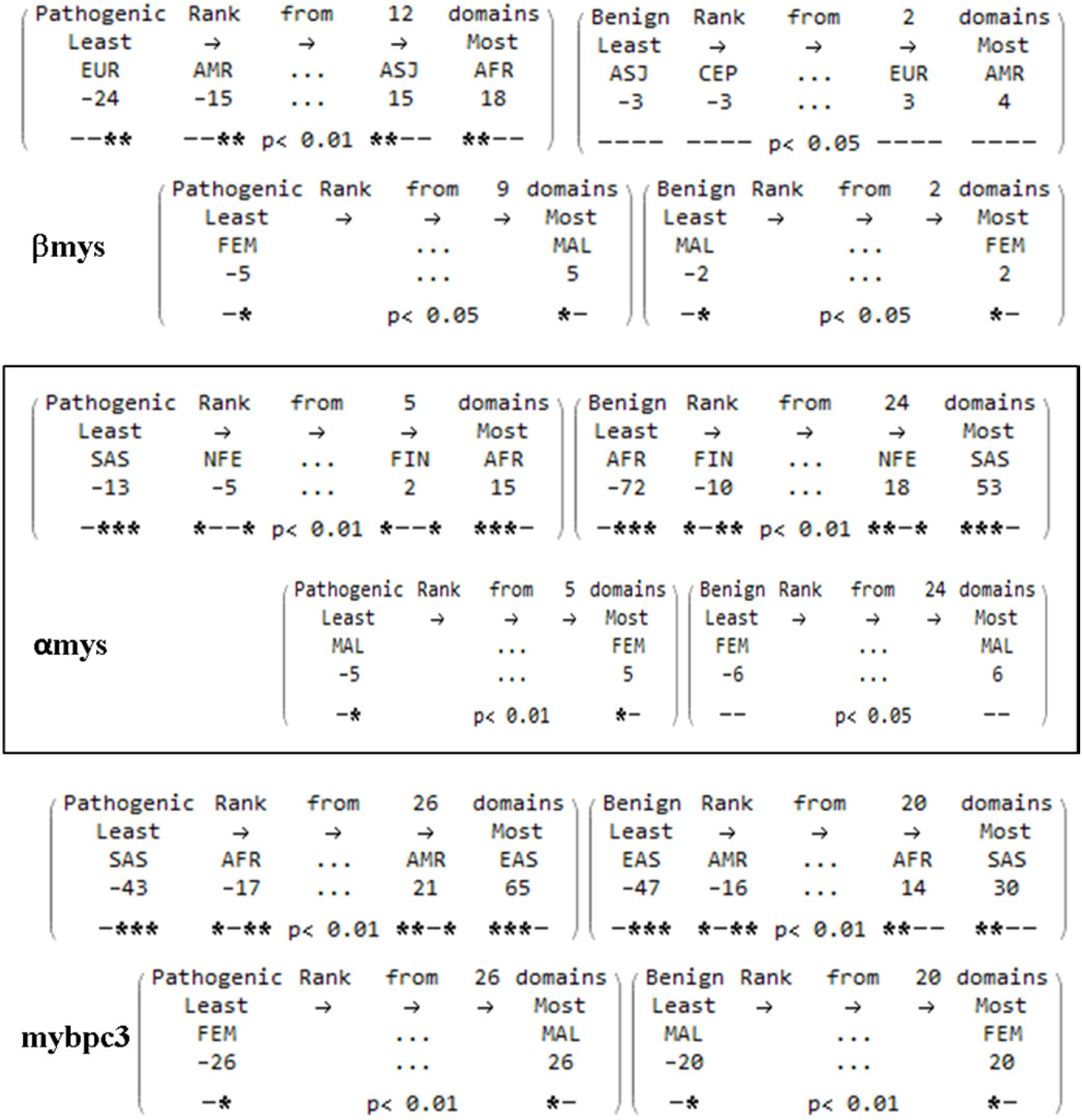
Summary ranking matrices for βmys, αmys, and mypbc3 compare ethnic population (upper row of matrices) or gender (lower row of matrices). The bottom row in each matrix indicates significance, p, at the 0.01 or 0.05 levels. Symbols – or * imply insignificant (–) or significant (*) differences between the current column population or gender and other columns in the matrix represented by the 4 (top matrices) or 2 (bottom matrices) column positions. The current column demographic is always insignificantly different from itself. Data from cardiac actin (actc1) is not shown because it did not give a significant summary ranking list.

### 3a. Ventricular cardiac myosin (βmys)

Disease demographics for βmys SNPs correlate pathogenicity probability with population. An example for one of the QFDs is shown in Fig 5. Pathogenicity probability vs population is negatively correlated and approximately linear for all qualifying domains as measured by quantity *R* (**SI Table S7**). Summary rankings in Fig 6 for βmys has African (AFR) and Ashkenazi Jewish (ASJ) ethnic populations scoring similarly at the high end of the probability scale, to develop pathologic cardiac disease, for the myosin domains represented in each of the 24 best-of-the-best implicit models. European (EUR) and Ad Mixed American (AMR) ethnic populations are more protected from serious disease by scoring at the low end of the probability scale for pathologic outcomes. The 2 most pathogenic populations (AFR and ASJ) are significantly different from the two least pathogenic populations (EUR and AMR) for ANOVA significance testing with p< 0.01. Comparing gender (Fig 6), males (MAL) are more likely to develop pathogenic heart disease than females (FEM) for ANOVA significance testing with p< 0.05. βmys is predominant in the ventricle implying a male preference for ventricle disease. On the benign side of the probability tables, results are less definitive with most probability density for benign outcomes spread over domains inconsistently represented in the 24 best-of-the best implicit models (except for the QFDs, see next paragraph). Allele frequency is low for every population. These prognosis indicators summarized above for βmys add personalized depth to a health plan addressing possible inheritable disease. Trends for protein domains in pathogenic or benign categories are immediately actionable for prognosis.

Two domains dominantly impacting disease demographics for βmys are QFDs with >1% of the total probability for both pathogenic and benign outcomes. They are the actin binding sites: C-loop (cl) and ELC N-terminus (en) identified in Fig 1. C-loop is a conserved structured loop on the surface of βmys that senses ATP binding, weak actin binding, and actin-activation of myosin ATPase ^24^.The C-loop participates in formation of the rigor bond with actin ^25^. The ELC N-terminus modulates strain-dependent mechanics ^38^. It is the site for one of two regulatory mechanisms (the other one is strain-dependent ADP release) that remixes the 3 different myosin unitary step-sizes with changed stepping frequencies in response to loading ^30^. On board machine intelligence in βmys down-shifts average displacement with increasing loads by utilizing these strain-dependent regulatory mechanisms ^31, 39^. The C-loop and ELC N-terminus QFDs have a site selective and specific response to residue substitutions in the peptide chain that are involved in cardiac disease. They are differently sensitive to ethnic population implying deeper (than the βmys sequence) genetic and cultural factors play a role that we capture in the neural net implicit models used here to classify a large dataset and to address population dependence of disease prognosis. New research will address the deeper genetic and cultural factors in play.

### 3b. Atrial cardiac myosin (αmys)

Disease demographics, summarized in **SI Fig S8** for αmys SNPs, correlates pathogenicity probability with population from the 25 best-of-the-best implicit models. Pathogenicity probability vs population for αmys is negatively correlated and approximately linear for all qualifying domains as measured by quantity *R* (**SI Table S8**). *R* for αmys occupies a higher amplitude and narrower range than for βmys implying a more uniform and definitive statement from the best implicit models. Pathogenic and benign outcome statistics for αmys vs βmys are qualitatively reversed. The benign outcomes produce the more definitive findings for αmys and where most probability density for pathogenic outcomes spread over domains inconsistently represented in the 25 best-of-the-best implicit models. The summary rankings in Fig 6 for αmys has South Asian (SAS) and Non-Finnish European (NFE) populations scored at the higher end of the probability scale for benign cardiac disease for the myosin domains represented in each of the 25 best-of-the-best implicit models. African (AFR) and Finnish in Finland (FIN) populations score at the low end of the probability scale for benign outcomes, i.e., they are less protected from serious disease. Populations in the benign disease summary are significantly different from each other for ANOVA significance testing with p < 0.01. Female (FEM) populations score higher for pathogenic outcomes compared to male (MAL) populations with the difference significant at the p < 0.01 level. The latter MAL/FEM ordering is also reflected in the probability scores of the benign outcomes. αmys is predominant in the atrium implying a female preference for atrium disease. Gender inequality for α- and βmys suggests that atrium centric inheritable cardiac disease is more detrimental to women’s health while ventriculum centric inheritable cardiac disease is more detrimental to men’s health. The results again suggest deep genetic and cultural factors play a role in the cardiac disease that we capture in the neural net implicit models used here to classify a large dataset and to address population dependence of disease prognosis.

**SI Table S8** for αmys indicates that five QFDs impact disease demographics for αmys. They are the ELC N-terminus (en, Fig 1), lever arm (la, Fig 1), ELC binding IQ domain on the lever arm (qe), and k7 and k5 that are large (200 and 400 residue) default domains corresponding to 27k and 50k molecular weight fragments produced by proteolysis at unstructured loops 1 and 2 (l1 and l2). The ELC N-terminus is the single QFD that presents in both ventriculum and atrium centric inheritable diseases and is the largest contributor to probability in both α- and βmys. ELC is intimately involved in myosin strain-dependent mechanics ^38^, its binding to the IQ domain (qe) stabilizes the lever arm ^40^, and it is critical for the native folding of the myosin heavy chain after translation ^41^. The lever arm (la) converts torque generated in the motor domain of myosin into linear displacement of actin. It is the ultimate determinant of the myosin step-size^42^. The k7 and k5 domains qualify as QFDs due to the size of their contribution to pathogenicity probabilities but are heterogeneous with respect to function as their sequences flank SH3 (h3) and the active site (ac) in k7, and, several of the actin binding sites in k5. The k7 and k5 are default domains that are assigned SNPs whose location falls outside the sequence range of other more specifically functionally identified domains. Identification of k7 and k5 as QFDs suggest that sequence assignment to functional domains needs further scrutiny possibly to involve a larger part of the protein sequence in SH3, active site, and actin binding domain in the heavy chain.

### 3c. Cardiac myosin binding protein C (mybpc3)

Disease demographics are summarized in **SI Fig S9**. Mybpc3 domains contributing substantially to the pathogenicity probabilities far outnumber those in βmys and αmys implying the protein and its human host readily tolerates these variations that measurably impact function. Pathogenicity probability vs population for mybpc3 is negatively correlated and approximately linear for all qualifying domains as measured by quantity *R* (**SI Table S9**). The summary rankings in Fig 6 for mybpc3 has East Asian (EAS) and Ad Mixed American (AMR) populations are most likely, while South Asian (SAS) and African (AFR) populations least likely to suffer pathologic cardiac disease for the mybpc3 domains represented in each of the 25 best-of-the-best implicit models. The benign pathogenicity outcomes (right matrix **SI Fig S9**) trend identically by favoring South Asian (SAS) and African (AFR) while disfavoring East Asian (EAS) and Ad Mixed American (AMR) populations. Prognosis is significantly impacted by gender. Female (FEM) populations always fare better than Male (MAL) counterparts for pathogenic outcomes. Summary distinctions between ethnic populations and genders are significant at the p< 0.01 level.

Four QFDs (c0, c1, l2 and c10) for mybpc3 (out of 20) are especially notable (see **SI Fig S9**). The c0-c1-l2-c2 Ig-like and linker domains at the mybpc3 N-terminus transiently bind myosin ^12^, while c1 and l2 (l2 also called M-domain containing 4 phosphorylation sites), transiently bind actin ^13–17^ (see Fig 1). The c10 (cx) Ig-like domain on the mybpc3 C-terminus binds to LMM ^17^ anchoring mybpc3 to the thick filament (**Figs 1 & 3**). The binding sites facilitate actomyosin translation velocity modulation in a mechanism regulated by mybpc3 phosphorylation in the l2 linker ^18, 43–45^. Transient mybpc3 N-terminus/actin binding imitates transient ELC N-terminus/actin binding ^46^ by targeting the same site on the actin surface (Fig 1) implying they compete for it within the C-zone. Actin binding of the ELC N-terminus performs a strain dependent down-shifting of myosin based displacement by altering the relative frequency of the three βmys step-sizes ^31, 47^. The specific effect of mybpc3 actin binding and competition with ELC on myosin step-size in the C-zone is unknown. Total probability for SNPs to contribute to pathogenicity is larger for *c1* than any other QFD in mybpc3 attesting to its central significance in the implicit disease mechanisms coded in the best-of-the-best neural network models.

### 3d. Cardiac actin (actc1)

Disease demographics for actin summarized in **SI Fig S10** are the simplest of the four cardiac sarcomeric proteins studied. The statistics do not identify any QFDs because there is little data on the actin SNPs, possibly because the human protein has few natural SNPs that are not lethal to the fetus. They identify just one domain, the ring-rich loop (rr), with a very substantial contribution to pathogenicity. Pathogenicity probability vs population for cardiac actin is negatively correlated and approximately linear for all qualifying domains as measured by *R* (**SI Fig S10**). No summary ranking of demographics equivalent to those in Fig 6 is possible in this case. The ring-rich loop is not implicated in F-actin intermolecular interactions possibly allowing individuals with SNPs in that domain to survive longer although the associated cardiac disease is pathological. Actin sequences are highly conserved between skeletal (acta1) and cardiac (actc1) isoforms implying that additional data relevant to cardiac function might be gleaned from SNPs in the skeletal actin although notable structural ^48^ and functional characteristics ^31^ differentiate them possibly complicating disease models.

## 4. DISCUSSION

Myosin is the engine powering the beating heart. Its motor domain transducer located within the heavy chain contains ATP and actin binding sites, and, mechanical elements coupling motor generated torque to the myosin filament backbone for transduction/mechanical coupling. The mechanical coupler is an α-helical lever arm, stabilized by essential and regulatory light chains (ELC and RLC), that rotates to impel strongly bound actin filaments (Fig 1). Linear actin displacement from unitary lever arm rotation produces a unitary displacement (step-size) that responds to conditions in real time by using a second cyclical interaction between actin and the ELC N-terminus ^22, 29^ to modulate step-size length ^47^. Myosin in the contraction cycle adapts to changing power demands by regulating contractile force and velocity using 3 distinctive unitary step-sizes with step-size choice decided mainly by load ^47^. Down-shifting average step-size changes myosin from a high-displacement transducer for high velocity auxotonic shortening into a low-displacement transducer maintaining tension in near-isometric contraction ^39^. Native myosin functionality could require the structured environment and proximity to ancillary protein components in the muscle sarcomere such as mypbc3, or, might fully replicate its native behavior in vitro as a purified, isolated, and independent motor translating actin. Whether force-velocity regulation is a systemic property of the sarcomere or an intrinsic property of an autonomous myosin impacts approached to researching disease mechanisms.

### Sarcomere protein integration explains demographic/SNP-pathogenicity correlation

*In vitro* single myosin mechanical characterization uses purified and isolated myosin and reveals a telling correspondence between *in vitro* and *in vivo* systems that indicates myosin is to some extent an autonomous molecule such that the cardiac myosin is functionally the muscle in a molecule ^47^. Autonomous myosin codes its mechanism for real time force-velocity regulation into the protein sequence that was captured in a Neural/Bayes network model ^34^. Single residue sequence variation from a SNP in myosin or mybpc3 is a common cause of inheritable heart disease that affects people worldwide. Our earlier work developed a machine intelligence model for disease that implicitly characterized SNP impacts on function providing a predictive Neural/Bayes network model for SNP variation disease pathogenicity. Predictability contingent on an autonomous motor implies that a SNP in the motor has implications independent of demographics providing the protein sequence is otherwise conserved. Moreover, structure/function studies of the motor always assume myosin motor autonomy. Now we involved human demographics in SNP classification and find pathogenicity correlates with human subpopulations and gender. Our realization that mybpc3 forms a third actomyosin interaction competing with the ELC N-terminus ratchet ^30^ implies a new strain-dependent mechanism outside the myosin molecule modulating motor adaptation to load. The new predictive Neural/Bayes network model for myosin, mybpc3, and actin variation disease pathogenicity, developed here from *in vivo* human data involving the population genetic/cultural background and gender, promises a more realistic statistical prognosis. This working model, and the integral myosin/mybpc3 motor concept, implies some of the wider considerations involved in understanding heart disease as a systemic maladaptation.

Earlier work introduced a systemic heart disease mechanism to explain how widely spatially distributed point mutations in myosin, mybpc3, or actin cause specific and unique motor functional alterations but induce a common phenotype such as hypertrophic cardiomyopathy ^49^. Single soleus muscle fibers carrying βmys mutations in the converter domain had substantial and significantly higher contractile variability compared to normal control muscle fibers due to variation in mutant allele frequency (af) among individual cells in the tissue ^50^. In the heart it was proposed that contractile force imbalance due to unequal fractions of mutated and wildtype protein among individual cardiomyocytes over time induces cardiac remodeling and hypertrophic cardiomyopathy.

The M-band contains connective elements in the sarcomere important for managing force imbalances during active muscle contraction. It has a role as shock absorber in contracting muscle dealing with dynamic mechanical stress by changing its protein composition in response to changing demands ^51^. It is an adaptive substructure of the sarcomere responding to disease related altered myosin, mybpc3, or actin function that is itself probably impacted by demographic variation. The M-band is a potential addition to the integral myosin/mybpc3 motor concept comprising the actual autonomous contractile system.

### Alternative role for mybpc3

Mybpc3 is the third actomyosin crosslinker after the myosin heavy chain and ELC in striated muscle and is now proposed to impact motor strain-dependent mechanics. Its multi-component structure for actomyosin connectivity has the C-terminus Ig-like domain (cx) anchored to the myosin thick filament and the c0-c1-l2-c2 N-terminus domains (l2 is a linker including 4 phosphorylation sites involved in regulation sometimes called the M-domain) engaged in transient binding to myosin (Fig 1). Each subdomain element in the connectivity mechanism, except c2, is a QFD by analysis of the relationship of their SNPs to disease pathogenicity independently suggesting they are critical to function. The atrial and ventricle motor proteins (α- and βmys) both identify the ELC N-terminus as a QFD while αmys also identifies its complementary Iq-domain for ELC binding as a QFD implying connecting sites from myosin heavy chain-to-ELC-to-actin are necessary for native functionality. An analogous situation occurs in mybpc3 with the *cx* anchor to myosin (equivalent to the ELC site that binds the lever arm Iq-domain) and transient actin binding activity site c0-c1-l2-c2 (equivalent to the ELC N-terminus in myosin) qualifying as QFDs (except for c2) forming the connectinons necessary for strain regulation. The N-terminus of ELC and the N-terminus of mybpc3 appear to compete for the binding site on actin (Fig 1) implying they competitively influence myosin strain-dependent mechanics. Mybpc3 could sterically disrupt the normal actin/ELC interaction in the C-zone or involve itself as an integral part of an independent myosin strain-dependent mechanism. New experimental work investigating these possibilities will address the impact of the mybpc3 on loaded myosin contractility in the context of single myosin mechanics as done previously for the ELC N-terminus/actin interaction ^30, 47^.

### Demographic inequalities in heart disease

Implicit neural network modeling of human disease mechanisms from nonsynonymous SNPs located in cardiac myosins (α- and βmys) and mybpc3 finds that demographics significantly influence heart disease pathology. Similar modeling for cardiac actin was inconclusive probably because of insufficient SNP data. Considering just myosin and mybpc3 findings, SNPs in these proteins collectively favor pathogenic outcomes in the African (AFR) or East Asian (EAS) populations while favoring benign outcomes in the South Asian (SAS) and Ad Mixed American (AMR) populations. These findings are a basis for assessing an individual’s risk for a particular sequence variant and population group. Statistically significant gender bias in mybpc3 has males (MAL) developing more pathological disease from closely related SNPs when compared to females (FEM). Statistically significant gender bias in β- and αmys suggests pathological and benign outcomes from closely related SNPs in these proteins correlate with pathological outcomes favoring male ventriculum and female atrium and with benign outcomes favoring male atrium and female ventriculum. This observation has value when evaluating risk from heart contractility metrics since it shows atrial or ventricular functional impairment has different implications for women or men.

## 5. CONCLUSION

SNPs cause unique residue substitutions in the functional domains of ventricular myosin (βmys), atrial myosin (αmys), mybpc3, and actin (actc1). Independent SNP characteristics of domain location, residue substitution, demographic, and allele frequency predict their dependent phenotype and pathogenicity using a feed-forward neural network model (Fig 2). The NCBI SNP database was mined to assign known independent and dependent discrete variables in 6 dimensional data points (fulfilled 6ddps) for each protein. The latter train and validate the neural network models that can then predict phenotype and pathogenicity for any single residue substitution in myosin, mybpc3, or actin. The SNP database also contains a majority of 6ddps having one or both dependent data points unknown (unfulfilled 6ddps). Unfulfilled 6ddps are predicted using the neural network models. A discrete Bayes network interprets combined fulfilled and predicted 6ddps with conditional probabilities for phenotype or pathogenicity given independent SNP characteristics. This Neural/Bayes network forecasting tests pathogenicity vs demographics of mutations in the protein domains and finds pathogenicity correlates with human subpopulations and gender. The latter implies functional cardiac motor health depends on myosin and ancillary protein components from the muscle sarcomere. In addition, the graphic realization that mybpc3 forms a third actomyosin interaction competing with the ELC N-terminus ratchet (Fig 1) implies a new strain-dependent mechanism outside myosin that contributes to motor adaptation to load. Our working models, and the integral myosin/mybpc3 motor concept, portends the wider considerations involved in understanding heart disease as a systemic maladaptation.

## Supporting information

myh7

myh6

mybpd3

actc1

myh7summary

myh6summary

mybpc3summary

actc1summary

## 6. ACKNOWLEDGEMENT

The author thanks Kevin Neff for writing the original Perl script for automated SNP extraction from the NCBI SNP database and Katalin Ajtai for scientific discussion and critical review of the manuscript.

## 7. SUPPLEMENTARY INFORMATION

Supplementary information (SI) consists of ten tables: **Tables S1-S10**, and four data sets for the fulfilled and unknown 6ddps from 2 myosins, mybpc3, and actc1. **SI Tables S7-S10** are contained in files: MYH7summary.pdf, MYH6summary.pdf, mybpc3summary.pdf, and actc1summary.pdf. **Data Sets 1-4** are contained in files: 6ddpMYH7.xls, 6ddpMYH6.xls, 6ddpMYBPC3.xls, 6ddpACTC1.xls.

## REFERENCES

[1] Murakami, K., Yasunaga, T., Noguchi, T. Q. P., Gomibuchi, Y., Ngo, K. X., Uyeda, T. Q. P., and Wakabayashi, T. (2010) Structural Basis for Actin Assembly, Activation of ATP Hydrolysis, and Delayed Phosphate Release, Cell 143, 275–287.

[2] Al-Khayat, H. A., Kensler, R. W., Squire, J. M., Marston, S. B., and Morris, E. P. (2013) Atomic model of the human cardiac muscle myosin filament, Proc. Natl. Acad. Sci. USA 110, 318–323.

[3] Rayment, I., and Holden, H. M. (1993) Myosin subfragment-1: structure and function of a molecular motor, Curr. Opin. Struct. Biol. 1993, 944–952.

[4] Lowey, S., Waller, G. S., and Trybus, K. M. (1993) Function of skeletal muscle myosin heavy and light chain isoforms by an in vitro motility assay, J. Biol. Chem. 268, 20414–20418.

[5] Sherwood, J. J., Waller, G. S., Warshaw, D. M., and Lowey, S. (2004) A point mutation in the regulatory light chain reduces the step size of skeletal muscle myosin, Proc. Natl. Acad. Sci. USA 101, 10973–10978.

[6] Pant, K., Watt, J., Greenberg, M., Jones, M., Szczesna-Cordary, D., and Moore, J. R. (2009) Removal of the cardiac myosin regulatory light chain increases isometric force production, The FASEB Journal 23, 3571–3580.

[7] Wang, L., Muthu, P., Szczesna-Cordary, D., and Kawai, M. (2013) Characterizations of myosin essential light chain’s N-terminal truncation mutant Δ43 in transgenic mouse papillary muscles by using tension transients in response to sinusoidal length alterations, J. Muscle Res. Cell Motil. 34, 93–105.

[8] Sadayappan, S., and de Tombe, P. P. (2012) Cardiac myosin binding protein-C: redefining its structure and function, Biophysical Reviews 4, 93–106.

[9] Ababou, A., Rostkova, E., Mistry, S., Masurier, C. L., Gautel, M., and Pfuhl, M. (2008) Myosin Binding Protein C Positioned to Play a Key Role in Regulation of Muscle Contraction: Structure and Interactions of Domain C1, J. Mol. Biol. 384, 615–630.

[10] Moolman-Smook, C. J., Flashman, E., de Lang, W., Li, Z., Corfield, V. A., Redwood, C., and Watkins, H. (2002) Identification of Novel Interactions Between Domains of Myosin Binding Protein-C That Are Modulated by Hypertrophic Cardiomyopathy Missense Mutations, Circ. Res. 91, 704–711.

[11] Singh, R. R., Dunn, J. W., Qadan, M. M., Hall, N., Wang, K. K., and Root, D. D. (2017) Whole length myosin binding protein C stabilizes myosin S2 as measured by gravitational force spectroscopy, Arch. Biochem. Biophys. 638, 41–51.

[12] Ratti, J., Rostkova, E., Gautel, M., and Pfuhl, M. (2011) Structure and Interactions of Myosin-binding Protein C Domain C0: Cardiac-Specific Regulation of Myosin at Its Neck?, In J. Biol. Chem., pp 12650–12658.

[13] Moos, C., Mason, C. M., Besterman, J. M., Feng, I. N. M., and Dubin, J. H. (1978) The binding of skeletal muscle C-protein to F-actin, and its relation to the interaction of actin with myosin subfragment-1, J. Mol. Biol. 124, 571–586.

[14] Whitten, A. E., Jeffries, C. M., Harris, S. P., and Trewhella, J. (2008) Cardiac myosin-binding protein C decorates F-actin: Implications for cardiac function, Proceedings of the National Academy of Sciences 105, 18360–18365.

[15] Inchingolo, A. V., Previs, S. B., Previs, M. J., Warshaw, D. M., and Kad, N. M. (2019) Revealing the mechanism of how cardiac myosin-binding protein C N-terminal fragments sensitize thin filaments for myosin binding, Proceedings of the National Academy of Sciences, 201816480.

[16] Luther, P. K., Winkler, H., Taylor, K., Zoghbi, M. E., Craig, R., Pedron, R., Squire, J. M., and Liu, J. (2011) Direct visualization of myosin-binding protein C bridging myosin and actin filament in intact muscle, Proc. Nat. Acad. Sci. USA 108, 11423–11428.

[17] Lee, K., Harris, S. P., Sadayappan, S., and Craig, R. (2015) Orientation of Myosin Binding Protein C in the Cardiac Muscle Sarcomere Determined by Domain-Specific Immuno-EM, J. Mol. Biol. 427, 274–286.

[18] Previs, M. J., Previs, S. B., Gulick, J., Robbins, J., and Warshaw, D. M. (2012) Molecular Mechanics of Cardiac Myosin-Binding Protein C in Native Thick Filaments, Science 337, 1215–1218.

[19] Walcott, S., Docken, S., and Harris, Samantha P. (2015) Effects of Cardiac Myosin Binding Protein-C on Actin Motility Are Explained with a Drag-Activation-Competition Model, Biophys. J. 108, 10–13.

[20] McNamara, J. W., Li, A., Lal, S., Bos, J. M., Harris, S. P., van der Velden, J., Ackerman, M. J., Cooke, R., and dos Remedios, C. G. (2017) MYBPC3 mutations are associated with a reduced super-relaxed state in patients with hypertrophic cardiomyopathy, PLoS ONE 12, e0180064.

[21] Kabsch, W., Mannherz, H. G., Suck, D., Pai, E. F., and Holmes, K. C. (1990) Atomic structure of the actin:DNAse I complex, Nature 347, 37–44.

[22] Aydt, E. M., Wolff, G., and Morano, I. (2007) Molecular modeling of the myosin-S1(A1) isoform, J. Struct. Biol. 159, 158–163.

[23] Lorenz, M., and Holmes, K. C. (2010) The actin-myosin interface, Proc. Nat. Acad. Sci. USA 107, 5.

[24] Ajtai, K., Garamszegi, S. P., Park, S., Velazquez Dones, A. L., and Burghardt, T. P. (2001) Structural characterization of β-cardiac myosin subfragment 1 in solution, Biochemistry 40, 12078–12093.

[25] Ajtai, K., Garamszegi, S. P., Watanabe, S., Ikebe, M., and Burghardt, T. P. (2004) The myosin cardiac loop participates functionally in the actomyosin interaction, J. Biol. Chem. 279, 23415–23421.

[26] Uyeda, T. Q. P., Ruppel, K. M., and Spudich, J. A. (1994) Enzymatic activities correlate with chimaeric substitutions at the actin-binding face of myosin, Nature 368, 567–569.

[27] Geeves, M. A., Fedorov, R., and Manstein, D. J. (2005) Molecular mechanism of actomyosin-based motility, Cell. Mol. Life Sci. 62, 1462–1477.

[28] Hayashibara, T., and Miyanishi, T. (1994) Binding of the amino-terminal region of myosin alkali l light chain to actin and its effect on actin-myosin interaction, Biochemistry 33, 12821–12827.

[29] Morano, I., Ritter, O., Bonz, A., Timek, T., Vahl, C. F., and Michel, G. (1995) Myosin light chain-actin interaction regulates cardiac contractility, Circ. Res. 76, 720–725.

[30] Wang, Y., Ajtai, K., Kazmierczak, K., Szczesna-Cordary, D., and Burghardt, T. P. (2015) N-terminus of Cardiac Myosin Essential Light Chain Modulates Myosin Step-Size, Biochemistry 55, 186–198.

[31] Wang, Y., Ajtai, K., and Burghardt Thomas, P. (2018) Cardiac and skeletal actin substrates uniquely tune cardiac myosin strain-dependent mechanics, Open Biology 8, 180143.

[32] Winkelmann, D. A., Forgacs, E., Miller, M. T., and Stock, A. M. (2015) Structural basis for drug-induced allosteric changes to human [beta]-cardiac myosin motor activity, Nature Communications 6, 7974.

[33] Planelles-Herrero, V. J., Hartman, J. J., Robert-Paganin, J., Malik, F. I., and Houdusse, A. (2017) Mechanistic and structural basis for activation of cardiac myosin force production by omecamtiv mecarbil, Nature Communications 8, 190.

[34] Burghardt, T. P., and Ajtai, K. (2018) Neural/Bayes network predictor for inheritable cardiac disease pathogenicity and phenotype, J Molec Cell Cardiol 119, 19–27.

[35] Monera, O. D., Sereda, T. J., Zhou, N. E., Kay, C. M., and Hodges, R. S. (1995) Relationship of sidechain hydrophobicity and α-helical propensity on the stability of the single-stranded amphipathic α-helix, Journal of Peptide Science 1, 319–329.

[36] Newell, K., Smith, W., Ghoshhajra, B., Isselbacher, E., Lin, A., and Lindsay, M. E. (2017) Cervical artery dissection expands the cardiovascular phenotype in FBN1-related Weill– Marchesani syndrome, American Journal of Medical Genetics Part A 173, 2551–2556.

[37] van der Linde, D., van de Laar, I. M. B. H., Bertoli-Avella, A. M., Oldenburg, R. A., Bekkers, J. A., Mattace-Raso, F. U. S., van den Meiracker, A. H., Moelker, A., van Kooten, F., Frohn-Mulder, I. M. E., Timmermans, J., Moltzer, E., Cobben, J. M., van Laer, L., Loeys, B., De Backer, J., Coucke, P. J., De Paepe, A., Hilhorst-Hofstee, Y., Wessels, M. W., and Roos-Hesselink, J. W. (2012) Aggressive Cardiovascular Phenotype of Aneurysms-Osteoarthritis Syndrome Caused by Pathogenic SMAD3 Variants, J. Am. Coll. Cardiol. 60, 397–403.

[38] Wang, Y., Ajtai, K., and Burghardt, T. P. (2013) Qdot labeled actin super-resolution motility assay measures low duty cycle muscle myosin step-size, Biochemistry 52, 1611–1621.

[39] Burghardt, T. P., Sun, X., Wang, Y., and Ajtai, K. (2017) Auxotonic to Isometric Contraction Transitioning in a Beating Heart Causes Myosin Step-Size to Down Shift, PLoS ONE 12, e0174690.

[40] Lossie, J., Köhncke, C., Mahmoodzadeh, S., Steffen, W., Canepari, M., Maffei, M., Taube, M., Larchevêque, O., Baumert, P., Haase, H., Bottinelli, R., Regitz-Zagrosek, V., and Morano, I. (2014) Molecular mechanism regulating myosin and cardiac functions by ELC, Biochem. Biophys. Res. Commun. 450, 464–469.

[41] Onishi, H., Maeda, K., Maeda, Y., Inoue, A., and Fujiwara, K. (1995) Functional chicken gizzard heavy meromyosin expression in and purification from baculovirus-infected insect cells, Proceedings of the National Academy of Sciences 92, 704–708.

[42] Uyeda, T. Q. P., Abramson, P. D., and Spudich, J. A. (1996) The neck region of the myosin motor domain acts as a lever arm to generate movement, Proc. Nat. Acad. Sci. USA 93, 4459–4464.

[43] Previs, M. J., Prosser, B. L., Mun, J. Y., Previs, S. B., Gulick, J., Lee, K., Robbins, J., Craig, R., Lederer, W. J., and Warshaw, D. M. (2015) Myosin-binding protein C corrects an intrinsic inhomogeneity in cardiac excitation-contraction coupling, Science Advances 1, e1400205.

[44] Kensler, R. W., Craig, R., and Moss, R. L. (2017) Phosphorylation of cardiac myosin binding protein C releases myosin heads from the surface of cardiac thick filaments, Proceedings of the National Academy of Sciences 114, E1355–E1364.

[45] O’Leary, T. S., Snyder, J., Sadayappan, S., Day, S. M., and Previs, M. J. (2018) MYBPC3 truncation mutations enhance actomyosin contractile mechanics in human hypertrophic cardiomyopathy, J. Mol. Cell. Cardiol.

[46] Petzhold, D., Simsek, B., Meißner, R., Mahmoodzadeh, S., and Morano, I. (2014) Distinct interactions between actin and essential myosin light chain isoforms, Biochem. Biophys. Res. Commun. 449, 284–288.

[47] Wang, Y., Yuan, C. C., Kazmierczak, K., Szczesna-Cordary, D., and Burghardt, T. P. (2018) Single Cardiac Ventricular Myosins are Autonomous Motors, Open Biology 8, 170240.

[48] Orbán, J., Lőrinczy, D., Nyitrai, M., and Hild, G. (2008) Nucleotide dependent differences between the α-skeletal and α-cardiac actin isoforms, Biochem. Biophys. Res. Commun. 368, 696–702.

[49] Kraft, T., and Montag, J. (2019) Altered force generation and cell-to-cell contractile imbalance in hypertrophic cardiomyopathy, Pflugers Arch - Eur J Physiol.

[50] Brenner, B., Seebohm, B., Tripathi, S., Montag, J., and Kraft, T. (2014) Familial hypertrophic cardiomyopathy: functional variance among individual cardiomyocytes as a trigger of FHC-phenotype development, Frontiers in Physiology 5, 392.

[51] Lange, S., Pinotsis, N., Agarkova, I., and Ehler, E. (2019) The M-band: The underestimated part of the sarcomere, Biochimica et Biophysica Acta (BBA) - Molecular Cell Research.

