## supplementary for "Demographic Model for Inheritable Cardiac Disease"

Thomas P. Burghardt

Department of Biochemistry and Molecular Biology and Physiology and Biomedical Engineering

200 First St. SW

Mayo Clinic Rochester

Rochester, MN 55905

April 2018

| MYH7 domain | code | Sequence | index | MYH6 domain | code | Sequence | index |
|---|---|---|---|---|---|---|---|
| SH3 | h3 | 28-78 | 1 | SH3 | h3 | 28-78 | 1 |
| active site | ac | <114-125, 167-178, 244-253, 260-267, 453-465, 666-673> | 2 | active site | ac | <114-125, 167-178, 245-254, 261-268, 454-466, 668-675> | 2 |
| OM binding | om | <91-92, 96, 118-119, 121, 698, 702, 705, 710-712> | 3 | OM binding | om | <91-92, 96, 118-119, 121, 700, 704, 707, 712-714> | 3 |
| Loop 1 | l1 | 202-212 | 4 | Loop 1 | l1 | 202-213 | 4 |
| C-loop | cl | 359-377 | 5 | C-loop | cl | 360-378 | 5 |
| Myopathy-loop | ml | 400-414 | 6 | Myopathy-loop | ml | 401-415 | 6 |
| Switch 2 helix | sw | 466-505 | 7 | Switch 2 helix | sw | 467-506 | 7 |
| Loop 3 | l3 | 564-576 | 8 | Loop 3 | l3 | 565-577 | 8 |
| MESA | me | 580-584 | 9 | MESA | me | 581-585 | 9 |
| Loop 2 | l2 | 622-646 | 10 | Loop 2 | l2 | 623-648 | 10 |
| actin binding | ab | <525-563, 647-655> | 11 | actin binding | ab | <526-564, 649-657> | 11 |
| SH1/SH2 hinge | sh | 683-710 | 12 | SH1/SH2 hinge | sh | 685-712 | 12 |
| Converter | cv | 711-768 | 13 | Converter | cv | 713-770 | 13 |
| IQ ELC | qe | 788-798 | 14 | IQ ELC | qe | 790-800 | 14 |
| IQ RLC | qr | 814-824 | 15 | IQ RLC | qr | 816-826 | 15 |
| Lever-arm | la | 769-843 | 16 | Lever-arm | la | 771-845 | 16 |
| S2 | s2 | 844-1356 | 17 | S2 | s2 | 846-1358 | 17 |
| LMM | lm | 1357-Cterm | 18 | LMM | lm | 1359-Cterm | 18 |
| 27k | k7 | Nterm-208 | 19 | 27k | k7 | Nterm-208 | 19 |
| 50k | k5 | 209-633 | 20 | 50k | k5 | 209-635 | 20 |
| 20k | k2 | 634-843 | 21 | 20k | k2 | 636-845 | 21 |
| N-term ELC | en | Nterm-60 | 22 | N-term ELC | en | Nterm-60 | 22 |
| EF1 ELC | e1 | 49-86 | 23 | EF1 ELC | e1 | 49-86 | 23 |
| linker ELC | e0 | 87-127 | 24 | linker ELC | e0 | 87-127 | 24 |
| EF2 ELC | e2 | 128-163 | 25 | EF2 ELC | e2 | 128-163 | 25 |
| EF3 ELC | e3 | 164-195 | 26 | EF3 ELC | e3 | 164-195 | 26 |
| RLC | rl | Nterm-Cterm | 27 | RLC | rl | Nterm-Cterm | 27 |

**Table S1.** Myosin domain list covering each residue in the sequences in 27 domains for the βmys (left) and αmys (right) isoforms. Domain assignment occurring highest in the table takes priority when sequences overlap. Myosin default domains 27k, 50k, and 20k refer to the tryptic proteolytic fragments from cleavage of the MHC sequence in Loop 1 at the active site and Loop 2 in the actin binding site [1]. A SNP falling outside all function specific domains is assigned to one of these three default domains.

| MYBPC3 domain | code | Sequence | index |
|---|---|---|---|
| phospho Ser 1 | s1 | 43-51 | 1 |
| c0-Ig like | c0 | 1-101 | 2 |
| proline rich | pr | 102-152 | 3 |
| zinc site 1 | z1 | <208, 210, 223, 225> | 4 |
| c1-Ig like | c1 | 153-256 | 5 |
| phospho Ser 2 | s2 | 271-279 | 6 |
| phospho Ser 3 | s3 | 280-288 | 7 |
| phospho Ser 4 | s4 | 300-307 | 8 |
| phospho Ser 5 | s5 | 308-315 | 9 |
| linker c1-c2 | l2 | 257-361 | 10 |
| phospho Ser 6 | s6 | 423-431 | 11 |
| c2-Ig like | c2 | 362-452 | 12 |
| c3-Ig like | c3 | 453-543 | 13 |
| phospho Ser 7 | s7 | 546-554 | 14 |
| phospho Thr 8 | s8 | 603-611 | 15 |
| c4-Ig like | c4 | 544-633 | 16 |
| linker c4-c5 | l5 | 634-644 | 17 |
| c5-Ig like | c5 | 645-711 | 18 |
| linker c5-f6 | l6 | 712-743 | 19 |
| c6-Fibronectin | f6 | 744-870 | 20 |
| c7-Fibronectin | f7 | 872-967 | 21 |
| linker c7-c8 | l8 | 968-970 | 22 |
| c8-Ig like | c8 | 971-1065 | 23 |
| c9-Fibronectin | f9 | 1066-1163 | 24 |
| linker c9-c10 | lx | 1164-1180 | 25 |
| c10-Ig like | cx | 1181-1274 | 26 |

| ACTC1 domain | code | Sequence | index |
|---|---|---|---|
| N-teminus | nt | 1-5 | 1 |
| Strand 1 | s1 | 8-9 | 2 |
| P-Loop 1 | p1 | 10-18 | 3 |
| Strand 4 | s4 | 30-37 | 4 |
| D-Loop | dl | 39-50 | 5 |
| Helix 1 | h1 | 53-60 | 6 |
| Strand 6 | s6 | 66-70 | 7 |
| Sensor-Loop | sl | 71-77 | 8 |
| Helix 2 | h2 | 78-87 | 9 |
| Proline-Rich Loop | pr | 104-112 | 10 |
| Helix 3 | h3 | 114-121 | 11 |
| Strand 9 | s9 | 153-155 | 12 |
| P-Loop 2 | p2 | 156-159 | 13 |
| Strand 10 | sx | 160-162 | 14 |
| W-Loop | wl | 166-170 | 15 |
| Ring-Rich Loop | rr | 171-180 | 16 |
| Helix 5 | h5 | 181-196 | 17 |
| Threonine-Rich Loop | tr | 197-204 | 18 |
| Helix 6 | h6 | 205-216 | 19 |
| Helix 7 | h7 | 226-230 | 20 |
| Serine-Rich Loop | sr | 231-235 | 21 |
| Strand 13 | sy | 244-250 | 22 |
| Hydrophobic Loop | hl | 262-268 | 23 |
| Helix 8b | hb | 269-281 | 24 |
| Subdomain 1 | d1 | 1-35 & 75-145 | 25 |
| Subdomain 2 | d2 | 36-74 | 26 |
| Subdomain 3 | d3 | 146-182 & 263-334 | 27 |
| Subdomain 4 | d4 | 183-262 | 28 |

**Table S2.** Mybpc3 (left) and actin (right) domain lists covering each residue in the sequences in 26 and 28 domains. Domain assignment occurring highest in the table takes priority when sequences overlap. Actin domain assignment taken mainly from Murakami et al. [2].

| residue | character | score |
|---|---|---|
| Ala | hydrophobic | +1 |
| Arg | hydrophilic | -1 |
| Asn | hydrophilic | -1 |
| Asp | hydrophilic | -1 |
| Cys | hydrophobic | +1 |
| Gln | neutral | 0 |
| Glp | hydrophilic | -1 |
| Glu | hydrophilic | -1 |
| Gly | neutral | 0 |
| His | neutral | 0 |
| Ile | very hydrophobic | +2 |
| Leu | very hydrophobic | +2 |
| Lys | hydrophilic | -1 |
| Met | very hydrophobic | +2 |
| Phe | very hydrophobic | +2 |
| Pro | hydrophilic | -1 |
| Sec | hydrophobic | +1 |
| Ser | neutral | 0 |
| Thr | neutral | 0 |
| Trp | very hydrophobic | +2 |
| Tyr | hydrophobic | +1 |
| Val | very hydrophobic | +2 |

| character | ref/sub score(Δ) | ref/sub code | index |
|---|---|---|---|
| hydrophobic/hydrophobic | 0 | mm | 1 |
| hydrophobic/hydrophilic | 2 | mp | 2 |
| hydrophobic/neutral | 1 | mn | 3 |
| hydrophobic/very hydrophobic | -1 | mh | 4 |
| hydrophilic/hydrophobic | -2 | pm | 5 |
| hydrophilic/hydrophilic | 0 | pp | 6 |
| hydrophilic/neutral | -1 | pn | 7 |
| hydrophilic/very hydrophobic | -3 | ph | 8 |
| neutral/hydrophobic | -1 | nm | 9 |
| neutral/hydrophilic | 1 | np | 10 |
| neutral/neutral | 0 | nn | 11 |
| neutral/very hydrophobic | -2 | nh | 12 |
| very hydrophobic/hydrophobic | 1 | hm | 13 |
| very hydrophobic/hydrophilic | 3 | hp | 14 |
| very hydrophobic/neutral | 2 | hn | 15 |
| very hydrophobic/very hydrophobic | 0 | hh | 16 |

**Table S3**. Amino acid residue, hydrophobicity character, and hydrophobicity integer scores in the left column matrix are derived from amphipathic α-helix models for stability [3]. Scores are applied to the mutation reference and substitution (ref/sub) pair giving the ref/sub score in the right column matrix indicated by Δ. The score implies substitution tendency towards higher (+) or lower hydrophilicity (–). The 2 letter codes for residue substitutions (ref/sub code) are indicated in the right column matrix.

| NCBI code | population |
|---|---|
| AFR | African |
| AMR | Ad Mixed American |
| ASI | Asian |
| ASJ | Ashkenazi Jewish |
| CEP | Utah EUR |
| CEU | Utah EUR |
| EAS | East Asian |
| ESP | Exome sequencing project (NHLBI) from >2x10$^5$ individuals from diverse racial and ethnic groups in the US |
| EUR | European |
| ExA | Exome Aggregation Consortium spanning 60,706 unrelated individuals worldwide |
| ExC | Exome Aggregation Consortium spanning 60,706 unrelated individuals worldwide |
| FEM | Female |
| FIN | Finnish in Finland |
| HIS | Hispanic |
| MAL | Male |
| NFE | Non-Finnish European |
| OTH | Other |
| SAS | South Asian |
| UKN | Unknown |
| YRI | Yoruba Nigeria |
| YOR | Yoruba Nigeria |

| Cps code | includes | ethnic identity subset | gender subset |
|---|---|---|---|
| AFR | AFR | ✓ | |
| AMR | AMR | ✓ | |
| ASI | ASI | ✓ | |
| ASJ | ASJ | ✓ | |
| CEP | CEP+CEU | ✓ | |
| EAS | EAS | ✓ | |
| ESP | ESP | | |
| EUR | EUR | ✓ | |
| ExA | ExA+ExC | | |
| FEM | FEM | | ✓ |
| FIN | FIN | ✓ | |
| HIS | HIS | ✓ | |
| MAL | MAL | | ✓ |
| NFE | NFE | ✓ | |
| OTH | OTH+UKN | | |
| SAS | SAS | ✓ | |
| YRI | YRI+YOR | ✓ | |
| Total | 17 | 12 | 2 |

**Table S4**. Ethnic or composite population codes and descriptions in the left column matrix are from the NCBI database. Other (OTH) category includes missense SNPs where a population category was not indicated. Similarly, NHLBI Exome sequencing (ESP) and Exome Aggregation Consortium (ExA) are aggregate missense SNP data from people in the United States and worldwide, respectively.  The right column matrix has the Consolidated Population Set (Cps) code that combines redundant populations as indicated under *includes* heading, the pure ethnic population subset from the Cps, and the gender subset from the Cps. Ethnic identity and gender subsets are used separately to probe data tendencies.

| code | allele frequency |
|---|---|
| 0 | >0.00 & ≤0.01 |
| 1 | >0.01 & ≤0.05 |
| 2 | >0.05 |

**Table S5**. Allele frequency code in the right column matrix digitizes the frequency value into 3 categories where it is ≤ 1% (code = 0), between 1 and 5% (code = 1), and >5% (code 2).

| phenotype | code |
|---|---|
| HCM | hc |
| DCM | dc |
| RCM | rc |
| left ventricle noncompaction | lv |
| cardiomyopathy | cm |
| congenital myopathy | gm |
| atrial fibrillation | af |
| ventricular fibrillation | vf |
| ventricular tachycardia | vt |
| cardiovascular phenotype | cp |
| atrial defect | ad |
| native | nv |
| uk | uk |

**Table S6**. Phenotype (ph) with 2 letter codes

**Table S7**. βmys pathogenicity vs demographics for each functional domain.

MYH7summary.pdf

**Table S8**. αmys pathogenicity vs demographics for each functional domain.

MYH6summary.pdf

**Table S9**. mybpc3 pathogenicity vs demographics for each functional domain.

mybpc3summary.pdf

**Table S10**. actc1 pathogenicity vs demographics for each functional domain.

actc1summary.pdf

**Data Sets**

1. Name: 6ddp beta myosin

Caption: Fulfilled and unknown 6 dimensional data points (6ddp) for ventricular myosin

Description: Fulfilled and unknown 6 dimensional data points (6ddp) for ventricular myosin containing mutation site, residue substitution, phenotype, and pathogenicity.

File name: 6ddpsMYH7.xls

2. Name:6ddp alpha myosin

Caption: Fulfilled and unknown 6 dimensional data points (6ddp) for atrial myosin

Description: Fulfilled and unknown 6 dimensional data points (6ddp) for atrial myosin containing mutation site, residue substitution, phenotype, and pathogenicity.

File name: 6ddpsMYH6.xls

3. Name: 6ddp actc1

Caption: Fulfilled and unknown 6 dimensional data points (6ddp) for cardiac actin

Description: Fulfilled and unknown 6 dimensional data points (6ddp) for cardiac actin containing mutation site, residue substitution, phenotype, and pathogenicity.

File name: 6ddpsACTC1.xls

4. Name: 6ddp mybpc3

Caption: Fulfilled and unknown 6 dimensional data points (6ddp) for cardiac myosin binding protein C

Description: Fulfilled and unknown 6 dimensional data points (6ddp) for cardiac myosin binding protein C containing mutation site, residue substitution, phenotype, and pathogenicity.

File name: 6ddpsMYBPC3.xls

References

[1] Balint, M., Sreter, F. A., Wolf, I., Nagy, B., and Gergely, J. (1975) The substructure of heavy meromyosin. The effect of Ca 2+ and Mg 2+ on the tryptic fragmentation of heavy meromyosin, *J. Biol. Chem. 250*, 6168-6177.

[2] Murakami, K., Yasunaga, T., Noguchi, T. Q. P., Gomibuchi, Y., Ngo, K. X., Uyeda, T. Q. P., and Wakabayashi, T. (2010) Structural Basis for Actin Assembly, Activation of ATP Hydrolysis, and Delayed Phosphate Release, *Cell 143*, 275-287.

[3] Monera, O. D., Sereda, T. J., Zhou, N. E., Kay, C. M., and Hodges, R. S. (1995) Relationship of sidechain hydrophobicity and α-helical propensity on the stability of the single-stranded amphipathic α-helix, *Journal of Peptide Science 1*, 319-329.
