## Supplementary material for "Demographic Model for Inheritable Cardiac Disease": myh7summary

{MYH7 Pathogenic Summary}

{MYH7 Benign Summary}

Pathogenic cd → ab

| $\langle P(cd \mid po)\rangle_N$ | SD | po | W(Da/Si) 24 sol | $\langle P(cd \mid af)\rangle_N$ |
|---|---|---|---|---|
| 0.001455 | 0.000044 | AMR | 72 | 0.000009 |

| $\langle\langle P(cd \mid po)\rangle_N\rangle_{po}$ | SD | SD% | x | Σx |
|---|---|---|---|---|
| 0.001455 | 0 | 0.000000 | 0.003143 | 0.003337 |

Benign cd → ab

| $\langle P(cd \mid po)\rangle_N$ | SD | po | W(Da/Si) 24 sol | $\langle P(cd \mid af)\rangle_N$ |
|---|---|---|---|---|
| 0.000090 | 0.000046 | AMR | 72 | 0.000009 |

| $\langle\langle P(cd \mid po)\rangle_N\rangle_{po}$ | SD | SD% | x | R |
|---|---|---|---|---|
| 0.000090 | 0 | 0.000000 | 0.000194 | 0 |

Pathogenic cd → ac

| $\langle P(cd \mid po)\rangle_N$ | SD | po | W(Da/Si) 24 sol | $\langle P(cd \mid af)\rangle_N$ |
|---|---|---|---|---|
| 0.001296 | 0.000157 | ASJ | 96 | 0.000012 |
| 0.001287 | 0.000176 | FEM | 96 | 0.000012 |
| 0.001271 | 0.000199 | EAS | 96 | 0.000012 |
| 0.001131 | 0.000252 | OTH | 1128 | 0.000147 |

| $\langle\langle P(cd \mid po)\rangle_N\rangle_{po}$ | SD | SD% | x | Σx |
|---|---|---|---|---|
| 0.001246 | 0.000077 | 6.204492 | 0.010765 | 0.013296 |

Benign cd → ac

| $\langle P(cd \mid po)\rangle_N$ | SD | po | W(Da/Si) 24 sol | $\langle P(cd \mid af)\rangle_N$ |
|---|---|---|---|---|
| 0.000412 | 0.000248 | OTH | 1128 | 0.000147 |
| 0.000267 | 0.000189 | EAS | 96 | 0.000012 |
| 0.000254 | 0.000163 | FEM | 96 | 0.000012 |
| 0.000239 | 0.000139 | ASJ | 96 | 0.000012 |

| $\langle\langle P(cd \mid po)\rangle_N\rangle_{po}$ | SD | SD% | x | R |
|---|---|---|---|---|
| 0.000293 | 0.000080 | 27.441630 | 0.002531 | -0.857142 |

Pathogenic cd → cl QFD 1

| $\langle P(cd \mid po)\rangle_N$ | SD | po | W(Da/Si) 24 sol | $\langle P(cd \mid af)\rangle_N$ |
|---|---|---|---|---|
| 0.001100 | 0.000268 | CEP | 24 | 0.000003 |
| 0.001067 | 0.000283 | YRI | 24 | 0.000003 |
| 0.001059 | 0.000277 | ASI | 24 | 0.000003 |
| 0.001054 | 0.000277 | HIS | 24 | 0.000696 |
| 0.000910 | 0.000298 | ExA | 96 | 0.000012 |
| 0.000896 | 0.000292 | AFR | 72 | 0.000009 |
| 0.000895 | 0.000261 | ASJ | 72 | 0.000009 |
| 0.000882 | 0.000281 | MAL | 72 | 0.000009 |
| 0.000878 | 0.000272 | FEM | 72 | 0.000009 |
| 0.000868 | 0.000307 | NFE | 72 | 0.000009 |
| 0.000847 | 0.000298 | EAS | 72 | 0.000009 |
| 0.000828 | 0.000258 | FIN | 72 | 0.000009 |
| 0.000809 | 0.000288 | AMR | 72 | 0.000009 |
| 0.000757 | 0.000290 | OTH | 288 | 0.000037 |

| $\langle\langle P(cd \mid po)\rangle_N\rangle_{po}$ | SD | SD% | x | Σx |
|---|---|---|---|---|
| 0.000918 | 0.000108 | 11.731245 | 0.027753 | 0.046176 |

Benign cd → cl QFD 1

| $\langle P(cd \mid po)\rangle_N$ | SD | po | W(Da/Si) 24 sol | $\langle P(cd \mid af)\rangle_N$ |
|---|---|---|---|---|
| 0.000782 | 0.000283 | OTH | 288 | 0.000037 |
| 0.000724 | 0.000294 | AMR | 72 | 0.000009 |
| 0.000701 | 0.000251 | FIN | 72 | 0.000009 |
| 0.000678 | 0.000303 | EAS | 72 | 0.000009 |
| 0.000665 | 0.000308 | NFE | 72 | 0.000009 |
| 0.000648 | 0.000285 | FEM | 72 | 0.000009 |
| 0.000647 | 0.000275 | MAL | 72 | 0.000009 |
| 0.000641 | 0.000289 | AFR | 72 | 0.000009 |
| 0.000628 | 0.000262 | ASJ | 72 | 0.000009 |
| 0.000599 | 0.000293 | ExA | 96 | 0.000012 |
| 0.000479 | 0.000275 | HIS | 24 | 0.000696 |
| 0.000460 | 0.000285 | YRI | 24 | 0.000003 |
| 0.000458 | 0.000275 | ASI | 24 | 0.000003 |
| 0.000420 | 0.000261 | CEP | 24 | 0.000003 |

| $\langle\langle P(cd \mid po)\rangle_N\rangle_{po}$ | SD | SD% | x | R |
|---|---|---|---|---|
| 0.000609 | 0.000111 | 18.295562 | 0.018423 | -0.955815 |

Pathogenic cd → e0

| $\langle P(cd \mid po)\rangle_N$ | SD | po | W(Da/Si) 24 sol | $\langle P(cd \mid af)\rangle_N$ |
|---|---|---|---|---|
| 0.001418 | 0.000044 | MAL | 96 | 0.000012 |
| 0.001412 | 0.000043 | AFR | 96 | 0.000012 |
| 0.001411 | 0.000049 | ASJ | 96 | 0.000012 |
| 0.001411 | 0.000037 | FEM | 96 | 0.000012 |
| 0.001401 | 0.000095 | EAS | 96 | 0.000012 |
| 0.001400 | 0.000066 | AMR | 96 | 0.000012 |
| 0.001394 | 0.000054 | ExA | 216 | 0.000028 |
| 0.001381 | 0.000040 | OTH | 360 | 0.000047 |

| $\langle\langle P(cd \mid po)\rangle_N\rangle_{po}$ | SD | SD% | x | Σx |
|---|---|---|---|---|
| 0.001404 | 0.000012 | 0.855573 | 0.024249 | 0.026685 |

Benign cd → e0

| $\langle P(cd \mid po)\rangle_N$ | SD | po | W(Da/Si) 24 sol | $\langle P(cd \mid af)\rangle_N$ |
|---|---|---|---|---|
| 0.000162 | 0.000039 | OTH | 360 | 0.000047 |
| 0.000152 | 0.000058 | ExA | 216 | 0.000028 |
| 0.000144 | 0.000095 | EAS | 96 | 0.000012 |
| 0.000144 | 0.000063 | AMR | 96 | 0.000012 |
| 0.000134 | 0.000045 | ASJ | 96 | 0.000012 |
| 0.000134 | 0.000035 | FEM | 96 | 0.000012 |
| 0.000131 | 0.000041 | AFR | 96 | 0.000012 |
| 0.000126 | 0.000042 | MAL | 96 | 0.000012 |

| $\langle\langle P(cd \mid po)\rangle_N\rangle_{po}$ | SD | SD% | x | R |
|---|---|---|---|---|
| 0.000141 | 0.000012 | 8.462547 | 0.002436 | -0.963370 |

Pathogenic cd → e2

| $\langle P(cd \mid po)\rangle_N$ | SD | po | W(Da/Si) 24 sol | $\langle P(cd \mid af)\rangle_N$ |
|---|---|---|---|---|
| 0.001372 | 0.000143 | AFR | 264 | 0.000034 |
| 0.001356 | 0.000151 | AMR | 264 | 0.000034 |
| 0.001353 | 0.000112 | OTH | 672 | 0.000087 |
| 0.001345 | 0.000170 | EAS | 264 | 0.000034 |

| $\langle\langle P(cd \mid po)\rangle_N\rangle_{po}$ | SD | SD% | x | Σx |
|---|---|---|---|---|
| 0.001357 | 0.000012 | 0.857838 | 0.011718 | 0.013355 |

Benign cd → e2

| $\langle P(cd \mid po)\rangle_N$ | SD | po | W(Da/Si) 24 sol | $\langle P(cd \mid af)\rangle_N$ |
|---|---|---|---|---|
| 0.000203 | 0.000168 | EAS | 264 | 0.000034 |
| 0.000193 | 0.000111 | OTH | 672 | 0.000087 |
| 0.000189 | 0.000151 | AMR | 264 | 0.000034 |
| 0.000173 | 0.000142 | AFR | 264 | 0.000034 |

| $\langle\langle P(cd \mid po)\rangle_N\rangle_{po}$ | SD | SD% | x | R |
|---|---|---|---|---|
| 0.000189 | 0.000012 | 6.539136 | 0.001637 | -0.970537 |

Pathogenic cd → e3

| $\langle P(cd \mid po)\rangle_N$ | SD | po | W(Da/Si) 24 sol | $\langle P(cd \mid af)\rangle_N$ |
|---|---|---|---|---|
| 0.001439 | 0.000039 | ASJ | 48 | 0.000006 |
| 0.001438 | 0.000043 | MAL | 48 | 0.000006 |
| 0.001438 | 0.000043 | AFR | 48 | 0.000006 |
| 0.001431 | 0.000031 | FEM | 48 | 0.000006 |
| 0.001429 | 0.000052 | EAS | 48 | 0.000006 |
| 0.001420 | 0.000077 | SAS | 24 | 0.000003 |
| 0.001418 | 0.000043 | AMR | 48 | 0.000006 |
| 0.001417 | 0.000032 | ExA | 144 | 0.000019 |
| 0.001397 | 0.000057 | EUR | 24 | 0.000003 |
| 0.001396 | 0.000036 | OTH | 336 | 0.000044 |

| $\langle\langle P(cd \mid po)\rangle_N\rangle_{po}$ | SD | SD% | x | Σx |
|---|---|---|---|---|
| 0.001422 | 0.000016 | 1.122468 | 0.030716 | 0.033375 |

Benign cd → e3

| $\langle P(cd \mid po)\rangle_N$ | SD | po | W(Da/Si) 24 sol | $\langle P(cd \mid af)\rangle_N$ |
|---|---|---|---|---|
| 0.000149 | 0.000059 | EUR | 24 | 0.000003 |
| 0.000149 | 0.000037 | OTH | 336 | 0.000044 |
| 0.000130 | 0.000035 | ExA | 144 | 0.000019 |
| 0.000127 | 0.000044 | AMR | 48 | 0.000006 |
| 0.000125 | 0.000079 | SAS | 24 | 0.000003 |
| 0.000116 | 0.000053 | EAS | 48 | 0.000006 |
| 0.000113 | 0.000031 | FEM | 48 | 0.000006 |
| 0.000107 | 0.000039 | ASJ | 48 | 0.000006 |
| 0.000107 | 0.000043 | MAL | 48 | 0.000006 |
| 0.000107 | 0.000043 | AFR | 48 | 0.000006 |

| $\langle\langle P(cd \mid po)\rangle_N\rangle_{po}$ | SD | SD% | x | R |
|---|---|---|---|---|
| 0.000123 | 0.000016 | 13.057821 | 0.002659 | -0.955553 |

Pathogenic cd → en QFD 2

| $\langle P(cd \mid po)\rangle_N$ | SD | po | W(Da/Si) 24 sol | $\langle P(cd \mid af)\rangle_N$ |
|---|---|---|---|---|
| 0.000631 | 0.000174 | ESP | 120 | 0.000016 |
| 0.000474 | 0.000123 | ExA | 576 | 0.000075 |
| 0.000471 | 0.000148 | MAL | 408 | 0.000053 |
| 0.000460 | 0.000146 | ASJ | 408 | 0.000053 |
| 0.000457 | 0.000122 | AFR | 456 | 0.000059 |
| 0.000455 | 0.000112 | FEM | 408 | 0.000053 |
| 0.000455 | 0.000267 | SAS | 144 | 0.000019 |
| 0.000449 | 0.000144 | NFE | 408 | 0.000053 |
| 0.000433 | 0.000146 | FIN | 408 | 0.000053 |
| 0.000414 | 0.000147 | EAS | 456 | 0.000059 |
| 0.000408 | 0.000178 | OTH | 936 | 0.000122 |
| 0.000405 | 0.000147 | AMR | 456 | 0.000059 |
| 0.000342 | 0.000177 | EUR | 144 | 0.000019 |

| $\langle\langle P(cd \mid po)\rangle_N\rangle_{po}$ | SD | SD% | x | Σx |
|---|---|---|---|---|
| 0.000450 | 0.000065 | 14.488866 | 0.012642 | 0.043394 |

Benign cd → en QFD 2

| $\langle P(cd \mid po)\rangle_N$ | SD | po | W(Da/Si) 24 sol | $\langle P(cd \mid af)\rangle_N$ |
|---|---|---|---|---|
| 0.001204 | 0.000178 | EUR | 144 | 0.000019 |
| 0.001140 | 0.000148 | AMR | 456 | 0.000059 |
| 0.001134 | 0.000180 | OTH | 936 | 0.000122 |
| 0.001134 | 0.000148 | EAS | 456 | 0.000059 |
| 0.001113 | 0.000147 | FIN | 408 | 0.000053 |
| 0.001097 | 0.000145 | NFE | 408 | 0.000053 |
| 0.001092 | 0.000268 | SAS | 144 | 0.000019 |
| 0.001091 | 0.000113 | FEM | 408 | 0.000053 |
| 0.001088 | 0.000122 | AFR | 456 | 0.000059 |
| 0.001086 | 0.000147 | ASJ | 408 | 0.000053 |
| 0.001075 | 0.000149 | MAL | 408 | 0.000053 |
| 0.001073 | 0.000126 | ExA | 576 | 0.000075 |
| 0.000913 | 0.000174 | ESP | 120 | 0.000016 |

| $\langle\langle P(cd \mid po)\rangle_N\rangle_{po}$ | SD | SD% | x | R |
|---|---|---|---|---|
| 0.001095 | 0.000066 | 5.981167 | 0.030753 | -0.828029 |

Pathogenic cd → h3

| $\langle P(cd \mid po)\rangle_N$ | SD | po | W(Da/Si) 24 sol | $\langle P(cd \mid af)\rangle_N$ |
|---|---|---|---|---|
| 0.001369 | 0.000066 | FEM | 144 | 0.000019 |
| 0.001357 | 0.000098 | FIN | 144 | 0.000019 |
| 0.001356 | 0.000071 | ASJ | 144 | 0.000019 |
| 0.001354 | 0.000096 | MAL | 144 | 0.000019 |
| 0.001349 | 0.000101 | AFR | 144 | 0.000019 |
| 0.001347 | 0.000104 | EAS | 144 | 0.000019 |
| 0.001347 | 0.000087 | ExA | 312 | 0.000041 |
| 0.001340 | 0.000093 | AMR | 144 | 0.000019 |
| 0.001307 | 0.000079 | OTH | 432 | 0.000056 |

| $\langle\langle P(cd \mid po)\rangle_N\rangle_{po}$ | SD | SD% | x | Σx |
|---|---|---|---|---|
| 0.001347 | 0.000017 | 1.267351 | 0.026188 | 0.030029 |

Benign cd → h3

| $\langle P(cd \mid po)\rangle_N$ | SD | po | W(Da/Si) 24 sol | $\langle P(cd \mid af)\rangle_N$ |
|---|---|---|---|---|
| 0.000233 | 0.000081 | OTH | 432 | 0.000056 |
| 0.000205 | 0.000094 | AMR | 144 | 0.000019 |
| 0.000200 | 0.000090 | ExA | 312 | 0.000041 |
| 0.000197 | 0.000104 | EAS | 144 | 0.000019 |
| 0.000196 | 0.000101 | AFR | 144 | 0.000019 |
| 0.000191 | 0.000072 | ASJ | 144 | 0.000019 |
| 0.000191 | 0.000096 | MAL | 144 | 0.000019 |
| 0.000188 | 0.000099 | FIN | 144 | 0.000019 |
| 0.000177 | 0.000065 | FEM | 144 | 0.000019 |

| $\langle\langle P(cd \mid po)\rangle_N\rangle_{po}$ | SD | SD% | x | R |
|---|---|---|---|---|
| 0.000198 | 0.000016 | 7.931561 | 0.003841 | -0.883938 |

Pathogenic cd → k2

| $\langle P(cd \mid po)\rangle_N$ | SD | po | W(Da/Si) 24 sol | $\langle P(cd \mid af)\rangle_N$ |
|---|---|---|---|---|
| 0.001408 | 0.000038 | ASJ | 24 | 0.000003 |
| 0.001403 | 0.000036 | AFR | 48 | 0.000006 |
| 0.001402 | 0.000050 | MAL | 24 | 0.000003 |
| 0.001398 | 0.000065 | AMR | 48 | 0.000006 |
| 0.001394 | 0.000039 | ExA | 96 | 0.000012 |
| 0.001385 | 0.000104 | SAS | 24 | 0.000003 |
| 0.001383 | 0.000084 | EAS | 48 | 0.000006 |
| 0.001379 | 0.000050 | OTH | 264 | 0.000034 |
| 0.001361 | 0.000089 | EUR | 24 | 0.000003 |

| $\langle\langle P(cd \mid po)\rangle_N\rangle_{po}$ | SD | SD% | x | Σx |
|---|---|---|---|---|
| 0.001390 | 0.000015 | 1.073068 | 0.027023 | 0.030053 |

Benign cd → k2

| $\langle P(cd \mid po)\rangle_N$ | SD | po | W(Da/Si) 24 sol | $\langle P(cd \mid af)\rangle_N$ |
|---|---|---|---|---|
| 0.000185 | 0.000090 | EUR | 24 | 0.000003 |
| 0.000168 | 0.000048 | OTH | 264 | 0.000034 |
| 0.000164 | 0.000083 | EAS | 48 | 0.000006 |
| 0.000161 | 0.000105 | SAS | 24 | 0.000003 |
| 0.000153 | 0.000041 | ExA | 96 | 0.000012 |
| 0.000148 | 0.000066 | AMR | 48 | 0.000006 |
| 0.000144 | 0.000051 | MAL | 24 | 0.000003 |
| 0.000142 | 0.000035 | AFR | 48 | 0.000006 |
| 0.000138 | 0.000037 | ASJ | 24 | 0.000003 |

| $\langle\langle P(cd \mid po)\rangle_N\rangle_{po}$ | SD | SD% | x | R |
|---|---|---|---|---|
| 0.000156 | 0.000015 | 9.674436 | 0.003029 | -0.964411 |

Pathogenic cd → k5

| $\langle P(cd \mid po)\rangle_N$ | SD | po | W(Da/Si) 24 sol | $\langle P(cd \mid af)\rangle_N$ |
|---|---|---|---|---|
| 0.001404 | 0.000051 | FEM | 312 | 0.000041 |
| 0.001398 | 0.000072 | MAL | 312 | 0.000041 |
| 0.001398 | 0.000055 | AFR | 408 | 0.000053 |
| 0.001397 | 0.000052 | ASJ | 312 | 0.000041 |
| 0.001396 | 0.000055 | FIN | 312 | 0.000041 |
| 0.001370 | 0.000078 | ExA | 1008 | 0.000131 |
| 0.001328 | 0.000114 | EUR | 120 | 0.000016 |
| 0.001272 | 0.000156 | OTH | 3816 | 0.000496 |

| $\langle\langle P(cd \mid po)\rangle_N\rangle_{po}$ | SD | SD% | x | Σx |
|---|---|---|---|---|
| 0.001370 | 0.000047 | 3.427302 | 0.023676 | 0.026634 |

Benign cd → k5

| $\langle P(cd \mid po)\rangle_N$ | SD | po | W(Da/Si) 24 sol | $\langle P(cd \mid af)\rangle_N$ |
|---|---|---|---|---|
| 0.000272 | 0.000155 | OTH | 3816 | 0.000496 |
| 0.000213 | 0.000111 | EUR | 120 | 0.000016 |
| 0.000173 | 0.000081 | ExA | 1008 | 0.000131 |
| 0.000148 | 0.000055 | FIN | 312 | 0.000041 |
| 0.000144 | 0.000070 | MAL | 312 | 0.000041 |
| 0.000142 | 0.000051 | ASJ | 312 | 0.000041 |
| 0.000140 | 0.000052 | FEM | 312 | 0.000041 |
| 0.000138 | 0.000049 | AFR | 408 | 0.000053 |

| $\langle\langle P(cd \mid po)\rangle_N\rangle_{po}$ | SD | SD% | x | R |
|---|---|---|---|---|
| 0.000171 | 0.000048 | 28.015663 | 0.002958 | -0.851197 |

Pathogenic cd → k7

| $\langle P(cd \mid po)\rangle_N$ | SD | po | W(Da/Si) 24 sol | $\langle P(cd \mid af)\rangle_N$ |
|---|---|---|---|---|
| 0.001410 | 0.000079 | FEM | 240 | 0.000031 |
| 0.001382 | 0.000111 | AMR | 312 | 0.000041 |
| 0.001365 | 0.000116 | OTH | 1392 | 0.000181 |

| $\langle\langle P(cd \mid po)\rangle_N\rangle_{po}$ | SD | SD% | x | Σx |
|---|---|---|---|---|
| 0.001385 | 0.000022 | 1.623884 | 0.008976 | 0.009993 |

Benign cd → k7

| $\langle P(cd \mid po)\rangle_N$ | SD | po | W(Da/Si) 24 sol | $\langle P(cd \mid af)\rangle_N$ |
|---|---|---|---|---|
| 0.000176 | 0.000114 | OTH | 1392 | 0.000181 |
| 0.000161 | 0.000113 | AMR | 312 | 0.000041 |
| 0.000134 | 0.000079 | FEM | 240 | 0.000031 |

| $\langle\langle P(cd \mid po)\rangle_N\rangle_{po}$ | SD | SD% | x | R |
|---|---|---|---|---|
| 0.000157 | 0.000021 | 13.454455 | 0.001017 | -0.989295 |

Pathogenic cd → l1

| $\langle P(cd \mid po)\rangle_N$ | SD | po | W(Da/Si) 24 sol | $\langle P(cd \mid af)\rangle_N$ |
|---|---|---|---|---|
| 0.001411 | 0.000053 | ASJ | 48 | 0.000006 |
| 0.001406 | 0.000070 | FIN | 48 | 0.000006 |
| 0.001406 | 0.000071 | AFR | 48 | 0.000006 |
| 0.001405 | 0.000047 | FEM | 48 | 0.000006 |
| 0.001399 | 0.000055 | AMR | 48 | 0.000006 |
| 0.001368 | 0.000064 | OTH | 192 | 0.000025 |

| $\langle\langle P(cd \mid po)\rangle_N\rangle_{po}$ | SD | SD% | x | Σx |
|---|---|---|---|---|
| 0.001399 | 0.000016 | 1.134558 | 0.018131 | 0.020000 |

Benign cd → l1

| $\langle P(cd \mid po)\rangle_N$ | SD | po | W(Da/Si) 24 sol | $\langle P(cd \mid af)\rangle_N$ |
|---|---|---|---|---|
| 0.000173 | 0.000064 | OTH | 192 | 0.000025 |
| 0.000145 | 0.000055 | AMR | 48 | 0.000006 |
| 0.000138 | 0.000071 | FIN | 48 | 0.000006 |
| 0.000138 | 0.000071 | AFR | 48 | 0.000006 |
| 0.000137 | 0.000046 | FEM | 48 | 0.000006 |
| 0.000134 | 0.000051 | ASJ | 48 | 0.000006 |

| $\langle\langle P(cd \mid po)\rangle_N\rangle_{po}$ | SD | SD% | x | R |
|---|---|---|---|---|
| 0.000144 | 0.000015 | 10.104049 | 0.001869 | -0.802778 |

Pathogenic cd → lm

| $\langle P(cd \mid po)\rangle_N$ | SD | po | W(Da/Si) 24 sol | $\langle P(cd \mid af)\rangle_N$ |
|---|---|---|---|---|
| 0.001322 | 0.000084 | ExA | 4680 | 0.000608 |
| 0.001297 | 0.000091 | AFR | 2328 | 0.000302 |
| 0.001294 | 0.000110 | MAL | 2040 | 0.000265 |
| 0.001293 | 0.000102 | NFE | 2040 | 0.000957 |
| 0.001284 | 0.000083 | FEM | 2040 | 0.000265 |
| 0.001284 | 0.000093 | OTH | 7608 | 0.000988 |
| 0.001275 | 0.000083 | EAS | 2328 | 0.000302 |
| 0.001271 | 0.000092 | ASJ | 2040 | 0.000265 |
| 0.001269 | 0.000101 | AMR | 2328 | 0.000995 |
| 0.001261 | 0.000136 | FIN | 2040 | 0.000265 |
| 0.001245 | 0.000111 | EUR | 384 | 0.000742 |

| $\langle\langle P(cd \mid po)\rangle_N\rangle_{po}$ | SD | SD% | x | Σx |
|---|---|---|---|---|
| 0.001281 | 0.000020 | 1.598311 | 0.030440 | 0.036645 |

Benign cd → lm

| $\langle P(cd \mid po)\rangle_N$ | SD | po | W(Da/Si) 24 sol | $\langle P(cd \mid af)\rangle_N$ |
|---|---|---|---|---|
| 0.000297 | 0.000110 | EUR | 384 | 0.000742 |
| 0.000282 | 0.000136 | FIN | 2040 | 0.000265 |
| 0.000277 | 0.000103 | AMR | 2328 | 0.000995 |
| 0.000270 | 0.000094 | ASJ | 2040 | 0.000265 |
| 0.000264 | 0.000089 | EAS | 2328 | 0.000302 |
| 0.000258 | 0.000084 | FEM | 2040 | 0.000265 |
| 0.000257 | 0.000093 | OTH | 7608 | 0.000988 |
| 0.000251 | 0.000100 | NFE | 2040 | 0.000957 |
| 0.000248 | 0.000109 | MAL | 2040 | 0.000265 |
| 0.000247 | 0.000090 | AFR | 2328 | 0.000302 |
| 0.000223 | 0.000088 | ExA | 4680 | 0.000608 |

| $\langle\langle P(cd \mid po)\rangle_N\rangle_{po}$ | SD | SD% | x | R |
|---|---|---|---|---|
| 0.000261 | 0.000020 | 7.661852 | 0.006204 | -0.965872 |

Pathogenic cd → ml

| $\langle P(cd \mid po)\rangle_N$ | SD | po | W(Da/Si) 24 sol | $\langle P(cd \mid af)\rangle_N$ |
|---|---|---|---|---|
| 0.001445 | 0.000039 | ASJ | 24 | 0.000003 |

| $\langle\langle P(cd \mid po)\rangle_N\rangle_{po}$ | SD | SD% | x | Σx |
|---|---|---|---|---|
| 0.001445 | 0 | 0.000000 | 0.003120 | 0.003340 |

Benign cd → ml

| $\langle P(cd \mid po)\rangle_N$ | SD | po | W(Da/Si) 24 sol | $\langle P(cd \mid af)\rangle_N$ |
|---|---|---|---|---|
| 0.000102 | 0.000039 | ASJ | 24 | 0.000003 |

| $\langle\langle P(cd \mid po)\rangle_N\rangle_{po}$ | SD | SD% | x | R |
|---|---|---|---|---|
| 0.000102 | 0 | 0.000000 | 0.000220 | 0 |

Pathogenic cd → qe

| $\langle P(cd \mid po)\rangle_N$ | SD | po | W(Da/Si) 24 sol | $\langle P(cd \mid af)\rangle_N$ |
|---|---|---|---|---|
| 0.001417 | 0.000065 | AFR | 24 | 0.000003 |
| 0.001410 | 0.000054 | EAS | 24 | 0.000003 |
| 0.001405 | 0.000059 | ExA | 96 | 0.000012 |
| 0.001398 | 0.000071 | AMR | 24 | 0.000003 |
| 0.001396 | 0.000104 | SAS | 24 | 0.000003 |
| 0.001382 | 0.000079 | EUR | 24 | 0.000003 |
| 0.001372 | 0.000084 | OTH | 96 | 0.000012 |

| $\langle\langle P(cd \mid po)\rangle_N\rangle_{po}$ | SD | SD% | x | Σx |
|---|---|---|---|---|
| 0.001397 | 0.000016 | 1.119593 | 0.021119 | 0.023369 |

Benign cd → qe

| $\langle P(cd \mid po)\rangle_N$ | SD | po | W(Da/Si) 24 sol | $\langle P(cd \mid af)\rangle_N$ |
|---|---|---|---|---|
| 0.000173 | 0.000085 | OTH | 96 | 0.000012 |
| 0.000164 | 0.000079 | EUR | 24 | 0.000003 |
| 0.000149 | 0.000105 | SAS | 24 | 0.000003 |
| 0.000148 | 0.000072 | AMR | 24 | 0.000003 |
| 0.000142 | 0.000062 | ExA | 96 | 0.000012 |
| 0.000137 | 0.000055 | EAS | 24 | 0.000003 |
| 0.000129 | 0.000064 | AFR | 24 | 0.000003 |

| $\langle\langle P(cd \mid po)\rangle_N\rangle_{po}$ | SD | SD% | x | R |
|---|---|---|---|---|
| 0.000149 | 0.000015 | 10.254578 | 0.002250 | -0.979577 |

Pathogenic cd → rl

| $\langle P(cd \mid po)\rangle_N$ | SD | po | W(Da/Si) 24 sol | $\langle P(cd \mid af)\rangle_N$ |
|---|---|---|---|---|
| 0.001367 | 0.000080 | MAL | 624 | 0.000081 |
| 0.001356 | 0.000101 | AFR | 696 | 0.000090 |
| 0.001355 | 0.000118 | FIN | 624 | 0.000081 |
| 0.001354 | 0.000134 | NFE | 624 | 0.000081 |
| 0.001353 | 0.000099 | AMR | 696 | 0.000090 |
| 0.001353 | 0.000122 | ASJ | 624 | 0.000081 |
| 0.001351 | 0.000151 | EAS | 696 | 0.000090 |
| 0.001348 | 0.000110 | FEM | 624 | 0.000081 |
| 0.001332 | 0.000162 | ExA | 1416 | 0.000184 |
| 0.001318 | 0.000148 | EUR | 120 | 0.000016 |
| 0.001164 | 0.000184 | OTH | 2616 | 0.000340 |

| $\langle\langle P(cd \mid po)\rangle_N\rangle_{po}$ | SD | SD% | x | Σx |
|---|---|---|---|---|
| 0.001332 | 0.000057 | 4.294191 | 0.031642 | 0.036400 |

Benign cd → rl

| $\langle P(cd \mid po)\rangle_N$ | SD | po | W(Da/Si) 24 sol | $\langle P(cd \mid af)\rangle_N$ |
|---|---|---|---|---|
| 0.000381 | 0.000184 | OTH | 2616 | 0.000340 |
| 0.000213 | 0.000148 | EUR | 120 | 0.000016 |
| 0.000200 | 0.000155 | ExA | 1416 | 0.000184 |
| 0.000184 | 0.000111 | FEM | 624 | 0.000081 |
| 0.000182 | 0.000135 | NFE | 624 | 0.000081 |
| 0.000180 | 0.000115 | FIN | 624 | 0.000081 |
| 0.000179 | 0.000151 | EAS | 696 | 0.000090 |
| 0.000176 | 0.000096 | AMR | 696 | 0.000090 |
| 0.000174 | 0.000104 | AFR | 696 | 0.000090 |
| 0.000173 | 0.000125 | ASJ | 624 | 0.000081 |
| 0.000161 | 0.000076 | MAL | 624 | 0.000081 |

| $\langle\langle P(cd \mid po)\rangle_N\rangle_{po}$ | SD | SD% | x | R |
|---|---|---|---|---|
| 0.000200 | 0.000061 | 30.696600 | 0.004758 | -0.665920 |

Pathogenic cd → s2

| $\langle P(cd \mid po)\rangle_N$ | SD | po | W(Da/Si) 24 sol | $\langle P(cd \mid af)\rangle_N$ |
|---|---|---|---|---|
| 0.001340 | 0.000085 | ExA | 3984 | 0.000517 |
| 0.001339 | 0.000075 | NFE | 1224 | 0.000159 |
| 0.001331 | 0.000075 | MAL | 1224 | 0.000159 |
| 0.001323 | 0.000068 | FEM | 1224 | 0.000159 |
| 0.001321 | 0.000072 | AFR | 1560 | 0.000203 |
| 0.001316 | 0.000085 | ASJ | 1224 | 0.000159 |
| 0.001315 | 0.000137 | SAS | 552 | 0.000072 |
| 0.001311 | 0.000084 | FIN | 1224 | 0.000159 |
| 0.001306 | 0.000094 | EAS | 1560 | 0.000203 |
| 0.001288 | 0.000096 | AMR | 1560 | 0.000203 |
| 0.001284 | 0.000080 | OTH | 7176 | 0.000932 |
| 0.001265 | 0.000149 | EUR | 552 | 0.000072 |

| $\langle\langle P(cd \mid po)\rangle_N\rangle_{po}$ | SD | SD% | x | Σx |
|---|---|---|---|---|
| 0.001311 | 0.000023 | 1.742219 | 0.033987 | 0.040069 |

Benign cd → s2

| $\langle P(cd \mid po)\rangle_N$ | SD | po | W(Da/Si) 24 sol | $\langle P(cd \mid af)\rangle_N$ |
|---|---|---|---|---|
| 0.000281 | 0.000149 | EUR | 552 | 0.000072 |
| 0.000263 | 0.000079 | OTH | 7176 | 0.000932 |
| 0.000258 | 0.000097 | AMR | 1560 | 0.000203 |
| 0.000241 | 0.000093 | EAS | 1560 | 0.000203 |
| 0.000235 | 0.000084 | FIN | 1224 | 0.000159 |
| 0.000232 | 0.000139 | SAS | 552 | 0.000072 |
| 0.000231 | 0.000087 | ASJ | 1224 | 0.000159 |
| 0.000225 | 0.000072 | AFR | 1560 | 0.000203 |
| 0.000223 | 0.000068 | FEM | 1224 | 0.000159 |
| 0.000215 | 0.000076 | MAL | 1224 | 0.000159 |
| 0.000207 | 0.000088 | ExA | 3984 | 0.000517 |
| 0.000207 | 0.000075 | NFE | 1224 | 0.000159 |

| $\langle\langle P(cd \mid po)\rangle_N\rangle_{po}$ | SD | SD% | x | R |
|---|---|---|---|---|
| 0.000235 | 0.000023 | 9.700920 | 0.006082 | -0.963256 |

Pathogenic cd → sh

| $\langle P(cd \mid po)\rangle_N$ | SD | po | W(Da/Si) 24 sol | $\langle P(cd \mid af)\rangle_N$ |
|---|---|---|---|---|
| 0.001447 | 0.000053 | AFR | 72 | 0.000009 |

| $\langle\langle P(cd \mid po)\rangle_N\rangle_{po}$ | SD | SD% | x | Σx |
|---|---|---|---|---|
| 0.001447 | 0 | 0.000000 | 0.003126 | 0.003337 |

Benign cd → sh

| $\langle P(cd \mid po)\rangle_N$ | SD | po | W(Da/Si) 24 sol | $\langle P(cd \mid af)\rangle_N$ |
|---|---|---|---|---|
| 0.000098 | 0.000053 | AFR | 72 | 0.000009 |

| $\langle\langle P(cd \mid po)\rangle_N\rangle_{po}$ | SD | SD% | x | R |
|---|---|---|---|---|
| 0.000098 | 0 | 0.000000 | 0.000211 | 0 |

Pathogenic cd → sw

| $\langle P(cd \mid po)\rangle_N$ | SD | po | W(Da/Si) 24 sol | $\langle P(cd \mid af)\rangle_N$ |
|---|---|---|---|---|
| 0.001453 | 0.000042 | AMR | 96 | 0.000012 |

| $\langle\langle P(cd \mid po)\rangle_N\rangle_{po}$ | SD | SD% | x | Σx |
|---|---|---|---|---|
| 0.001453 | 0 | 0.000000 | 0.003137 | 0.003323 |

Benign cd → sw

| $\langle P(cd \mid po)\rangle_N$ | SD | po | W(Da/Si) 24 sol | $\langle P(cd \mid af)\rangle_N$ |
|---|---|---|---|---|
| 0.000086 | 0.000039 | AMR | 96 | 0.000012 |

| $\langle\langle P(cd \mid po)\rangle_N\rangle_{po}$ | SD | SD% | x | R |
|---|---|---|---|---|
| 0.000086 | 0 | 0.000000 | 0.000185 | 0 |

{MYH7 R-correlation range over domains} {-0.989295, to , -0.66592}
