## Supplementary material for "Demographic Model for Inheritable Cardiac Disease": myh6summary

{MYH6 Pathogenic Summary} {MYH6 Benign Summary}

Pathogenic cd → ab

| $\langle P(cd \mid po)\rangle_N$ | SD | po | W(Da/Si) 25 sol | $\langle P(cd \mid af)\rangle_N$ |
|---|---|---|---|---|
| 0.000344 | 0.000120 | ExA | 275 | 0.000064 |
| 0.000309 | 0.000127 | AFR | 175 | 0.000041 |
| 0.000279 | 0.000096 | OTH | 475 | 0.000111 |
| 0.000274 | 0.000119 | MAL | 150 | 0.000035 |
| 0.000269 | 0.000116 | FIN | 150 | 0.000035 |
| 0.000264 | 0.000117 | AMR | 175 | 0.000041 |
| 0.000259 | 0.000098 | EAS | 175 | 0.000041 |
| 0.000258 | 0.000111 | FEM | 150 | 0.000035 |
| 0.000253 | 0.000115 | ASJ | 150 | 0.000035 |
| 0.000241 | 0.000110 | EUR | 25 | 0.000006 |
| 0.000226 | 0.000098 | NFE | 150 | 0.000035 |
| 0.000186 | 0.000096 | SAS | 25 | 0.000006 |

| $\langle\langle P(cd \mid po)\rangle_N\rangle_{po}$ | SD | SD% | x | Σx |
|---|---|---|---|---|
| 0.000264 | 0.000039 | 14.915257 | 0.006207 | 0.034889 |

Benign cd → ab

| $\langle P(cd \mid po)\rangle_N$ | SD | po | W(Da/Si) 25 sol | $\langle P(cd \mid af)\rangle_N$ |
|---|---|---|---|---|
| 0.001295 | 0.000096 | SAS | 25 | 0.000006 |
| 0.001256 | 0.000098 | NFE | 150 | 0.000035 |
| 0.001241 | 0.000110 | EUR | 25 | 0.000006 |
| 0.001228 | 0.000115 | ASJ | 150 | 0.000035 |
| 0.001224 | 0.000111 | FEM | 150 | 0.000035 |
| 0.001222 | 0.000098 | EAS | 175 | 0.000041 |
| 0.001217 | 0.000117 | AMR | 175 | 0.000041 |
| 0.001212 | 0.000116 | FIN | 150 | 0.000035 |
| 0.001207 | 0.000119 | MAL | 150 | 0.000035 |
| 0.001202 | 0.000096 | OTH | 475 | 0.000111 |
| 0.001173 | 0.000127 | AFR | 175 | 0.000041 |
| 0.001137 | 0.000120 | ExA | 275 | 0.000064 |

| $\langle\langle P(cd \mid po)\rangle_N\rangle_{po}$ | SD | SD% | x | R |
|---|---|---|---|---|
| 0.001218 | 0.000039 | 3.228053 | 0.028681 | −0.927994 |

Pathogenic cd → ac

| $\langle P(cd \mid po)\rangle_N$ | SD | po | W(Da/Si) 25 sol | $\langle P(cd \mid af)\rangle_N$ |
|---|---|---|---|---|
| 0.000378 | 0.000123 | ExA | 450 | 0.000105 |
| 0.000356 | 0.000135 | ESP | 50 | 0.000012 |
| 0.000346 | 0.000121 | AFR | 75 | 0.000018 |
| 0.000301 | 0.000120 | EAS | 75 | 0.000018 |
| 0.000301 | 0.000126 | OTH | 550 | 0.000129 |
| 0.000290 | 0.000120 | MAL | 75 | 0.000018 |
| 0.000288 | 0.000127 | AMR | 75 | 0.000018 |
| 0.000286 | 0.000127 | FIN | 75 | 0.000018 |
| 0.000282 | 0.000125 | FEM | 75 | 0.000018 |
| 0.000268 | 0.000126 | NFE | 75 | 0.000018 |
| 0.000264 | 0.000110 | ASJ | 75 | 0.000018 |

| $\langle\langle P(cd \mid po)\rangle_N\rangle_{po}$ | SD | SD% | x | Σx |
|---|---|---|---|---|
| 0.000305 | 0.000037 | 12.249388 | 0.006593 | 0.031981 |

Benign cd → ac

| $\langle P(cd \mid po)\rangle_N$ | SD | po | W(Da/Si) 25 sol | $\langle P(cd \mid af)\rangle_N$ |
|---|---|---|---|---|
| 0.001217 | 0.000110 | ASJ | 75 | 0.000018 |
| 0.001214 | 0.000126 | NFE | 75 | 0.000018 |
| 0.001200 | 0.000125 | FEM | 75 | 0.000018 |
| 0.001195 | 0.000127 | FIN | 75 | 0.000018 |
| 0.001194 | 0.000127 | AMR | 75 | 0.000018 |
| 0.001191 | 0.000120 | MAL | 75 | 0.000018 |
| 0.001180 | 0.000120 | EAS | 75 | 0.000018 |
| 0.001180 | 0.000126 | OTH | 550 | 0.000129 |
| 0.001136 | 0.000121 | AFR | 75 | 0.000018 |
| 0.001126 | 0.000135 | ESP | 50 | 0.000012 |
| 0.001104 | 0.000123 | ExA | 450 | 0.000105 |

| $\langle\langle P(cd \mid po)\rangle_N\rangle_{po}$ | SD | SD% | x | R |
|---|---|---|---|---|
| 0.001176 | 0.000037 | 3.181589 | 0.025389 | −0.930355 |

Pathogenic cd → cl

| $\langle P(cd \mid po)\rangle_N$ | SD | po | W(Da/Si) 25 sol | $\langle P(cd \mid af)\rangle_N$ |
|---|---|---|---|---|
| 0.000037 | 0.000013 | ExA | 175 | 0.000041 |

| $\langle\langle P(cd \mid po)\rangle_N\rangle_{po}$ | SD | SD% | x | Σx |
|---|---|---|---|---|
| 0.000037 | 0 | 0.000000 | 0.000073 | 0.002907 |

Benign cd → cl

| $\langle P(cd \mid po)\rangle_N$ | SD | po | W(Da/Si) 25 sol | $\langle P(cd \mid af)\rangle_N$ |
|---|---|---|---|---|
| 0.001444 | 0.000013 | ExA | 175 | 0.000041 |

| $\langle\langle P(cd \mid po)\rangle_N\rangle_{po}$ | SD | SD% | x | R |
|---|---|---|---|---|
| 0.001444 | 0 | 0.000000 | 0.002835 | 0 |

Pathogenic cd → cv

| $\langle P(cd \mid po)\rangle_N$ | SD | po | W(Da/Si) 25 sol | $\langle P(cd \mid af)\rangle_N$ |
|---|---|---|---|---|
| 0.000354 | 0.000098 | ExA | 475 | 0.000111 |
| 0.000338 | 0.000120 | ESP | 25 | 0.000006 |
| 0.000334 | 0.000102 | AFR | 275 | 0.000064 |
| 0.000297 | 0.000106 | OTH | 500 | 0.000117 |
| 0.000293 | 0.000136 | FEM | 200 | 0.000047 |
| 0.000285 | 0.000110 | MAL | 200 | 0.000047 |
| 0.000281 | 0.000098 | EAS | 275 | 0.000064 |
| 0.000277 | 0.000127 | FIN | 200 | 0.000047 |
| 0.000275 | 0.000109 | AMR | 275 | 0.000064 |
| 0.000260 | 0.000100 | ASJ | 200 | 0.000047 |
| 0.000256 | 0.000123 | NFE | 200 | 0.000047 |
| 0.000242 | 0.000106 | EUR | 125 | 0.000029 |
| 0.000232 | 0.000136 | SAS | 125 | 0.000029 |

| $\langle\langle P(cd \mid po)\rangle_N\rangle_{po}$ | SD | SD% | x | Σx |
|---|---|---|---|---|
| 0.000286 | 0.000037 | 12.950948 | 0.007308 | 0.037796 |

Benign cd → cv

| $\langle P(cd \mid po)\rangle_N$ | SD | po | W(Da/Si) 25 sol | $\langle P(cd \mid af)\rangle_N$ |
|---|---|---|---|---|
| 0.001249 | 0.000136 | SAS | 125 | 0.000029 |
| 0.001240 | 0.000106 | EUR | 125 | 0.000029 |
| 0.001225 | 0.000124 | NFE | 200 | 0.000047 |
| 0.001221 | 0.000100 | ASJ | 200 | 0.000047 |
| 0.001207 | 0.000109 | AMR | 275 | 0.000064 |
| 0.001205 | 0.000127 | FIN | 200 | 0.000047 |
| 0.001201 | 0.000098 | EAS | 275 | 0.000064 |
| 0.001197 | 0.000110 | MAL | 200 | 0.000047 |
| 0.001189 | 0.000136 | FEM | 200 | 0.000047 |
| 0.001185 | 0.000106 | OTH | 500 | 0.000117 |
| 0.001147 | 0.000102 | AFR | 275 | 0.000064 |
| 0.001143 | 0.000120 | ESP | 25 | 0.000006 |
| 0.001127 | 0.000098 | ExA | 475 | 0.000111 |

| $\langle\langle P(cd \mid po)\rangle_N\rangle_{po}$ | SD | SD% | x | R |
|---|---|---|---|---|
| 0.001195 | 0.000037 | 3.104369 | 0.030488 | −0.969156 |

Pathogenic cd → e0

| $\langle P(cd \mid po)\rangle_N$ | SD | po | W(Da/Si) 25 sol | $\langle P(cd \mid af)\rangle_N$ |
|---|---|---|---|---|
| 0.000380 | 0.000112 | ExA | 425 | 0.000099 |
| 0.000362 | 0.000120 | ESP | 75 | 0.000018 |
| 0.000351 | 0.000110 | AFR | 275 | 0.000064 |
| 0.000312 | 0.000125 | OTH | 600 | 0.000140 |
| 0.000289 | 0.000118 | FEM | 250 | 0.000059 |
| 0.000289 | 0.000116 | FIN | 250 | 0.000059 |
| 0.000286 | 0.000103 | MAL | 250 | 0.000059 |
| 0.000280 | 0.000098 | EAS | 275 | 0.000064 |
| 0.000278 | 0.000112 | AMR | 275 | 0.000064 |
| 0.000269 | 0.000123 | ASJ | 250 | 0.000059 |
| 0.000252 | 0.000089 | NFE | 250 | 0.000059 |
| 0.000247 | 0.000109 | EUR | 125 | 0.000029 |
| 0.000209 | 0.000101 | SAS | 125 | 0.000029 |

| $\langle\langle P(cd \mid po)\rangle_N\rangle_{po}$ | SD | SD% | x | Σx |
|---|---|---|---|---|
| 0.000293 | 0.000048 | 16.509300 | 0.007465 | 0.037796 |

Benign cd → e0

| $\langle P(cd \mid po)\rangle_N$ | SD | po | W(Da/Si) 25 sol | $\langle P(cd \mid af)\rangle_N$ |
|---|---|---|---|---|
| 0.001273 | 0.000101 | SAS | 125 | 0.000029 |
| 0.001235 | 0.000109 | EUR | 125 | 0.000029 |
| 0.001230 | 0.000089 | NFE | 250 | 0.000059 |
| 0.001212 | 0.000123 | ASJ | 250 | 0.000059 |
| 0.001204 | 0.000112 | AMR | 275 | 0.000064 |
| 0.001201 | 0.000098 | EAS | 275 | 0.000064 |
| 0.001195 | 0.000103 | MAL | 250 | 0.000059 |
| 0.001193 | 0.000118 | FEM | 250 | 0.000059 |
| 0.001193 | 0.000116 | FIN | 250 | 0.000059 |
| 0.001169 | 0.000125 | OTH | 600 | 0.000140 |
| 0.001131 | 0.000110 | AFR | 275 | 0.000064 |
| 0.001120 | 0.000120 | ESP | 75 | 0.000018 |
| 0.001101 | 0.000112 | ExA | 425 | 0.000099 |

| $\langle\langle P(cd \mid po)\rangle_N\rangle_{po}$ | SD | SD% | x | R |
|---|---|---|---|---|
| 0.001189 | 0.000048 | 4.063256 | 0.030331 | −0.956491 |

Pathogenic cd → e1

| $\langle P(cd \mid po)\rangle_N$ | SD | po | W(Da/Si) 25 sol | $\langle P(cd \mid af)\rangle_N$ |
|---|---|---|---|---|
| 0.000347 | 0.000126 | ExA | 625 | 0.000146 |
| 0.000344 | 0.000123 | ESP | 25 | 0.000006 |
| 0.000335 | 0.000121 | AFR | 375 | 0.000088 |
| 0.000294 | 0.000102 | OTH | 625 | 0.000146 |
| 0.000280 | 0.000131 | FEM | 325 | 0.000076 |
| 0.000269 | 0.000111 | AMR | 375 | 0.000088 |
| 0.000269 | 0.000108 | EAS | 375 | 0.000088 |
| 0.000263 | 0.000125 | FIN | 325 | 0.000076 |
| 0.000259 | 0.000114 | MAL | 325 | 0.000076 |
| 0.000253 | 0.000113 | ASJ | 325 | 0.000076 |
| 0.000244 | 0.000117 | EUR | 75 | 0.000018 |
| 0.000235 | 0.000113 | NFE | 325 | 0.000076 |
| 0.000210 | 0.000117 | SAS | 75 | 0.000018 |

| $\langle\langle P(cd \mid po)\rangle_N\rangle_{po}$ | SD | SD% | x | Σx |
|---|---|---|---|---|
| 0.000277 | 0.000042 | 15.316231 | 0.007070 | 0.037796 |

Benign cd → e1

| $\langle P(cd \mid po)\rangle_N$ | SD | po | W(Da/Si) 25 sol | $\langle P(cd \mid af)\rangle_N$ |
|---|---|---|---|---|
| 0.001272 | 0.000117 | SAS | 75 | 0.000018 |
| 0.001247 | 0.000113 | NFE | 325 | 0.000076 |
| 0.001237 | 0.000117 | EUR | 75 | 0.000018 |
| 0.001228 | 0.000113 | ASJ | 325 | 0.000076 |
| 0.001222 | 0.000114 | MAL | 325 | 0.000076 |
| 0.001219 | 0.000125 | FIN | 325 | 0.000076 |
| 0.001212 | 0.000111 | AMR | 375 | 0.000088 |
| 0.001212 | 0.000108 | EAS | 375 | 0.000088 |
| 0.001201 | 0.000131 | FEM | 325 | 0.000076 |
| 0.001188 | 0.000102 | OTH | 625 | 0.000146 |
| 0.001147 | 0.000121 | AFR | 375 | 0.000088 |
| 0.001137 | 0.000123 | ESP | 25 | 0.000006 |
| 0.001135 | 0.000126 | ExA | 625 | 0.000146 |

| $\langle\langle P(cd \mid po)\rangle_N\rangle_{po}$ | SD | SD% | x | R |
|---|---|---|---|---|
| 0.001204 | 0.000042 | 3.524151 | 0.030726 | −0.959168 |

Pathogenic cd → e2

| $\langle P(cd \mid po)\rangle_N$ | SD | po | W(Da/Si) 25 sol | $\langle P(cd \mid af)\rangle_N$ |
|---|---|---|---|---|
| 0.000359 | 0.000114 | ExA | 225 | 0.000053 |
| 0.000319 | 0.000103 | AFR | 175 | 0.000041 |
| 0.000285 | 0.000103 | OTH | 450 | 0.000105 |
| 0.000278 | 0.000113 | AMR | 175 | 0.000041 |
| 0.000270 | 0.000124 | FEM | 150 | 0.000035 |
| 0.000268 | 0.000110 | MAL | 150 | 0.000035 |
| 0.000267 | 0.000099 | EAS | 175 | 0.000041 |
| 0.000265 | 0.000120 | FIN | 150 | 0.000035 |
| 0.000257 | 0.000103 | ASJ | 150 | 0.000035 |
| 0.000236 | 0.000103 | NFE | 150 | 0.000035 |
| 0.000232 | 0.000116 | EUR | 50 | 0.000012 |
| 0.000216 | 0.000119 | SAS | 50 | 0.000012 |

| $\langle\langle P(cd \mid po)\rangle_N\rangle_{po}$ | SD | SD% | x | Σx |
|---|---|---|---|---|
| 0.000271 | 0.000039 | 14.217607 | 0.006382 | 0.034889 |

Benign cd → e2

| $\langle P(cd \mid po)\rangle_N$ | SD | po | W(Da/Si) 25 sol | $\langle P(cd \mid af)\rangle_N$ |
|---|---|---|---|---|
| 0.001265 | 0.000119 | SAS | 50 | 0.000012 |
| 0.001249 | 0.000116 | EUR | 50 | 0.000012 |
| 0.001246 | 0.000103 | NFE | 150 | 0.000035 |
| 0.001225 | 0.000103 | ASJ | 150 | 0.000035 |
| 0.001216 | 0.000120 | FIN | 150 | 0.000035 |
| 0.001215 | 0.000099 | EAS | 175 | 0.000041 |
| 0.001214 | 0.000110 | MAL | 150 | 0.000035 |
| 0.001211 | 0.000124 | FEM | 150 | 0.000035 |
| 0.001204 | 0.000113 | AMR | 175 | 0.000041 |
| 0.001196 | 0.000103 | OTH | 450 | 0.000105 |
| 0.001163 | 0.000103 | AFR | 175 | 0.000041 |
| 0.001122 | 0.000114 | ExA | 225 | 0.000053 |

| $\langle\langle P(cd \mid po)\rangle_N\rangle_{po}$ | SD | SD% | x | R |
|---|---|---|---|---|
| 0.001211 | 0.000039 | 3.182667 | 0.028507 | −0.922710 |

Pathogenic cd → e3

| $\langle P(cd \mid po)\rangle_N$ | SD | po | W(Da/Si) 25 sol | $\langle P(cd \mid af)\rangle_N$ |
|---|---|---|---|---|
| 0.000360 | 0.000102 | ExA | 250 | 0.000059 |
| 0.000344 | 0.000118 | AFR | 175 | 0.000950 |
| 0.000337 | 0.000122 | ESP | 25 | 0.000915 |
| 0.000277 | 0.000116 | OTH | 425 | 0.000099 |
| 0.000275 | 0.000120 | FIN | 150 | 0.000035 |
| 0.000273 | 0.000116 | FEM | 150 | 0.000035 |
| 0.000270 | 0.000110 | MAL | 150 | 0.000035 |
| 0.000268 | 0.000110 | AMR | 175 | 0.000041 |
| 0.000265 | 0.000099 | EAS | 175 | 0.000041 |
| 0.000253 | 0.000117 | ASJ | 150 | 0.000035 |
| 0.000231 | 0.000111 | NFE | 150 | 0.000035 |
| 0.000227 | 0.000106 | EUR | 50 | 0.000012 |
| 0.000212 | 0.000114 | SAS | 50 | 0.000012 |

| $\langle\langle P(cd \mid po)\rangle_N\rangle_{po}$ | SD | SD% | x | Σx |
|---|---|---|---|---|
| 0.000276 | 0.000045 | 16.432883 | 0.007053 | 0.037796 |

Benign cd → e3

| $\langle P(cd \mid po)\rangle_N$ | SD | po | W(Da/Si) 25 sol | $\langle P(cd \mid af)\rangle_N$ |
|---|---|---|---|---|
| 0.001269 | 0.000114 | SAS | 50 | 0.000012 |
| 0.001254 | 0.000106 | EUR | 50 | 0.000012 |
| 0.001251 | 0.000111 | NFE | 150 | 0.000035 |
| 0.001228 | 0.000117 | ASJ | 150 | 0.000035 |
| 0.001216 | 0.000099 | EAS | 175 | 0.000041 |
| 0.001214 | 0.000110 | AMR | 175 | 0.000041 |
| 0.001211 | 0.000110 | MAL | 150 | 0.000035 |
| 0.001209 | 0.000116 | FEM | 150 | 0.000035 |
| 0.001206 | 0.000120 | FIN | 150 | 0.000035 |
| 0.001205 | 0.000116 | OTH | 425 | 0.000099 |
| 0.001144 | 0.000122 | ESP | 25 | 0.000915 |
| 0.001137 | 0.000118 | AFR | 175 | 0.000950 |
| 0.001121 | 0.000102 | ExA | 250 | 0.000059 |

| $\langle\langle P(cd \mid po)\rangle_N\rangle_{po}$ | SD | SD% | x | R |
|---|---|---|---|---|
| 0.001205 | 0.000045 | 3.769981 | 0.030743 | −0.939713 |

Pathogenic cd → en QFD 1

| $\langle P(cd \mid po)\rangle_N$ | SD | po | W(Da/Si) 25 sol | $\langle P(cd \mid af)\rangle_N$ |
|---|---|---|---|---|
| 0.000874 | 0.000187 | ESP | 100 | 0.000023 |
| 0.000817 | 0.000193 | ExA | 550 | 0.000129 |
| 0.000772 | 0.000129 | OTH | 975 | 0.000228 |
| 0.000740 | 0.000202 | AFR | 375 | 0.000088 |
| 0.000699 | 0.000134 | CEP | 25 | 0.000006 |
| 0.000689 | 0.000221 | FEM | 350 | 0.000082 |
| 0.000679 | 0.000204 | EAS | 375 | 0.000088 |
| 0.000673 | 0.000213 | EUR | 50 | 0.000012 |
| 0.000659 | 0.000215 | FIN | 350 | 0.000082 |
| 0.000656 | 0.000180 | MAL | 350 | 0.000082 |
| 0.000653 | 0.000163 | ASJ | 350 | 0.000082 |
| 0.000621 | 0.000157 | AMR | 375 | 0.000088 |
| 0.000621 | 0.000277 | SAS | 50 | 0.000012 |
| 0.000606 | 0.000180 | NFE | 350 | 0.000082 |

| $\langle\langle P(cd \mid po)\rangle_N\rangle_{po}$ | SD | SD% | x | Σx |
|---|---|---|---|---|
| 0.000697 | 0.000078 | 11.186760 | 0.019150 | 0.040689 |

Benign cd → en QFD 1

| $\langle P(cd \mid po)\rangle_N$ | SD | po | W(Da/Si) 25 sol | $\langle P(cd \mid af)\rangle_N$ |
|---|---|---|---|---|
| 0.000874 | 0.000179 | NFE | 350 | 0.000082 |
| 0.000860 | 0.000277 | SAS | 50 | 0.000012 |
| 0.000858 | 0.000160 | AMR | 375 | 0.000088 |
| 0.000828 | 0.000163 | ASJ | 350 | 0.000082 |
| 0.000826 | 0.000180 | MAL | 350 | 0.000082 |
| 0.000822 | 0.000215 | FIN | 350 | 0.000082 |
| 0.000809 | 0.000213 | EUR | 50 | 0.000012 |
| 0.000801 | 0.000205 | EAS | 375 | 0.000088 |
| 0.000791 | 0.000224 | FEM | 350 | 0.000082 |
| 0.000783 | 0.000134 | CEP | 25 | 0.000006 |
| 0.000741 | 0.000203 | AFR | 375 | 0.000088 |
| 0.000710 | 0.000129 | OTH | 975 | 0.000228 |
| 0.000664 | 0.000193 | ExA | 550 | 0.000129 |
| 0.000607 | 0.000187 | ESP | 100 | 0.000023 |

| $\langle\langle P(cd \mid po)\rangle_N\rangle_{po}$ | SD | SD% | x | R |
|---|---|---|---|---|
| 0.000784 | 0.000078 | 9.912670 | 0.021539 | −0.933396 |

Pathogenic cd → h3

| $\langle P(cd \mid po)\rangle_N$ | SD | po | W(Da/Si) 25 sol | $\langle P(cd \mid af)\rangle_N$ |
|---|---|---|---|---|
| 0.000221 | 0.000079 | AFR | 400 | 0.007045 |
| 0.000214 | 0.000085 | YRI | 25 | 0.006048 |
| 0.000212 | 0.000076 | ExA | 575 | 0.006177 |
| 0.000212 | 0.000082 | ESP | 75 | 0.006969 |
| 0.000196 | 0.000097 | FEM | 400 | 0.006136 |
| 0.000185 | 0.000085 | FIN | 400 | 0.006136 |
| 0.000174 | 0.000068 | MAL | 400 | 0.006136 |
| 0.000167 | 0.000072 | ASJ | 400 | 0.006136 |
| 0.000164 | 0.000078 | OTH | 775 | 0.006224 |
| 0.000158 | 0.000067 | NFE | 400 | 0.006136 |
| 0.000158 | 0.000059 | CEP | 25 | 0.006048 |
| 0.000144 | 0.000071 | EUR | 50 | 0.006054 |

| $\langle\langle P(cd \mid po)\rangle_N\rangle_{po}$ | SD | SD% | x | Σx |
|---|---|---|---|---|
| 0.000184 | 0.000026 | 14.398687 | 0.004330 | 0.034889 |

Benign cd → h3

| $\langle P(cd \mid po)\rangle_N$ | SD | po | W(Da/Si) 25 sol | $\langle P(cd \mid af)\rangle_N$ |
|---|---|---|---|---|
| 0.001337 | 0.000071 | EUR | 50 | 0.006054 |
| 0.001323 | 0.000067 | NFE | 400 | 0.006136 |
| 0.001323 | 0.000059 | CEP | 25 | 0.006048 |
| 0.001317 | 0.000078 | OTH | 775 | 0.006224 |
| 0.001315 | 0.000072 | ASJ | 400 | 0.006136 |
| 0.001307 | 0.000068 | MAL | 400 | 0.006136 |
| 0.001296 | 0.000085 | FIN | 400 | 0.006136 |
| 0.001285 | 0.000097 | FEM | 400 | 0.006136 |
| 0.001269 | 0.000076 | ExA | 575 | 0.006177 |
| 0.001269 | 0.000082 | ESP | 75 | 0.006969 |
| 0.001268 | 0.000085 | YRI | 25 | 0.006048 |
| 0.001261 | 0.000079 | AFR | 400 | 0.007045 |

| $\langle\langle P(cd \mid po)\rangle_N\rangle_{po}$ | SD | SD% | x | R |
|---|---|---|---|---|
| 0.001298 | 0.000026 | 2.039198 | 0.030559 | −0.982520 |

Pathogenic cd → k2 QFD 2

| $\langle P(cd \mid po)\rangle_N$ | SD | po | W(Da/Si) 25 sol | $\langle P(cd \mid af)\rangle_N$ |
|---|---|---|---|---|
| 0.001060 | 0.000107 | ExA | 125 | 0.000029 |
| 0.000954 | 0.000155 | AFR | 25 | 0.000006 |
| 0.000868 | 0.000201 | FEM | 25 | 0.000006 |
| 0.000859 | 0.000186 | FIN | 25 | 0.000006 |
| 0.000849 | 0.000171 | ESP | 25 | 0.000006 |
| 0.000831 | 0.000201 | NFE | 25 | 0.000006 |
| 0.000828 | 0.000212 | MAL | 25 | 0.000006 |
| 0.000806 | 0.000214 | AMR | 25 | 0.000006 |
| 0.000800 | 0.000172 | EAS | 25 | 0.000006 |
| 0.000799 | 0.000201 | ASJ | 25 | 0.000006 |
| 0.000793 | 0.000168 | OTH | 250 | 0.000059 |
| 0.000725 | 0.000184 | EUR | 25 | 0.000006 |
| 0.000672 | 0.000204 | SAS | 25 | 0.000006 |

| $\langle\langle P(cd \mid po)\rangle_N\rangle_{po}$ | SD | SD% | x | Σx |
|---|---|---|---|---|
| 0.000834 | 0.000096 | 11.532038 | 0.021282 | 0.037794 |

Benign cd → k2 QFD 2

| $\langle P(cd \mid po)\rangle_N$ | SD | po | W(Da/Si) 25 sol | $\langle P(cd \mid af)\rangle_N$ |
|---|---|---|---|---|
| 0.000810 | 0.000204 | SAS | 25 | 0.000006 |
| 0.000757 | 0.000184 | EUR | 25 | 0.000006 |
| 0.000689 | 0.000168 | OTH | 250 | 0.000059 |
| 0.000683 | 0.000201 | ASJ | 25 | 0.000006 |
| 0.000682 | 0.000172 | EAS | 25 | 0.000006 |
| 0.000675 | 0.000214 | AMR | 25 | 0.000006 |
| 0.000653 | 0.000212 | MAL | 25 | 0.000006 |
| 0.000651 | 0.000201 | NFE | 25 | 0.000006 |
| 0.000632 | 0.000171 | ESP | 25 | 0.000006 |
| 0.000622 | 0.000186 | FIN | 25 | 0.000006 |
| 0.000612 | 0.000200 | FEM | 25 | 0.000006 |
| 0.000527 | 0.000155 | AFR | 25 | 0.000006 |
| 0.000421 | 0.000107 | ExA | 125 | 0.000029 |

| $\langle\langle P(cd \mid po)\rangle_N\rangle_{po}$ | SD | SD% | x | R |
|---|---|---|---|---|
| 0.000647 | 0.000096 | 14.868964 | 0.016512 | −0.909975 |

Pathogenic cd → k5 QFD 3

| $\langle P(cd \mid po)\rangle_N$ | SD | po | W(Da/Si) 25 sol | $\langle P(cd \mid af)\rangle_N$ |
|---|---|---|---|---|
| 0.000726 | 0.000160 | ExA | 2000 | 0.000468 |
| 0.000691 | 0.000186 | AFR | 1150 | 0.000269 |
| 0.000615 | 0.000241 | FEM | 900 | 0.000211 |
| 0.000590 | 0.000197 | AMR | 1150 | 0.000269 |
| 0.000581 | 0.000176 | FIN | 900 | 0.000211 |
| 0.000580 | 0.000139 | ESP | 250 | 0.000059 |
| 0.000570 | 0.000209 | NFE | 900 | 0.000211 |
| 0.000565 | 0.000200 | ASJ | 900 | 0.000211 |
| 0.000557 | 0.000177 | MAL | 900 | 0.000211 |
| 0.000554 | 0.000166 | EAS | 1150 | 0.000269 |
| 0.000500 | 0.000171 | OTH | 1925 | 0.000451 |
| 0.000463 | 0.000153 | EUR | 525 | 0.000123 |
| 0.000440 | 0.000176 | SAS | 525 | 0.000123 |

| $\langle\langle P(cd \mid po)\rangle_N\rangle_{po}$ | SD | SD% | x | Σx |
|---|---|---|---|---|
| 0.000572 | 0.000079 | 13.870498 | 0.014588 | 0.037796 |

Benign cd → k5 QFD 3

| $\langle P(cd \mid po)\rangle_N$ | SD | po | W(Da/Si) 25 sol | $\langle P(cd \mid af)\rangle_N$ |
|---|---|---|---|---|
| 0.001042 | 0.000176 | SAS | 525 | 0.000123 |
| 0.001019 | 0.000153 | EUR | 525 | 0.000123 |
| 0.000981 | 0.000171 | OTH | 1925 | 0.000451 |
| 0.000927 | 0.000166 | EAS | 1150 | 0.000269 |
| 0.000925 | 0.000177 | MAL | 900 | 0.000211 |
| 0.000916 | 0.000200 | ASJ | 900 | 0.000211 |
| 0.000911 | 0.000209 | NFE | 900 | 0.000211 |
| 0.000901 | 0.000139 | ESP | 250 | 0.000059 |
| 0.000900 | 0.000176 | FIN | 900 | 0.000211 |
| 0.000891 | 0.000198 | AMR | 1150 | 0.000269 |
| 0.000867 | 0.000241 | FEM | 900 | 0.000211 |
| 0.000790 | 0.000186 | AFR | 1150 | 0.000269 |
| 0.000756 | 0.000160 | ExA | 2000 | 0.000468 |

| $\langle\langle P(cd \mid po)\rangle_N\rangle_{po}$ | SD | SD% | x | R |
|---|---|---|---|---|
| 0.000910 | 0.000079 | 8.717313 | 0.023208 | −0.942042 |

Pathogenic cd → k7

| $\langle P(cd \mid po)\rangle_N$ | SD | po | W(Da/Si) 25 sol | $\langle P(cd \mid af)\rangle_N$ |
|---|---|---|---|---|
| 0.000351 | 0.000120 | ExA | 1100 | 0.000257 |
| 0.000335 | 0.000124 | ESP | 100 | 0.000023 |
| 0.000323 | 0.000102 | AFR | 600 | 0.000140 |
| 0.000302 | 0.000095 | OTH | 1150 | 0.000269 |
| 0.000277 | 0.000121 | FIN | 475 | 0.000111 |
| 0.000275 | 0.000121 | MAL | 475 | 0.000111 |
| 0.000270 | 0.000116 | AMR | 600 | 0.000140 |
| 0.000267 | 0.000124 | FEM | 475 | 0.000111 |
| 0.000264 | 0.000084 | CEP | 25 | 0.000006 |
| 0.000262 | 0.000113 | EAS | 600 | 0.000140 |
| 0.000259 | 0.000112 | ASJ | 475 | 0.000111 |
| 0.000241 | 0.000098 | NFE | 475 | 0.000111 |
| 0.000235 | 0.000096 | EUR | 225 | 0.000053 |
| 0.000215 | 0.000125 | SAS | 225 | 0.000053 |

| $\langle\langle P(cd \mid po)\rangle_N\rangle_{po}$ | SD | SD% | x | Σx |
|---|---|---|---|---|
| 0.000277 | 0.000039 | 13.941795 | 0.007604 | 0.040703 |

Benign cd → k7

| $\langle P(cd \mid po)\rangle_N$ | SD | po | W(Da/Si) 25 sol | $\langle P(cd \mid af)\rangle_N$ |
|---|---|---|---|---|
| 0.001267 | 0.000125 | SAS | 225 | 0.000053 |
| 0.001247 | 0.000096 | EUR | 225 | 0.000053 |
| 0.001241 | 0.000098 | NFE | 475 | 0.000111 |
| 0.001222 | 0.000112 | ASJ | 475 | 0.000111 |
| 0.001220 | 0.000113 | EAS | 600 | 0.000140 |
| 0.001217 | 0.000084 | CEP | 25 | 0.000006 |
| 0.001215 | 0.000124 | FEM | 475 | 0.000111 |
| 0.001211 | 0.000116 | AMR | 600 | 0.000140 |
| 0.001206 | 0.000121 | MAL | 475 | 0.000111 |
| 0.001205 | 0.000121 | FIN | 475 | 0.000111 |
| 0.001180 | 0.000095 | OTH | 1150 | 0.000269 |
| 0.001158 | 0.000102 | AFR | 600 | 0.000140 |
| 0.001147 | 0.000124 | ESP | 100 | 0.000023 |
| 0.001131 | 0.000120 | ExA | 1100 | 0.000257 |

| $\langle\langle P(cd \mid po)\rangle_N\rangle_{po}$ | SD | SD% | x | R |
|---|---|---|---|---|
| 0.001205 | 0.000039 | 3.203191 | 0.033099 | −0.957640 |

Pathogenic cd → l1

| $\langle P(cd \mid po)\rangle_N$ | SD | po | W(Da/Si) 25 sol | $\langle P(cd \mid af)\rangle_N$ |
|---|---|---|---|---|
| 0.000349 | 0.000122 | ExA | 150 | 0.000035 |
| 0.000344 | 0.000136 | ESP | 25 | 0.000006 |
| 0.000328 | 0.000116 | AFR | 150 | 0.000035 |
| 0.000293 | 0.000122 | OTH | 250 | 0.000059 |
| 0.000282 | 0.000126 | AMR | 150 | 0.000035 |
| 0.000274 | 0.000112 | EAS | 150 | 0.000035 |
| 0.000267 | 0.000133 | FEM | 125 | 0.000029 |
| 0.000267 | 0.000123 | MAL | 125 | 0.000029 |
| 0.000267 | 0.000125 | FIN | 125 | 0.000029 |
| 0.000258 | 0.000103 | ASJ | 125 | 0.000029 |
| 0.000244 | 0.000097 | NFE | 125 | 0.000029 |
| 0.000236 | 0.000130 | EUR | 50 | 0.000012 |
| 0.000223 | 0.000130 | SAS | 50 | 0.000012 |

| $\langle\langle P(cd \mid po)\rangle_N\rangle_{po}$ | SD | SD% | x | Σx |
|---|---|---|---|---|
| 0.000279 | 0.000040 | 14.215991 | 0.007128 | 0.037796 |

Benign cd → l1

| $\langle P(cd \mid po)\rangle_N$ | SD | po | W(Da/Si) 25 sol | $\langle P(cd \mid af)\rangle_N$ |
|---|---|---|---|---|
| 0.001258 | 0.000130 | SAS | 50 | 0.000012 |
| 0.001246 | 0.000130 | EUR | 50 | 0.000012 |
| 0.001237 | 0.000097 | NFE | 125 | 0.000029 |
| 0.001223 | 0.000103 | ASJ | 125 | 0.000029 |
| 0.001215 | 0.000133 | FEM | 125 | 0.000029 |
| 0.001215 | 0.000123 | MAL | 125 | 0.000029 |
| 0.001215 | 0.000125 | FIN | 125 | 0.000029 |
| 0.001207 | 0.000112 | EAS | 150 | 0.000035 |
| 0.001200 | 0.000125 | AMR | 150 | 0.000035 |
| 0.001189 | 0.000122 | OTH | 250 | 0.000059 |
| 0.001153 | 0.000116 | AFR | 150 | 0.000035 |
| 0.001137 | 0.000136 | ESP | 25 | 0.000006 |
| 0.001132 | 0.000122 | ExA | 150 | 0.000035 |

| $\langle\langle P(cd \mid po)\rangle_N\rangle_{po}$ | SD | SD% | x | R |
|---|---|---|---|---|
| 0.001202 | 0.000040 | 3.304237 | 0.030668 | −0.956461 |

Pathogenic cd → l2

| $\langle P(cd \mid po)\rangle_N$ | SD | po | W(Da/Si) 25 sol | $\langle P(cd \mid af)\rangle_N$ |
|---|---|---|---|---|
| 0.000358 | 0.000122 | ExA | 225 | 0.000053 |
| 0.000320 | 0.000115 | AFR | 100 | 0.000023 |
| 0.000299 | 0.000122 | OTH | 250 | 0.000059 |
| 0.000279 | 0.000122 | MAL | 75 | 0.000018 |
| 0.000278 | 0.000127 | FEM | 75 | 0.000018 |
| 0.000273 | 0.000121 | FIN | 75 | 0.000018 |
| 0.000262 | 0.000115 | AMR | 100 | 0.000023 |
| 0.000258 | 0.000117 | ASJ | 75 | 0.000018 |
| 0.000254 | 0.000091 | EAS | 100 | 0.000023 |
| 0.000238 | 0.000105 | NFE | 75 | 0.000018 |
| 0.000221 | 0.000093 | EUR | 50 | 0.000012 |
| 0.000211 | 0.000113 | SAS | 50 | 0.000012 |

| $\langle\langle P(cd \mid po)\rangle_N\rangle_{po}$ | SD | SD% | x | Σx |
|---|---|---|---|---|
| 0.000271 | 0.000041 | 15.149670 | 0.006379 | 0.034889 |

Benign cd → l2

| $\langle P(cd \mid po)\rangle_N$ | SD | po | W(Da/Si) 25 sol | $\langle P(cd \mid af)\rangle_N$ |
|---|---|---|---|---|
| 0.001270 | 0.000113 | SAS | 50 | 0.000012 |
| 0.001260 | 0.000093 | EUR | 50 | 0.000012 |
| 0.001243 | 0.000105 | NFE | 75 | 0.000018 |
| 0.001227 | 0.000091 | EAS | 100 | 0.000023 |
| 0.001223 | 0.000117 | ASJ | 75 | 0.000018 |
| 0.001220 | 0.000115 | AMR | 100 | 0.000023 |
| 0.001209 | 0.000121 | FIN | 75 | 0.000018 |
| 0.001204 | 0.000127 | FEM | 75 | 0.000018 |
| 0.001202 | 0.000122 | MAL | 75 | 0.000018 |
| 0.001183 | 0.000122 | OTH | 250 | 0.000059 |
| 0.001162 | 0.000115 | AFR | 100 | 0.000023 |
| 0.001123 | 0.000122 | ExA | 225 | 0.000053 |

| $\langle\langle P(cd \mid po)\rangle_N\rangle_{po}$ | SD | SD% | x | R |
|---|---|---|---|---|
| 0.001211 | 0.000041 | 3.389952 | 0.028509 | −0.959134 |

Pathogenic cd → l3

| $\langle P(cd \mid po)\rangle_N$ | SD | po | W(Da/Si) 25 sol | $\langle P(cd \mid af)\rangle_N$ |
|---|---|---|---|---|
| 0.000389 | 0.000124 | ExA | 75 | 0.000018 |
| 0.000369 | 0.000128 | ESP | 25 | 0.000006 |
| 0.000332 | 0.000112 | AFR | 50 | 0.000012 |
| 0.000306 | 0.000117 | OTH | 175 | 0.000041 |
| 0.000295 | 0.000138 | FEM | 50 | 0.000012 |
| 0.000293 | 0.000135 | FIN | 50 | 0.000012 |
| 0.000288 | 0.000115 | AMR | 50 | 0.000012 |
| 0.000284 | 0.000112 | MAL | 50 | 0.000012 |
| 0.000278 | 0.000105 | EAS | 50 | 0.000012 |
| 0.000272 | 0.000123 | ASJ | 50 | 0.000012 |
| 0.000262 | 0.000132 | NFE | 50 | 0.000012 |

| $\langle\langle P(cd \mid po)\rangle_N\rangle_{po}$ | SD | SD% | x | Σx |
|---|---|---|---|---|
| 0.000306 | 0.000041 | 13.310980 | 0.006607 | 0.031981 |

Benign cd → l3

| $\langle P(cd \mid po)\rangle_N$ | SD | po | W(Da/Si) 25 sol | $\langle P(cd \mid af)\rangle_N$ |
|---|---|---|---|---|
| 0.001220 | 0.000132 | NFE | 50 | 0.000012 |
| 0.001210 | 0.000123 | ASJ | 50 | 0.000012 |
| 0.001204 | 0.000105 | EAS | 50 | 0.000012 |
| 0.001198 | 0.000112 | MAL | 50 | 0.000012 |
| 0.001194 | 0.000115 | AMR | 50 | 0.000012 |
| 0.001189 | 0.000135 | FIN | 50 | 0.000012 |
| 0.001186 | 0.000138 | FEM | 50 | 0.000012 |
| 0.001175 | 0.000117 | OTH | 175 | 0.000041 |
| 0.001149 | 0.000113 | AFR | 50 | 0.000012 |
| 0.001112 | 0.000128 | ESP | 25 | 0.000006 |
| 0.001093 | 0.000124 | ExA | 75 | 0.000018 |

| $\langle\langle P(cd \mid po)\rangle_N\rangle_{po}$ | SD | SD% | x | R |
|---|---|---|---|---|
| 0.001175 | 0.000041 | 3.466963 | 0.025374 | −0.918233 |

Pathogenic cd → la QFD 4

| $\langle P(cd \mid po)\rangle_N$ | SD | po | W(Da/Si) 25 sol | $\langle P(cd \mid af)\rangle_N$ |
|---|---|---|---|---|
| 0.000635 | 0.000167 | ESP | 25 | 0.000006 |
| 0.000602 | 0.000193 | ExA | 525 | 0.000123 |
| 0.000583 | 0.000176 | AFR | 375 | 0.000088 |
| 0.000576 | 0.000132 | OTH | 700 | 0.000164 |
| 0.000532 | 0.000177 | FEM | 250 | 0.000059 |
| 0.000507 | 0.000194 | MAL | 250 | 0.000059 |
| 0.000505 | 0.000229 | FIN | 250 | 0.000059 |
| 0.000503 | 0.000178 | AMR | 375 | 0.000088 |
| 0.000501 | 0.000178 | EUR | 225 | 0.000053 |
| 0.000496 | 0.000147 | EAS | 375 | 0.000088 |
| 0.000490 | 0.000148 | ASJ | 250 | 0.000059 |
| 0.000451 | 0.000193 | NFE | 250 | 0.000059 |
| 0.000438 | 0.000195 | SAS | 225 | 0.000053 |

| $\langle\langle P(cd \mid po)\rangle_N\rangle_{po}$ | SD | SD% | x | Σx |
|---|---|---|---|---|
| 0.000525 | 0.000058 | 11.123742 | 0.013382 | 0.037796 |

Benign cd → la QFD 4

| $\langle P(cd \mid po)\rangle_N$ | SD | po | W(Da/Si) 25 sol | $\langle P(cd \mid af)\rangle_N$ |
|---|---|---|---|---|
| 0.001043 | 0.000195 | SAS | 225 | 0.000053 |
| 0.001031 | 0.000193 | NFE | 250 | 0.000059 |
| 0.000991 | 0.000148 | ASJ | 250 | 0.000059 |
| 0.000985 | 0.000147 | EAS | 375 | 0.000088 |
| 0.000980 | 0.000178 | EUR | 225 | 0.000053 |
| 0.000979 | 0.000178 | AMR | 375 | 0.000088 |
| 0.000977 | 0.000229 | FIN | 250 | 0.000059 |
| 0.000974 | 0.000194 | MAL | 250 | 0.000059 |
| 0.000949 | 0.000177 | FEM | 250 | 0.000059 |
| 0.000906 | 0.000132 | OTH | 700 | 0.000164 |
| 0.000899 | 0.000177 | AFR | 375 | 0.000088 |
| 0.000879 | 0.000193 | ExA | 525 | 0.000123 |
| 0.000847 | 0.000167 | ESP | 25 | 0.000006 |

| $\langle\langle P(cd \mid po)\rangle_N\rangle_{po}$ | SD | SD% | x | R |
|---|---|---|---|---|
| 0.000957 | 0.000058 | 6.098053 | 0.024414 | −0.958229 |

Pathogenic cd → lm

| $\langle P(cd \mid po)\rangle_N$ | SD | po | W(Da/Si) 25 sol | $\langle P(cd \mid af)\rangle_N$ |
|---|---|---|---|---|
| 0.000181 | 0.000223 | ExA | 6300 | 0.003293 |
| 0.000104 | 0.000135 | OTH | 7525 | 0.001761 |

| $\langle\langle P(cd \mid po)\rangle_N\rangle_{po}$ | SD | SD% | x | Σx |
|---|---|---|---|---|
| 0.000143 | 0.000054 | 37.908434 | 0.000560 | 0.005815 |

Benign cd → lm

| $\langle P(cd \mid po)\rangle_N$ | SD | po | W(Da/Si) 25 sol | $\langle P(cd \mid af)\rangle_N$ |
|---|---|---|---|---|
| 0.001377 | 0.000135 | OTH | 7525 | 0.001761 |
| 0.001301 | 0.000223 | ExA | 6300 | 0.003293 |

| $\langle\langle P(cd \mid po)\rangle_N\rangle_{po}$ | SD | SD% | x | R |
|---|---|---|---|---|
| 0.001339 | 0.000054 | 4.036868 | 0.005255 | 0 |

Pathogenic cd → me

| $\langle P(cd \mid po)\rangle_N$ | SD | po | W(Da/Si) 25 sol | $\langle P(cd \mid af)\rangle_N$ |
|---|---|---|---|---|
| 0.000381 | 0.000105 | ExA | 225 | 0.000053 |
| 0.000362 | 0.000123 | ESP | 75 | 0.000018 |
| 0.000346 | 0.000123 | AFR | 200 | 0.000047 |
| 0.000322 | 0.000115 | OTH | 225 | 0.000053 |
| 0.000309 | 0.000128 | AMR | 200 | 0.000047 |
| 0.000298 | 0.000125 | FEM | 150 | 0.000035 |
| 0.000295 | 0.000115 | EAS | 200 | 0.000047 |
| 0.000285 | 0.000116 | FIN | 150 | 0.000035 |
| 0.000283 | 0.000120 | MAL | 150 | 0.000035 |
| 0.000274 | 0.000119 | ASJ | 150 | 0.000035 |
| 0.000259 | 0.000121 | EUR | 50 | 0.000012 |
| 0.000258 | 0.000102 | NFE | 150 | 0.000035 |
| 0.000222 | 0.000118 | SAS | 50 | 0.000012 |

| $\langle\langle P(cd \mid po)\rangle_N\rangle_{po}$ | SD | SD% | x | Σx |
|---|---|---|---|---|
| 0.000300 | 0.000045 | 14.866442 | 0.007642 | 0.037796 |

Benign cd → me

| $\langle P(cd \mid po)\rangle_N$ | SD | po | W(Da/Si) 25 sol | $\langle P(cd \mid af)\rangle_N$ |
|---|---|---|---|---|
| 0.001259 | 0.000118 | SAS | 50 | 0.000012 |
| 0.001223 | 0.000102 | NFE | 150 | 0.000035 |
| 0.001222 | 0.000121 | EUR | 50 | 0.000012 |
| 0.001207 | 0.000119 | ASJ | 150 | 0.000035 |
| 0.001199 | 0.000120 | MAL | 150 | 0.000035 |
| 0.001196 | 0.000116 | FIN | 150 | 0.000035 |
| 0.001186 | 0.000115 | EAS | 200 | 0.000047 |
| 0.001184 | 0.000125 | FEM | 150 | 0.000035 |
| 0.001173 | 0.000128 | AMR | 200 | 0.000047 |
| 0.001159 | 0.000115 | OTH | 225 | 0.000053 |
| 0.001136 | 0.000123 | AFR | 200 | 0.000047 |
| 0.001120 | 0.000123 | ESP | 75 | 0.000018 |
| 0.001100 | 0.000105 | ExA | 225 | 0.000053 |

| $\langle\langle P(cd \mid po)\rangle_N\rangle_{po}$ | SD | SD% | x | R |
|---|---|---|---|---|
| 0.001182 | 0.000045 | 3.767291 | 0.030154 | −0.974758 |

Pathogenic cd → ml

| $\langle P(cd \mid po)\rangle_N$ | SD | po | W(Da/Si) 25 sol | $\langle P(cd \mid af)\rangle_N$ |
|---|---|---|---|---|
| 0.000363 | 0.000130 | ExA | 75 | 0.000018 |
| 0.000320 | 0.000104 | AFR | 100 | 0.000023 |
| 0.000288 | 0.000112 | OTH | 150 | 0.000035 |
| 0.000274 | 0.000118 | AMR | 100 | 0.000023 |
| 0.000273 | 0.000109 | MAL | 75 | 0.000018 |
| 0.000270 | 0.000133 | FEM | 75 | 0.000018 |
| 0.000267 | 0.000130 | FIN | 75 | 0.000018 |
| 0.000265 | 0.000098 | EAS | 100 | 0.000023 |
| 0.000257 | 0.000104 | ASJ | 75 | 0.000018 |
| 0.000242 | 0.000107 | NFE | 75 | 0.000018 |
| 0.000235 | 0.000100 | EUR | 25 | 0.000006 |
| 0.000202 | 0.000106 | SAS | 25 | 0.000006 |

| $\langle\langle P(cd \mid po)\rangle_N\rangle_{po}$ | SD | SD% | x | Σx |
|---|---|---|---|---|
| 0.000271 | 0.000041 | 15.010175 | 0.006389 | 0.034889 |

Benign cd → ml

| $\langle P(cd \mid po)\rangle_N$ | SD | po | W(Da/Si) 25 sol | $\langle P(cd \mid af)\rangle_N$ |
|---|---|---|---|---|
| 0.001279 | 0.000106 | SAS | 25 | 0.000006 |
| 0.001247 | 0.000100 | EUR | 25 | 0.000006 |
| 0.001240 | 0.000107 | NFE | 75 | 0.000018 |
| 0.001225 | 0.000104 | ASJ | 75 | 0.000018 |
| 0.001216 | 0.000098 | EAS | 100 | 0.000023 |
| 0.001215 | 0.000130 | FIN | 75 | 0.000018 |
| 0.001211 | 0.000133 | FEM | 75 | 0.000018 |
| 0.001209 | 0.000109 | MAL | 75 | 0.000018 |
| 0.001207 | 0.000118 | AMR | 100 | 0.000023 |
| 0.001194 | 0.000112 | OTH | 150 | 0.000035 |
| 0.001162 | 0.000104 | AFR | 100 | 0.000023 |
| 0.001119 | 0.000130 | ExA | 75 | 0.000018 |

| $\langle\langle P(cd \mid po)\rangle_N\rangle_{po}$ | SD | SD% | x | R |
|---|---|---|---|---|
| 0.001210 | 0.000041 | 3.364804 | 0.028500 | −0.917680 |

Pathogenic cd → om

| $\langle P(cd \mid po)\rangle_N$ | SD | po | W(Da/Si) 25 sol | $\langle P(cd \mid af)\rangle_N$ |
|---|---|---|---|---|
| 0.000365 | 0.000117 | ExA | 125 | 0.000029 |
| 0.000328 | 0.000106 | AFR | 25 | 0.000006 |
| 0.000291 | 0.000107 | OTH | 50 | 0.000012 |
| 0.000289 | 0.000124 | AMR | 25 | 0.000006 |
| 0.000288 | 0.000100 | EAS | 25 | 0.000006 |
| 0.000273 | 0.000124 | FEM | 25 | 0.000006 |
| 0.000273 | 0.000113 | MAL | 25 | 0.000006 |
| 0.000272 | 0.000127 | FIN | 25 | 0.000006 |
| 0.000249 | 0.000101 | ASJ | 25 | 0.000006 |
| 0.000244 | 0.000102 | NFE | 25 | 0.000006 |

| $\langle\langle P(cd \mid po)\rangle_N\rangle_{po}$ | SD | SD% | x | Σx |
|---|---|---|---|---|
| 0.000287 | 0.000036 | 12.577595 | 0.005638 | 0.029074 |

Benign cd → om

| $\langle P(cd \mid po)\rangle_N$ | SD | po | W(Da/Si) 25 sol | $\langle P(cd \mid af)\rangle_N$ |
|---|---|---|---|---|
| 0.001237 | 0.000102 | NFE | 25 | 0.000006 |
| 0.001232 | 0.000101 | ASJ | 25 | 0.000006 |
| 0.001210 | 0.000127 | FIN | 25 | 0.000006 |
| 0.001209 | 0.000124 | FEM | 25 | 0.000006 |
| 0.001209 | 0.000113 | MAL | 25 | 0.000006 |
| 0.001194 | 0.000099 | EAS | 25 | 0.000006 |
| 0.001193 | 0.000124 | AMR | 25 | 0.000006 |
| 0.001190 | 0.000107 | OTH | 50 | 0.000012 |
| 0.001153 | 0.000106 | AFR | 25 | 0.000006 |
| 0.001116 | 0.000117 | ExA | 125 | 0.000029 |

| $\langle\langle P(cd \mid po)\rangle_N\rangle_{po}$ | SD | SD% | x | R |
|---|---|---|---|---|
| 0.001194 | 0.000036 | 3.026080 | 0.023436 | −0.915977 |

Pathogenic cd → qe QFD 5

| $\langle P(cd \mid po)\rangle_N$ | SD | po | W(Da/Si) 25 sol | $\langle P(cd \mid af)\rangle_N$ |
|---|---|---|---|---|
| 0.000662 | 0.000176 | ExA | 200 | 0.000047 |
| 0.000641 | 0.000259 | AFR | 150 | 0.000035 |
| 0.000602 | 0.000226 | OTH | 175 | 0.000041 |
| 0.000540 | 0.000212 | EAS | 150 | 0.000035 |
| 0.000540 | 0.000240 | EUR | 75 | 0.000018 |
| 0.000532 | 0.000195 | FEM | 125 | 0.000029 |
| 0.000531 | 0.000240 | FIN | 125 | 0.000029 |
| 0.000528 | 0.000209 | ASJ | 125 | 0.000029 |
| 0.000523 | 0.000221 | MAL | 125 | 0.000029 |
| 0.000507 | 0.000189 | AMR | 150 | 0.000035 |
| 0.000473 | 0.000255 | NFE | 125 | 0.000029 |
| 0.000473 | 0.000295 | SAS | 75 | 0.000018 |

| $\langle\langle P(cd \mid po)\rangle_N\rangle_{po}$ | SD | SD% | x | Σx |
|---|---|---|---|---|
| 0.000546 | 0.000060 | 10.928648 | 0.012859 | 0.034889 |

Benign cd → qe QFD 5

| $\langle P(cd \mid po)\rangle_N$ | SD | po | W(Da/Si) 25 sol | $\langle P(cd \mid af)\rangle_N$ |
|---|---|---|---|---|
| 0.001009 | 0.000255 | NFE | 125 | 0.000029 |
| 0.001009 | 0.000295 | SAS | 75 | 0.000018 |
| 0.000974 | 0.000189 | AMR | 150 | 0.000035 |
| 0.000958 | 0.000221 | MAL | 125 | 0.000029 |
| 0.000953 | 0.000209 | ASJ | 125 | 0.000029 |
| 0.000951 | 0.000240 | FIN | 125 | 0.000029 |
| 0.000949 | 0.000195 | FEM | 125 | 0.000029 |
| 0.000942 | 0.000212 | EAS | 150 | 0.000035 |
| 0.000942 | 0.000240 | EUR | 75 | 0.000018 |
| 0.000879 | 0.000226 | OTH | 175 | 0.000041 |
| 0.000841 | 0.000259 | AFR | 150 | 0.000035 |
| 0.000819 | 0.000176 | ExA | 200 | 0.000047 |

| $\langle\langle P(cd \mid po)\rangle_N\rangle_{po}$ | SD | SD% | x | R |
|---|---|---|---|---|
| 0.000935 | 0.000060 | 6.379631 | 0.022030 | −0.923986 |

Pathogenic cd → qr

| $\langle P(cd \mid po)\rangle_N$ | SD | po | W(Da/Si) 25 sol | $\langle P(cd \mid af)\rangle_N$ |
|---|---|---|---|---|
| 0.000046 | 0.000033 | OTH | 150 | 0.000035 |

| $\langle\langle P(cd \mid po)\rangle_N\rangle_{po}$ | SD | SD% | x | Σx |
|---|---|---|---|---|
| 0.000046 | 0 | 0.000000 | 0.000090 | 0.002907 |

Benign cd → qr

| $\langle P(cd \mid po)\rangle_N$ | SD | po | W(Da/Si) 25 sol | $\langle P(cd \mid af)\rangle_N$ |
|---|---|---|---|---|
| 0.001436 | 0.000033 | OTH | 150 | 0.000035 |

| $\langle\langle P(cd \mid po)\rangle_N\rangle_{po}$ | SD | SD% | x | R |
|---|---|---|---|---|
| 0.001436 | 0 | 0.000000 | 0.002818 | 0 |

Pathogenic cd → rl

| $\langle P(cd \mid po)\rangle_N$ | SD | po | W(Da/Si) 25 sol | $\langle P(cd \mid af)\rangle_N$ |
|---|---|---|---|---|
| 0.000352 | 0.000099 | ExA | 2450 | 0.000573 |
| 0.000326 | 0.000127 | ESP | 275 | 0.000064 |
| 0.000322 | 0.000117 | AFR | 1550 | 0.012447 |
| 0.000290 | 0.000113 | OTH | 3575 | 0.012921 |
| 0.000274 | 0.000118 | MAL | 1300 | 0.007256 |
| 0.000269 | 0.000091 | YRI | 75 | 0.012102 |
| 0.000264 | 0.000128 | FIN | 1300 | 0.006346 |
| 0.000264 | 0.000091 | CEP | 100 | 0.006066 |
| 0.000263 | 0.000127 | FEM | 1300 | 0.007256 |
| 0.000259 | 0.000109 | AMR | 1550 | 0.007314 |
| 0.000258 | 0.000085 | EAS | 1550 | 0.006405 |
| 0.000253 | 0.000103 | ASJ | 1300 | 0.002123 |
| 0.000238 | 0.000098 | NFE | 1300 | 0.001213 |
| 0.000225 | 0.000097 | EUR | 500 | 0.006159 |
| 0.000207 | 0.000121 | SAS | 500 | 0.006159 |

| $\langle\langle P(cd \mid po)\rangle_N\rangle_{po}$ | SD | SD% | x | Σx |
|---|---|---|---|---|
| 0.000271 | 0.000038 | 14.152758 | 0.007978 | 0.043611 |

Benign cd → rl

| $\langle P(cd \mid po)\rangle_N$ | SD | po | W(Da/Si) 25 sol | $\langle P(cd \mid af)\rangle_N$ |
|---|---|---|---|---|
| 0.001274 | 0.000121 | SAS | 500 | 0.006159 |
| 0.001257 | 0.000097 | EUR | 500 | 0.006159 |
| 0.001243 | 0.000098 | NFE | 1300 | 0.001213 |
| 0.001228 | 0.000103 | ASJ | 1300 | 0.002123 |
| 0.001223 | 0.000085 | EAS | 1550 | 0.006405 |
| 0.001222 | 0.000109 | AMR | 1550 | 0.007314 |
| 0.001219 | 0.000127 | FEM | 1300 | 0.007256 |
| 0.001217 | 0.000128 | FIN | 1300 | 0.006346 |
| 0.001217 | 0.000091 | CEP | 100 | 0.006066 |
| 0.001212 | 0.000091 | YRI | 75 | 0.012102 |
| 0.001207 | 0.000118 | MAL | 1300 | 0.007256 |
| 0.001191 | 0.000113 | OTH | 3575 | 0.012921 |
| 0.001159 | 0.000117 | AFR | 1550 | 0.012447 |
| 0.001156 | 0.000127 | ESP | 275 | 0.000064 |
| 0.001130 | 0.000099 | ExA | 2450 | 0.000573 |

| $\langle\langle P(cd \mid po)\rangle_N\rangle_{po}$ | SD | SD% | x | R |
|---|---|---|---|---|
| 0.001210 | 0.000038 | 3.169170 | 0.035632 | −0.939204 |

Pathogenic cd → s2

| $\langle P(cd \mid po)\rangle_N$ | SD | po | W(Da/Si) 25 sol | $\langle P(cd \mid af)\rangle_N$ |
|---|---|---|---|---|
| 0.000273 | 0.000102 | ExA | 4850 | 0.013220 |
| 0.000258 | 0.000081 | AFR | 3100 | 0.003453 |
| 0.000218 | 0.000078 | ESP | 625 | 0.001965 |
| 0.000211 | 0.000049 | YRI | 50 | 0.000921 |
| 0.000207 | 0.000093 | FIN | 2800 | 0.012740 |
| 0.000205 | 0.000101 | FEM | 2800 | 0.012740 |
| 0.000200 | 0.000072 | EAS | 3125 | 0.013725 |
| 0.000198 | 0.000060 | CEP | 50 | 0.012096 |
| 0.000197 | 0.000077 | ASJ | 2775 | 0.006692 |
| 0.000196 | 0.000069 | AMR | 3125 | 0.012816 |
| 0.000192 | 0.000063 | MAL | 2800 | 0.012740 |
| 0.000186 | 0.000065 | NFE | 2800 | 0.012740 |
| 0.000173 | 0.000072 | OTH | 6725 | 0.013658 |
| 0.000165 | 0.000050 | EUR | 1000 | 0.012318 |
| 0.000147 | 0.000061 | SAS | 1000 | 0.012318 |

| $\langle\langle P(cd \mid po)\rangle_N\rangle_{po}$ | SD | SD% | x | Σx |
|---|---|---|---|---|
| 0.000202 | 0.000032 | 15.821586 | 0.005937 | 0.043611 |

Benign cd → s2

| $\langle P(cd \mid po)\rangle_N$ | SD | po | W(Da/Si) 25 sol | $\langle P(cd \mid af)\rangle_N$ |
|---|---|---|---|---|
| 0.001335 | 0.000061 | SAS | 1000 | 0.012318 |
| 0.001317 | 0.000050 | EUR | 1000 | 0.012318 |
| 0.001308 | 0.000072 | OTH | 6725 | 0.013658 |
| 0.001295 | 0.000065 | NFE | 2800 | 0.012740 |
| 0.001289 | 0.000063 | MAL | 2800 | 0.012740 |
| 0.001286 | 0.000069 | AMR | 3125 | 0.012816 |
| 0.001284 | 0.000077 | ASJ | 2775 | 0.006692 |
| 0.001284 | 0.000060 | CEP | 50 | 0.012096 |
| 0.001282 | 0.000072 | EAS | 3125 | 0.013725 |
| 0.001277 | 0.000101 | FEM | 2800 | 0.012740 |
| 0.001275 | 0.000093 | FIN | 2800 | 0.012740 |
| 0.001270 | 0.000049 | YRI | 50 | 0.000921 |
| 0.001264 | 0.000078 | ESP | 625 | 0.001965 |
| 0.001223 | 0.000081 | AFR | 3100 | 0.003453 |
| 0.001208 | 0.000102 | ExA | 4850 | 0.013220 |

| $\langle\langle P(cd \mid po)\rangle_N\rangle_{po}$ | SD | SD% | x | R |
|---|---|---|---|---|
| 0.001280 | 0.000032 | 2.492903 | 0.037674 | −0.916277 |

Pathogenic cd → sh

| $\langle P(cd \mid po)\rangle_N$ | SD | po | W(Da/Si) 25 sol | $\langle P(cd \mid af)\rangle_N$ |
|---|---|---|---|---|
| 0.000368 | 0.000114 | ExA | 300 | 0.000070 |
| 0.000341 | 0.000120 | ESP | 25 | 0.000006 |
| 0.000333 | 0.000102 | AFR | 100 | 0.000933 |
| 0.000290 | 0.000103 | OTH | 225 | 0.000053 |
| 0.000279 | 0.000132 | FIN | 100 | 0.000023 |
| 0.000275 | 0.000114 | MAL | 100 | 0.000023 |
| 0.000274 | 0.000129 | FEM | 100 | 0.000023 |
| 0.000267 | 0.000108 | AMR | 100 | 0.000023 |
| 0.000267 | 0.000092 | EAS | 100 | 0.000023 |
| 0.000258 | 0.000112 | ASJ | 100 | 0.000023 |
| 0.000241 | 0.000102 | NFE | 100 | 0.000023 |
| 0.000235 | 0.000105 | EUR | 25 | 0.000006 |
| 0.000209 | 0.000099 | SAS | 25 | 0.000006 |

| $\langle\langle P(cd \mid po)\rangle_N\rangle_{po}$ | SD | SD% | x | Σx |
|---|---|---|---|---|
| 0.000280 | 0.000045 | 15.974996 | 0.007135 | 0.037796 |

Benign cd → sh

| $\langle P(cd \mid po)\rangle_N$ | SD | po | W(Da/Si) 25 sol | $\langle P(cd \mid af)\rangle_N$ |
|---|---|---|---|---|
| 0.001273 | 0.000099 | SAS | 25 | 0.000006 |
| 0.001247 | 0.000105 | EUR | 25 | 0.000006 |
| 0.001241 | 0.000102 | NFE | 100 | 0.000023 |
| 0.001223 | 0.000112 | ASJ | 100 | 0.000023 |
| 0.001215 | 0.000108 | AMR | 100 | 0.000023 |
| 0.001215 | 0.000092 | EAS | 100 | 0.000023 |
| 0.001207 | 0.000129 | FEM | 100 | 0.000023 |
| 0.001206 | 0.000114 | MAL | 100 | 0.000023 |
| 0.001202 | 0.000132 | FIN | 100 | 0.000023 |
| 0.001191 | 0.000103 | OTH | 225 | 0.000053 |
| 0.001148 | 0.000102 | AFR | 100 | 0.000933 |
| 0.001141 | 0.000120 | ESP | 25 | 0.000006 |
| 0.001114 | 0.000114 | ExA | 300 | 0.000070 |

| $\langle\langle P(cd \mid po)\rangle_N\rangle_{po}$ | SD | SD% | x | R |
|---|---|---|---|---|
| 0.001202 | 0.000045 | 3.717868 | 0.030661 | −0.951371 |

Pathogenic cd → sw

| $\langle P(cd \mid po)\rangle_N$ | SD | po | W(Da/Si) 25 sol | $\langle P(cd \mid af)\rangle_N$ |
|---|---|---|---|---|
| 0.000305 | 0.000092 | ExA | 250 | 0.000059 |
| 0.000260 | 0.000088 | AFR | 75 | 0.000018 |
| 0.000220 | 0.000061 | FIN | 50 | 0.000012 |
| 0.000216 | 0.000068 | FEM | 50 | 0.000012 |
| 0.000205 | 0.000084 | ASJ | 50 | 0.000012 |
| 0.000201 | 0.000062 | MAL | 50 | 0.000012 |
| 0.000178 | 0.000063 | EUR | 25 | 0.000006 |
| 0.000146 | 0.000070 | OTH | 350 | 0.000082 |

| $\langle\langle P(cd \mid po)\rangle_N\rangle_{po}$ | SD | SD% | x | Σx |
|---|---|---|---|---|
| 0.000216 | 0.000049 | 22.541162 | 0.003397 | 0.023259 |

Benign cd → sw

| $\langle P(cd \mid po)\rangle_N$ | SD | po | W(Da/Si) 25 sol | $\langle P(cd \mid af)\rangle_N$ |
|---|---|---|---|---|
| 0.001336 | 0.000070 | OTH | 350 | 0.000082 |
| 0.001304 | 0.000063 | EUR | 25 | 0.000006 |
| 0.001280 | 0.000062 | MAL | 50 | 0.000012 |
| 0.001277 | 0.000084 | ASJ | 50 | 0.000012 |
| 0.001266 | 0.000068 | FEM | 50 | 0.000012 |
| 0.001262 | 0.000061 | FIN | 50 | 0.000012 |
| 0.001221 | 0.000088 | AFR | 75 | 0.000018 |
| 0.001177 | 0.000092 | ExA | 250 | 0.000059 |

| $\langle\langle P(cd \mid po)\rangle_N\rangle_{po}$ | SD | SD% | x | R |
|---|---|---|---|---|
| 0.001265 | 0.000049 | 3.854459 | 0.019862 | −0.953742 |

{MYH6 R−correlation range over domains} {−0.98252, to , −0.909975}
