## Supplementary material for "Demographic Model for Inheritable Cardiac Disease": mybpc3summary

{MYBPC3 Pathogenic Summary}

Pathogenic cd → c0 QFD 1

| $\langle P(cd\mid po)\rangle_N$ | SD | po | W(Da/Si) 25 sol | $\langle P(cd\mid af)\rangle_N$ |
|---|---|---|---|---|
| 0.000903 | 0.000194 | ExA | 925 | 0.000242 |
| 0.000888 | 0.000169 | OTH | 1675 | 0.000439 |
| 0.000834 | 0.000167 | MAL | 650 | 0.000170 |
| 0.000819 | 0.000167 | FIN | 650 | 0.000170 |
| 0.000816 | 0.000161 | EAS | 800 | 0.000210 |
| 0.000808 | 0.000184 | FEM | 650 | 0.000170 |
| 0.000804 | 0.000180 | AMR | 800 | 0.000210 |
| 0.000802 | 0.000175 | ASJ | 650 | 0.000170 |
| 0.000798 | 0.000176 | AFR | 800 | 0.000210 |
| 0.000797 | 0.000186 | NFE | 650 | 0.000170 |
| 0.000782 | 0.000137 | EUR | 250 | 0.000066 |
| 0.000674 | 0.000164 | SAS | 250 | 0.000066 |

| $\langle\langle P(cd\mid po)\rangle_N\rangle_{po}$ | SD | SD% | x | Σx |
|---|---|---|---|---|
| 0.000810 | 0.000056 | 6.957113 | 0.021238 | 0.040318 |

Pathogenic cd → c1 QFD 2

| $\langle P(cd\mid po)\rangle_N$ | SD | po | W(Da/Si) 25 sol | $\langle P(cd\mid af)\rangle_N$ |
|---|---|---|---|---|
| 0.001268 | 0.000086 | ExA | 1175 | 0.007957 |
| 0.001256 | 0.000081 | OTH | 2025 | 0.012937 |
| 0.001227 | 0.000108 | MAL | 700 | 0.007833 |
| 0.001218 | 0.000098 | AFR | 750 | 0.002992 |
| 0.001214 | 0.000089 | CEP | 50 | 0.007662 |
| 0.001212 | 0.000117 | AMR | 750 | 0.004953 |
| 0.001209 | 0.000128 | FIN | 700 | 0.007833 |
| 0.001208 | 0.000130 | FEM | 700 | 0.007833 |
| 0.001208 | 0.000135 | EAS | 750 | 0.001128 |
| 0.001203 | 0.000117 | ASJ | 700 | 0.007833 |
| 0.001203 | 0.000127 | NFE | 700 | 0.007833 |
| 0.001203 | 0.000107 | EUR | 325 | 0.007735 |
| 0.001104 | 0.000146 | SAS | 325 | 0.004842 |
| 0.000922 | 0.000165 | YRI | 25 | 0.000938 |

| $\langle\langle P(cd\mid po)\rangle_N\rangle_{po}$ | SD | SD% | x | Σx |
|---|---|---|---|---|
| 0.001190 | 0.000085 | 7.162088 | 0.036366 | 0.047013 |

Pathogenic cd → c2

| $\langle P(cd\mid po)\rangle_N$ | SD | po | W(Da/Si) 25 sol | $\langle P(cd\mid af)\rangle_N$ |
|---|---|---|---|---|
| 0.001298 | 0.000222 | OTH | 1200 | 0.000315 |
| 0.001265 | 0.000266 | EAS | 425 | 0.000111 |
| 0.001260 | 0.000255 | MAL | 350 | 0.001024 |
| 0.001250 | 0.000273 | FEM | 350 | 0.001024 |
| 0.001242 | 0.000282 | ASJ | 350 | 0.000092 |
| 0.001242 | 0.000257 | EUR | 125 | 0.000033 |
| 0.001238 | 0.000281 | AFR | 425 | 0.001043 |
| 0.001236 | 0.000281 | FIN | 350 | 0.000092 |
| 0.001227 | 0.000289 | NFE | 350 | 0.000092 |
| 0.001159 | 0.000315 | SAS | 125 | 0.000033 |
| 0.001051 | 0.000394 | YRI | 25 | 0.003831 |

| $\langle\langle P(cd\mid po)\rangle_N\rangle_{po}$ | SD | SD% | x | Σx |
|---|---|---|---|---|
| 0.001225 | 0.000067 | 5.431082 | 0.029416 | 0.036959 |

Pathogenic cd → c3 QFD 3

| $\langle P(cd\mid po)\rangle_N$ | SD | po | W(Da/Si) 25 sol | $\langle P(cd\mid af)\rangle_N$ |
|---|---|---|---|---|
| 0.000621 | 0.000211 | ExA | 1100 | 0.000288 |
| 0.000600 | 0.000188 | OTH | 1325 | 0.000347 |
| 0.000518 | 0.000207 | MAL | 550 | 0.000144 |
| 0.000499 | 0.000192 | FEM | 550 | 0.000144 |
| 0.000496 | 0.000233 | EAS | 550 | 0.000144 |
| 0.000493 | 0.000200 | AMR | 550 | 0.000144 |
| 0.000480 | 0.000182 | NFE | 550 | 0.000144 |
| 0.000478 | 0.000168 | FIN | 550 | 0.000144 |
| 0.000465 | 0.000195 | ASJ | 550 | 0.000144 |
| 0.000464 | 0.000169 | AFR | 550 | 0.000144 |
| 0.000437 | 0.000172 | EUR | 150 | 0.000039 |
| 0.000341 | 0.000162 | SAS | 150 | 0.000039 |
| 0.000196 | 0.000077 | YRI | 50 | 0.000945 |

| $\langle\langle P(cd\mid po)\rangle_N\rangle_{po}$ | SD | SD% | x | Σx |
|---|---|---|---|---|
| 0.000468 | 0.000107 | 22.800019 | 0.013297 | 0.043678 |

Pathogenic cd → c4 QFD 4

| $\langle P(cd\mid po)\rangle_N$ | SD | po | W(Da/Si) 25 sol | $\langle P(cd\mid af)\rangle_N$ |
|---|---|---|---|---|
| 0.001000 | 0.000330 | ExA | 450 | 0.000118 |
| 0.000980 | 0.000327 | MAL | 250 | 0.000066 |
| 0.000965 | 0.000317 | FEM | 250 | 0.000066 |
| 0.000951 | 0.000366 | EAS | 250 | 0.000066 |
| 0.000950 | 0.000353 | OTH | 1225 | 0.000321 |
| 0.000949 | 0.000342 | AMR | 250 | 0.000066 |
| 0.000949 | 0.000344 | AFR | 250 | 0.000066 |
| 0.000946 | 0.000352 | FIN | 250 | 0.000066 |
| 0.000945 | 0.000314 | NFE | 250 | 0.000066 |
| 0.000931 | 0.000347 | ASJ | 250 | 0.000066 |
| 0.000927 | 0.000361 | EUR | 75 | 0.000020 |
| 0.000803 | 0.000345 | SAS | 75 | 0.000020 |

| $\langle\langle P(cd\mid po)\rangle_N\rangle_{po}$ | SD | SD% | x | Σx |
|---|---|---|---|---|
| 0.000941 | 0.000048 | 5.097858 | 0.024666 | 0.040315 |

Pathogenic cd → c5

| $\langle P(cd\mid po)\rangle_N$ | SD | po | W(Da/Si) 25 sol | $\langle P(cd\mid af)\rangle_N$ |
|---|---|---|---|---|
| 0.001260 | 0.000086 | ExA | 550 | 0.000144 |
| 0.001244 | 0.000082 | OTH | 1025 | 0.000269 |
| 0.001235 | 0.000095 | MAL | 375 | 0.000098 |
| 0.001223 | 0.000111 | FEM | 375 | 0.000098 |
| 0.001223 | 0.000100 | EAS | 375 | 0.000098 |
| 0.001218 | 0.000115 | AFR | 375 | 0.000098 |
| 0.001215 | 0.000107 | ASJ | 375 | 0.000098 |
| 0.001214 | 0.000115 | FIN | 375 | 0.000098 |
| 0.001213 | 0.000115 | AMR | 375 | 0.000098 |
| 0.001201 | 0.000124 | NFE | 375 | 0.000098 |
| 0.001197 | 0.000113 | EUR | 50 | 0.000013 |
| 0.001109 | 0.000152 | SAS | 50 | 0.000013 |

| $\langle\langle P(cd\mid po)\rangle_N\rangle_{po}$ | SD | SD% | x | Σx |
|---|---|---|---|---|
| 0.001213 | 0.000037 | 3.053087 | 0.031781 | 0.040312 |

Pathogenic cd → c8 QFD 5

| $\langle P(cd\mid po)\rangle_N$ | SD | po | W(Da/Si) 25 sol | $\langle P(cd\mid af)\rangle_N$ |
|---|---|---|---|---|
| 0.001132 | 0.000161 | ExA | 1075 | 0.000282 |
| 0.001103 | 0.000153 | OTH | 1525 | 0.000400 |
| 0.001064 | 0.000150 | MAL | 475 | 0.000124 |
| 0.001040 | 0.000150 | FEM | 475 | 0.000124 |
| 0.001037 | 0.000176 | EAS | 575 | 0.001083 |
| 0.001037 | 0.000177 | AMR | 575 | 0.001083 |
| 0.001022 | 0.000154 | FIN | 475 | 0.000124 |
| 0.001017 | 0.000148 | EUR | 275 | 0.000072 |
| 0.001013 | 0.000162 | AFR | 575 | 0.000151 |
| 0.001012 | 0.000160 | ASJ | 475 | 0.000124 |
| 0.000995 | 0.000188 | NFE | 475 | 0.000124 |
| 0.000868 | 0.000163 | SAS | 275 | 0.000072 |
| 0.000713 | 0.000121 | YRI | 50 | 0.000945 |

| $\langle\langle P(cd\mid po)\rangle_N\rangle_{po}$ | SD | SD% | x | Σx |
|---|---|---|---|---|
| 0.001004 | 0.000107 | 10.650413 | 0.028503 | 0.043678 |

Pathogenic cd → cx QFD 6

| $\langle P(cd\mid po)\rangle_N$ | SD | po | W(Da/Si) 25 sol | $\langle P(cd\mid af)\rangle_N$ |
|---|---|---|---|---|
| 0.001140 | 0.000170 | ExA | 775 | 0.000203 |
| 0.001116 | 0.000180 | OTH | 1925 | 0.000505 |
| 0.001099 | 0.000215 | MAL | 450 | 0.000118 |
| 0.001097 | 0.000219 | EAS | 475 | 0.000124 |
| 0.001077 | 0.000212 | FEM | 450 | 0.000118 |
| 0.001075 | 0.000223 | AMR | 475 | 0.000124 |
| 0.001073 | 0.000224 | FIN | 450 | 0.000118 |
| 0.001072 | 0.000203 | AFR | 475 | 0.000124 |
| 0.001067 | 0.000244 | NFE | 450 | 0.000118 |
| 0.001064 | 0.000221 | ASJ | 450 | 0.000118 |
| 0.001050 | 0.000230 | EUR | 125 | 0.000033 |
| 0.000957 | 0.000290 | SAS | 125 | 0.000033 |

| $\langle\langle P(cd\mid po)\rangle_N\rangle_{po}$ | SD | SD% | x | Σx |
|---|---|---|---|---|
| 0.001074 | 0.000045 | 4.153200 | 0.028145 | 0.040315 |

Pathogenic cd → f6

| $\langle P(cd\mid po)\rangle_N$ | SD | po | W(Da/Si) 25 sol | $\langle P(cd\mid af)\rangle_N$ |
|---|---|---|---|---|
| 0.001374 | 0.000066 | ExA | 1500 | 0.000393 |
| 0.001369 | 0.000074 | OTH | 2450 | 0.000642 |
| 0.001347 | 0.000086 | MAL | 600 | 0.000157 |
| 0.001344 | 0.000090 | EAS | 675 | 0.000177 |
| 0.001340 | 0.000079 | FEM | 600 | 0.000157 |
| 0.001336 | 0.000083 | AMR | 675 | 0.000177 |
| 0.001333 | 0.000081 | AFR | 675 | 0.001109 |
| 0.001326 | 0.000089 | ASJ | 600 | 0.000157 |
| 0.001323 | 0.000095 | FIN | 600 | 0.000157 |
| 0.001314 | 0.000099 | NFE | 600 | 0.000157 |
| 0.001313 | 0.000106 | EUR | 175 | 0.000046 |
| 0.001210 | 0.000178 | SAS | 175 | 0.000046 |

| $\langle\langle P(cd\mid po)\rangle_N\rangle_{po}$ | SD | SD% | x | Σx |
|---|---|---|---|---|
| 0.001327 | 0.000041 | 3.125636 | 0.034786 | 0.040318 |

Pathogenic cd → f7 QFD 7

| $\langle P(cd\mid po)\rangle_N$ | SD | po | W(Da/Si) 25 sol | $\langle P(cd\mid af)\rangle_N$ |
|---|---|---|---|---|
| 0.000552 | 0.000245 | ExA | 975 | 0.001187 |
| 0.000546 | 0.000175 | OTH | 1525 | 0.001332 |
| 0.000464 | 0.000198 | MAL | 550 | 0.000144 |
| 0.000436 | 0.000201 | AMR | 600 | 0.003982 |
| 0.000431 | 0.000204 | EAS | 600 | 0.001089 |
| 0.000428 | 0.000189 | FEM | 550 | 0.000144 |
| 0.000426 | 0.000236 | NFE | 550 | 0.001076 |
| 0.000406 | 0.000199 | FIN | 550 | 0.001076 |
| 0.000403 | 0.000191 | ASJ | 550 | 0.000144 |
| 0.000391 | 0.000192 | AFR | 600 | 0.001089 |
| 0.000374 | 0.000124 | EUR | 300 | 0.001010 |
| 0.000276 | 0.000126 | SAS | 300 | 0.000079 |

| $\langle\langle P(cd\mid po)\rangle_N\rangle_{po}$ | SD | SD% | x | Σx |
|---|---|---|---|---|
| 0.000428 | 0.000073 | 17.166463 | 0.011209 | 0.040310 |

Pathogenic cd → f9 QFD 8

| $\langle P(cd\mid po)\rangle_N$ | SD | po | W(Da/Si) 25 sol | $\langle P(cd\mid af)\rangle_N$ |
|---|---|---|---|---|
| 0.001219 | 0.000127 | ExA | 950 | 0.000249 |
| 0.001190 | 0.000126 | OTH | 1775 | 0.000465 |
| 0.001168 | 0.000126 | MAL | 525 | 0.000138 |
| 0.001145 | 0.000162 | EAS | 700 | 0.000183 |
| 0.001128 | 0.000155 | FIN | 525 | 0.000138 |
| 0.001128 | 0.000168 | FEM | 525 | 0.000138 |
| 0.001123 | 0.000151 | ASJ | 525 | 0.000138 |
| 0.001123 | 0.000153 | AMR | 700 | 0.000183 |
| 0.001121 | 0.000151 | AFR | 700 | 0.000183 |
| 0.001117 | 0.000132 | EUR | 250 | 0.000066 |
| 0.001113 | 0.000153 | NFE | 525 | 0.000138 |
| 0.001008 | 0.000173 | SAS | 250 | 0.000066 |

| $\langle\langle P(cd\mid po)\rangle_N\rangle_{po}$ | SD | SD% | x | Σx |
|---|---|---|---|---|
| 0.001132 | 0.000051 | 4.518160 | 0.029661 | 0.040315 |

Pathogenic cd → l2 QFD 9

| $\langle P(cd\mid po)\rangle_N$ | SD | po | W(Da/Si) 25 sol | $\langle P(cd\mid af)\rangle_N$ |
|---|---|---|---|---|
| 0.000565 | 0.000283 | ExA | 600 | 0.000157 |
| 0.000528 | 0.000271 | OTH | 1175 | 0.001240 |
| 0.000481 | 0.000252 | MAL | 350 | 0.000092 |
| 0.000460 | 0.000243 | EAS | 450 | 0.000118 |
| 0.000455 | 0.000246 | FEM | 350 | 0.000092 |
| 0.000450 | 0.000249 | ASJ | 350 | 0.000092 |
| 0.000444 | 0.000236 | AMR | 450 | 0.000118 |
| 0.000441 | 0.000221 | FIN | 350 | 0.001024 |
| 0.000432 | 0.000228 | NFE | 350 | 0.000092 |
| 0.000422 | 0.000248 | AFR | 450 | 0.000118 |
| 0.000413 | 0.000199 | EUR | 200 | 0.000052 |
| 0.000315 | 0.000191 | SAS | 200 | 0.000052 |

| $\langle\langle P(cd\mid po)\rangle_N\rangle_{po}$ | SD | SD% | x | Σx |
|---|---|---|---|---|
| 0.000450 | 0.000061 | 13.545600 | 0.011802 | 0.040318 |

Pathogenic cd → l5 QFD 10

| $\langle P(cd\mid po)\rangle_N$ | SD | po | W(Da/Si) 25 sol | $\langle P(cd\mid af)\rangle_N$ |
|---|---|---|---|---|
| 0.001029 | 0.000164 | ExA | 75 | 0.000020 |
| 0.001012 | 0.000141 | OTH | 150 | 0.000039 |
| 0.000971 | 0.000157 | MAL | 75 | 0.000020 |
| 0.000937 | 0.000161 | FEM | 75 | 0.000020 |
| 0.000935 | 0.000161 | EAS | 75 | 0.000020 |
| 0.000922 | 0.000169 | NFE | 75 | 0.000020 |
| 0.000921 | 0.000144 | FIN | 75 | 0.000020 |
| 0.000915 | 0.000143 | AMR | 75 | 0.000020 |
| 0.000912 | 0.000155 | ASJ | 75 | 0.000020 |
| 0.000910 | 0.000163 | AFR | 75 | 0.000020 |

| $\langle\langle P(cd\mid po)\rangle_N\rangle_{po}$ | SD | SD% | x | Σx |
|---|---|---|---|---|
| 0.000946 | 0.000043 | 4.563071 | 0.020666 | 0.033590 |

Pathogenic cd → l6

| $\langle P(cd\mid po)\rangle_N$ | SD | po | W(Da/Si) 25 sol | $\langle P(cd\mid af)\rangle_N$ |
|---|---|---|---|---|
| 0.001387 | 0.000064 | ExA | 425 | 0.000111 |
| 0.001376 | 0.000072 | EAS | 275 | 0.000072 |
| 0.001373 | 0.000054 | AMR | 275 | 0.000072 |
| 0.001367 | 0.000067 | OTH | 500 | 0.000131 |
| 0.001367 | 0.000069 | MAL | 275 | 0.000072 |
| 0.001365 | 0.000062 | FEM | 275 | 0.000072 |
| 0.001364 | 0.000068 | FIN | 275 | 0.000072 |
| 0.001359 | 0.000071 | ASJ | 275 | 0.000072 |
| 0.001358 | 0.000073 | NFE | 275 | 0.000072 |
| 0.001356 | 0.000076 | AFR | 275 | 0.000072 |
| 0.001353 | 0.000069 | EUR | 75 | 0.000020 |
| 0.001298 | 0.000106 | SAS | 75 | 0.000020 |

| $\langle\langle P(cd\mid po)\rangle_N\rangle_{po}$ | SD | SD% | x | Σx |
|---|---|---|---|---|
| 0.001360 | 0.000022 | 1.608230 | 0.035645 | 0.040318 |

Pathogenic cd → l8 QFD 11

| $\langle P(cd\mid po)\rangle_N$ | SD | po | W(Da/Si) 25 sol | $\langle P(cd\mid af)\rangle_N$ |
|---|---|---|---|---|
| 0.001115 | 0.000145 | ExA | 50 | 0.000013 |
| 0.001076 | 0.000144 | OTH | 25 | 0.000007 |
| 0.001054 | 0.000147 | MAL | 25 | 0.000007 |
| 0.001027 | 0.000148 | EAS | 25 | 0.000007 |
| 0.001026 | 0.000147 | FEM | 25 | 0.000007 |
| 0.001017 | 0.000161 | AMR | 25 | 0.000007 |
| 0.001012 | 0.000155 | FIN | 25 | 0.000007 |
| 0.001006 | 0.000164 | ASJ | 25 | 0.000007 |
| 0.001005 | 0.000168 | AFR | 25 | 0.000007 |
| 0.000996 | 0.000170 | NFE | 25 | 0.000007 |

| $\langle\langle P(cd\mid po)\rangle_N\rangle_{po}$ | SD | SD% | x | Σx |
|---|---|---|---|---|
| 0.001033 | 0.000038 | 3.641732 | 0.022567 | 0.033598 |

Pathogenic cd → lx QFD 12

| $\langle P(cd\mid po)\rangle_N$ | SD | po | W(Da/Si) 25 sol | $\langle P(cd\mid af)\rangle_N$ |
|---|---|---|---|---|
| 0.001006 | 0.000152 | ExA | 75 | 0.000020 |
| 0.000995 | 0.000175 | OTH | 200 | 0.000052 |
| 0.000942 | 0.000151 | MAL | 100 | 0.000026 |
| 0.000912 | 0.000173 | EAS | 100 | 0.000026 |
| 0.000903 | 0.000150 | FEM | 100 | 0.000026 |
| 0.000903 | 0.000142 | FIN | 100 | 0.000026 |
| 0.000892 | 0.000183 | NFE | 100 | 0.000026 |
| 0.000886 | 0.000152 | ASJ | 100 | 0.000026 |
| 0.000882 | 0.000150 | AMR | 100 | 0.000026 |
| 0.000878 | 0.000159 | AFR | 100 | 0.000026 |

| $\langle\langle P(cd\mid po)\rangle_N\rangle_{po}$ | SD | SD% | x | Σx |
|---|---|---|---|---|
| 0.000920 | 0.000046 | 5.052821 | 0.020089 | 0.033596 |

Pathogenic cd → pr

| $\langle P(cd\mid po)\rangle_N$ | SD | po | W(Da/Si) 25 sol | $\langle P(cd\mid af)\rangle_N$ |
|---|---|---|---|---|
| 0.001240 | 0.000102 | ExA | 375 | 0.000098 |
| 0.001209 | 0.000105 | AMR | 350 | 0.000092 |
| 0.001206 | 0.000148 | MAL | 350 | 0.000092 |
| 0.001206 | 0.000123 | OTH | 1025 | 0.000269 |
| 0.001190 | 0.000133 | FEM | 350 | 0.000092 |
| 0.001185 | 0.000157 | EAS | 350 | 0.000092 |
| 0.001178 | 0.000147 | ASJ | 350 | 0.000092 |
| 0.001174 | 0.000126 | EUR | 100 | 0.000026 |
| 0.001171 | 0.000149 | AFR | 350 | 0.000092 |
| 0.001167 | 0.000142 | FIN | 350 | 0.000092 |
| 0.001159 | 0.000138 | NFE | 350 | 0.000092 |
| 0.001055 | 0.000200 | SAS | 100 | 0.000026 |

| $\langle\langle P(cd\mid po)\rangle_N\rangle_{po}$ | SD | SD% | x | Σx |
|---|---|---|---|---|
| 0.001178 | 0.000045 | 3.806742 | 0.030876 | 0.040318 |

Pathogenic cd → s1 QFD 13

| $\langle P(cd\mid po)\rangle_N$ | SD | po | W(Da/Si) 25 sol | $\langle P(cd\mid af)\rangle_N$ |
|---|---|---|---|---|
| 0.001050 | 0.000159 | ExA | 150 | 0.000039 |
| 0.001026 | 0.000173 | OTH | 100 | 0.000026 |
| 0.000964 | 0.000172 | MAL | 25 | 0.000007 |
| 0.000953 | 0.000177 | EAS | 100 | 0.000026 |
| 0.000936 | 0.000145 | FEM | 25 | 0.000007 |
| 0.000929 | 0.000158 | AMR | 100 | 0.000026 |
| 0.000927 | 0.000159 | FIN | 25 | 0.000007 |
| 0.000917 | 0.000190 | NFE | 25 | 0.000007 |
| 0.000914 | 0.000139 | ASJ | 25 | 0.000007 |
| 0.000912 | 0.000161 | AFR | 100 | 0.000026 |
| 0.000905 | 0.000142 | EUR | 75 | 0.000020 |
| 0.000803 | 0.000152 | SAS | 75 | 0.000020 |

| $\langle\langle P(cd\mid po)\rangle_N\rangle_{po}$ | SD | SD% | x | Σx |
|---|---|---|---|---|
| 0.000936 | 0.000062 | 6.638607 | 0.024535 | 0.040318 |

Pathogenic cd → s2 QFD 14

| $\langle P(cd\mid po)\rangle_N$ | SD | po | W(Da/Si) 25 sol | $\langle P(cd\mid af)\rangle_N$ |
|---|---|---|---|---|
| 0.001027 | 0.000186 | ExA | 175 | 0.000046 |
| 0.000989 | 0.000163 | OTH | 150 | 0.000039 |
| 0.000941 | 0.000167 | MAL | 100 | 0.000026 |
| 0.000922 | 0.000171 | FEM | 100 | 0.000026 |
| 0.000915 | 0.000156 | EAS | 125 | 0.000033 |
| 0.000912 | 0.000164 | FIN | 100 | 0.000026 |
| 0.000905 | 0.000183 | AMR | 125 | 0.000033 |
| 0.000904 | 0.000164 | ASJ | 100 | 0.000026 |
| 0.000901 | 0.000186 | NFE | 100 | 0.000026 |
| 0.000886 | 0.000175 | AFR | 125 | 0.000965 |
| 0.000877 | 0.000136 | EUR | 50 | 0.000013 |
| 0.000771 | 0.000163 | SAS | 50 | 0.000013 |

| $\langle\langle P(cd\mid po)\rangle_N\rangle_{po}$ | SD | SD% | x | Σx |
|---|---|---|---|---|
| 0.000912 | 0.000062 | 6.789107 | 0.023911 | 0.040318 |

Pathogenic cd → s3 QFD 15

| $\langle P(cd\mid po)\rangle_N$ | SD | po | W(Da/Si) 25 sol | $\langle P(cd\mid af)\rangle_N$ |
|---|---|---|---|---|
| 0.001074 | 0.000176 | ExA | 125 | 0.000033 |
| 0.001065 | 0.000138 | OTH | 200 | 0.000052 |
| 0.000990 | 0.000164 | MAL | 50 | 0.000013 |
| 0.000972 | 0.000181 | AMR | 50 | 0.000013 |
| 0.000964 | 0.000175 | FEM | 50 | 0.000013 |
| 0.000964 | 0.000175 | EAS | 50 | 0.000013 |
| 0.000959 | 0.000163 | FIN | 50 | 0.000013 |
| 0.000955 | 0.000178 | NFE | 50 | 0.000013 |
| 0.000947 | 0.000155 | ASJ | 50 | 0.000013 |
| 0.000942 | 0.000175 | AFR | 50 | 0.000013 |

| $\langle\langle P(cd\mid po)\rangle_N\rangle_{po}$ | SD | SD% | x | Σx |
|---|---|---|---|---|
| 0.000983 | 0.000047 | 4.814980 | 0.021472 | 0.033598 |

Pathogenic cd → s4 QFD 16

| $\langle P(cd\mid po)\rangle_N$ | SD | po | W(Da/Si) 25 sol | $\langle P(cd\mid af)\rangle_N$ |
|---|---|---|---|---|
| 0.001021 | 0.000150 | OTH | 100 | 0.000026 |
| 0.001012 | 0.000183 | ExA | 50 | 0.000013 |
| 0.000949 | 0.000176 | MAL | 25 | 0.000007 |
| 0.000923 | 0.000174 | EAS | 25 | 0.000007 |
| 0.000922 | 0.000183 | AMR | 25 | 0.000007 |
| 0.000919 | 0.000188 | FEM | 25 | 0.000007 |
| 0.000908 | 0.000157 | ASJ | 25 | 0.000007 |
| 0.000908 | 0.000173 | FIN | 25 | 0.000007 |
| 0.000904 | 0.000174 | NFE | 25 | 0.000007 |
| 0.000896 | 0.000171 | AFR | 25 | 0.000007 |

| $\langle\langle P(cd\mid po)\rangle_N\rangle_{po}$ | SD | SD% | x | Σx |
|---|---|---|---|---|
| 0.000936 | 0.000045 | 4.773704 | 0.020442 | 0.033598 |

Pathogenic cd → s5

| $\langle P(cd\mid po)\rangle_N$ | SD | po | W(Da/Si) 25 sol | $\langle P(cd\mid af)\rangle_N$ |
|---|---|---|---|---|
| 0.001139 | 0.000124 | OTH | 175 | 0.000046 |
| 0.001077 | 0.000154 | EAS | 75 | 0.000020 |
| 0.001074 | 0.000134 | MAL | 75 | 0.000020 |
| 0.001049 | 0.000153 | FEM | 75 | 0.000020 |
| 0.001044 | 0.000167 | AMR | 75 | 0.000020 |
| 0.001042 | 0.000134 | FIN | 75 | 0.000020 |
| 0.001041 | 0.000153 | ASJ | 75 | 0.000020 |
| 0.001030 | 0.000156 | NFE | 75 | 0.000020 |
| 0.001027 | 0.000157 | AFR | 75 | 0.000020 |

| $\langle\langle P(cd\mid po)\rangle_N\rangle_{po}$ | SD | SD% | x | Σx |
|---|---|---|---|---|
| 0.001058 | 0.000035 | 3.309930 | 0.020795 | 0.030239 |

Pathogenic cd → s6 QFD 17

| $\langle P(cd\mid po)\rangle_N$ | SD | po | W(Da/Si) 25 sol | $\langle P(cd\mid af)\rangle_N$ |
|---|---|---|---|---|
| 0.001049 | 0.000153 | ExA | 75 | 0.000020 |
| 0.001011 | 0.000163 | OTH | 150 | 0.000039 |
| 0.000960 | 0.000152 | MAL | 75 | 0.000020 |
| 0.000942 | 0.000182 | EAS | 75 | 0.000020 |
| 0.000931 | 0.000165 | FEM | 75 | 0.000020 |
| 0.000927 | 0.000182 | AMR | 75 | 0.000020 |
| 0.000923 | 0.000171 | FIN | 75 | 0.000020 |
| 0.000915 | 0.000169 | ASJ | 75 | 0.000020 |
| 0.000915 | 0.000197 | NFE | 75 | 0.000020 |
| 0.000912 | 0.000155 | EUR | 25 | 0.000007 |
| 0.000908 | 0.000157 | AFR | 75 | 0.000020 |
| 0.000786 | 0.000172 | SAS | 25 | 0.000007 |

| $\langle\langle P(cd\mid po)\rangle_N\rangle_{po}$ | SD | SD% | x | Σx |
|---|---|---|---|---|
| 0.000932 | 0.000063 | 6.766703 | 0.024412 | 0.040315 |

Pathogenic cd → s7 QFD 18

| $\langle P(cd\mid po)\rangle_N$ | SD | po | W(Da/Si) 25 sol | $\langle P(cd\mid af)\rangle_N$ |
|---|---|---|---|---|
| 0.001035 | 0.000170 | ExA | 25 | 0.000007 |
| 0.001005 | 0.000155 | OTH | 125 | 0.000033 |
| 0.000959 | 0.000165 | MAL | 25 | 0.000007 |
| 0.000935 | 0.000153 | EAS | 25 | 0.000007 |
| 0.000932 | 0.000179 | FEM | 25 | 0.000007 |
| 0.000913 | 0.000165 | AMR | 25 | 0.000007 |
| 0.000912 | 0.000136 | FIN | 25 | 0.000007 |
| 0.000910 | 0.000165 | ASJ | 25 | 0.000007 |
| 0.000908 | 0.000174 | AFR | 25 | 0.000007 |
| 0.000905 | 0.000146 | NFE | 25 | 0.000007 |

| $\langle\langle P(cd\mid po)\rangle_N\rangle_{po}$ | SD | SD% | x | Σx |
|---|---|---|---|---|
| 0.000941 | 0.000045 | 4.791049 | 0.020555 | 0.033582 |

Pathogenic cd → s8 QFD 19

| $\langle P(cd\mid po)\rangle_N$ | SD | po | W(Da/Si) 25 sol | $\langle P(cd\mid af)\rangle_N$ |
|---|---|---|---|---|
| 0.001133 | 0.000153 | ExA | 275 | 0.000072 |
| 0.001098 | 0.000175 | OTH | 250 | 0.000066 |
| 0.001067 | 0.000167 | MAL | 100 | 0.000026 |
| 0.001046 | 0.000174 | FEM | 100 | 0.000026 |
| 0.001036 | 0.000198 | EAS | 100 | 0.000026 |
| 0.001035 | 0.000160 | AMR | 100 | 0.000026 |
| 0.001026 | 0.000163 | FIN | 100 | 0.000026 |
| 0.001026 | 0.000183 | AFR | 100 | 0.000026 |
| 0.001021 | 0.000195 | ASJ | 100 | 0.000026 |
| 0.000994 | 0.000233 | NFE | 100 | 0.000026 |

| $\langle\langle P(cd\mid po)\rangle_N\rangle_{po}$ | SD | SD% | x | Σx |
|---|---|---|---|---|
| 0.001048 | 0.000041 | 3.918583 | 0.022886 | 0.033598 |

Pathogenic cd → z1 QFD 20

| $\langle P(cd\mid po)\rangle_N$ | SD | po | W(Da/Si) 25 sol | $\langle P(cd\mid af)\rangle_N$ |
|---|---|---|---|---|
| 0.000990 | 0.000148 | ExA | 50 | 0.000013 |
| 0.000974 | 0.000142 | OTH | 25 | 0.000007 |
| 0.000928 | 0.000133 | MAL | 25 | 0.000007 |
| 0.000905 | 0.000144 | FEM | 25 | 0.000007 |
| 0.000900 | 0.000139 | FIN | 25 | 0.000007 |
| 0.000895 | 0.000152 | EAS | 25 | 0.000007 |
| 0.000881 | 0.000151 | NFE | 25 | 0.000007 |
| 0.000874 | 0.000159 | AMR | 25 | 0.000007 |
| 0.000871 | 0.000163 | ASJ | 25 | 0.000007 |
| 0.000863 | 0.000138 | AFR | 25 | 0.000007 |

| $\langle\langle P(cd\mid po)\rangle_N\rangle_{po}$ | SD | SD% | x | Σx |
|---|---|---|---|---|
| 0.000908 | 0.000043 | 4.781024 | 0.019829 | 0.033598 |

{MYBPC3 R-correlation range over domains}

{MYBPC3 Benign Summary}

Benign cd → c0 QFD 1

| $\langle P(cd\mid po)\rangle_N$ | SD | po | W(Da/Si) 25 sol | $\langle P(cd\mid af)\rangle_N$ |
|---|---|---|---|---|
| 0.000864 | 0.000164 | SAS | 250 | 0.000066 |
| 0.000756 | 0.000137 | EUR | 250 | 0.000066 |
| 0.000742 | 0.000186 | NFE | 650 | 0.000170 |
| 0.000740 | 0.000176 | AFR | 800 | 0.000210 |
| 0.000736 | 0.000175 | ASJ | 650 | 0.000170 |
| 0.000734 | 0.000180 | AMR | 800 | 0.000210 |
| 0.000731 | 0.000184 | FEM | 650 | 0.000170 |
| 0.000722 | 0.000161 | EAS | 800 | 0.000210 |
| 0.000720 | 0.000167 | FIN | 650 | 0.000170 |
| 0.000705 | 0.000167 | MAL | 650 | 0.000170 |
| 0.000651 | 0.000170 | OTH | 1675 | 0.000439 |
| 0.000635 | 0.000194 | ExA | 925 | 0.000242 |

| $\langle\langle P(cd\mid po)\rangle_N\rangle_{po}$ | SD | SD% | x | R |
|---|---|---|---|---|
| 0.000728 | 0.000056 | 7.734975 | 0.019081 | −0.865990 |

Benign cd → c1 QFD 2

| $\langle P(cd\mid po)\rangle_N$ | SD | po | W(Da/Si) 25 sol | $\langle P(cd\mid af)\rangle_N$ |
|---|---|---|---|---|
| 0.000617 | 0.000165 | YRI | 25 | 0.000938 |
| 0.000435 | 0.000146 | SAS | 325 | 0.004842 |
| 0.000336 | 0.000117 | ASJ | 700 | 0.007833 |
| 0.000336 | 0.000127 | NFE | 700 | 0.007833 |
| 0.000336 | 0.000107 | EUR | 325 | 0.007735 |
| 0.000331 | 0.000130 | FEM | 700 | 0.007833 |
| 0.000331 | 0.000135 | EAS | 750 | 0.001128 |
| 0.000329 | 0.000128 | FIN | 700 | 0.007833 |
| 0.000327 | 0.000117 | AMR | 750 | 0.004953 |
| 0.000322 | 0.000090 | CEP | 50 | 0.007662 |
| 0.000321 | 0.000098 | AFR | 750 | 0.002992 |
| 0.000312 | 0.000108 | MAL | 700 | 0.007833 |
| 0.000274 | 0.000075 | OTH | 2025 | 0.012937 |
| 0.000271 | 0.000086 | ExA | 1175 | 0.007957 |

| $\langle\langle P(cd\mid po)\rangle_N\rangle_{po}$ | SD | SD% | x | R |
|---|---|---|---|---|
| 0.000348 | 0.000086 | 24.644263 | 0.010647 | −0.715106 |

Benign cd → c2

| $\langle P(cd\mid po)\rangle_N$ | SD | po | W(Da/Si) 25 sol | $\langle P(cd\mid af)\rangle_N$ |
|---|---|---|---|---|
| 0.000487 | 0.000394 | YRI | 25 | 0.003831 |
| 0.000380 | 0.000315 | SAS | 125 | 0.000033 |
| 0.000312 | 0.000289 | NFE | 350 | 0.000092 |
| 0.000303 | 0.000281 | FIN | 350 | 0.000092 |
| 0.000300 | 0.000281 | AFR | 425 | 0.001043 |
| 0.000296 | 0.000282 | ASJ | 350 | 0.000092 |
| 0.000296 | 0.000257 | EUR | 125 | 0.000033 |
| 0.000288 | 0.000273 | FEM | 350 | 0.001024 |
| 0.000278 | 0.000255 | MAL | 350 | 0.001024 |
| 0.000273 | 0.000266 | EAS | 425 | 0.000111 |
| 0.000241 | 0.000222 | OTH | 1200 | 0.000315 |

| $\langle\langle P(cd\mid po)\rangle_N\rangle_{po}$ | SD | SD% | x | R |
|---|---|---|---|---|
| 0.000314 | 0.000066 | 21.174111 | 0.007543 | −0.812095 |

Benign cd → c3 QFD 3

| $\langle P(cd\mid po)\rangle_N$ | SD | po | W(Da/Si) 25 sol | $\langle P(cd\mid af)\rangle_N$ |
|---|---|---|---|---|
| 0.001343 | 0.000077 | YRI | 50 | 0.000945 |
| 0.001198 | 0.000162 | SAS | 150 | 0.000039 |
| 0.001101 | 0.000172 | EUR | 150 | 0.000039 |
| 0.001074 | 0.000169 | AFR | 550 | 0.000144 |
| 0.001073 | 0.000195 | ASJ | 550 | 0.000144 |
| 0.001060 | 0.000168 | FIN | 550 | 0.000144 |
| 0.001059 | 0.000182 | NFE | 550 | 0.000144 |
| 0.001046 | 0.000200 | AMR | 550 | 0.000144 |
| 0.001042 | 0.000233 | EAS | 550 | 0.000144 |
| 0.001039 | 0.000192 | FEM | 550 | 0.000144 |
| 0.001021 | 0.000207 | MAL | 550 | 0.000144 |
| 0.000939 | 0.000188 | OTH | 1325 | 0.000347 |
| 0.000918 | 0.000211 | ExA | 1100 | 0.000288 |

| $\langle\langle P(cd\mid po)\rangle_N\rangle_{po}$ | SD | SD% | x | R |
|---|---|---|---|---|
| 0.001070 | 0.000107 | 9.976527 | 0.030381 | −0.870811 |

Benign cd → c4 QFD 4

| $\langle P(cd\mid po)\rangle_N$ | SD | po | W(Da/Si) 25 sol | $\langle P(cd\mid af)\rangle_N$ |
|---|---|---|---|---|
| 0.000736 | 0.000344 | SAS | 75 | 0.000020 |
| 0.000612 | 0.000361 | EUR | 75 | 0.000020 |
| 0.000606 | 0.000348 | ASJ | 250 | 0.000066 |
| 0.000594 | 0.000314 | NFE | 250 | 0.000066 |
| 0.000592 | 0.000352 | FIN | 250 | 0.000066 |
| 0.000590 | 0.000342 | AMR | 250 | 0.000066 |
| 0.000590 | 0.000344 | AFR | 250 | 0.000066 |
| 0.000589 | 0.000353 | OTH | 1225 | 0.000321 |
| 0.000587 | 0.000366 | EAS | 250 | 0.000066 |
| 0.000573 | 0.000317 | FEM | 250 | 0.000066 |
| 0.000559 | 0.000327 | MAL | 250 | 0.000066 |
| 0.000538 | 0.000330 | ExA | 450 | 0.000118 |

| $\langle\langle P(cd\mid po)\rangle_N\rangle_{po}$ | SD | SD% | x | R |
|---|---|---|---|---|
| 0.000597 | 0.000048 | 8.038947 | 0.015649 | −0.769593 |

Benign cd → c5

| $\langle P(cd\mid po)\rangle_N$ | SD | po | W(Da/Si) 25 sol | $\langle P(cd\mid af)\rangle_N$ |
|---|---|---|---|---|
| 0.000430 | 0.000152 | SAS | 50 | 0.000013 |
| 0.000341 | 0.000113 | EUR | 50 | 0.000013 |
| 0.000337 | 0.000123 | NFE | 375 | 0.000098 |
| 0.000326 | 0.000115 | AMR | 375 | 0.000098 |
| 0.000324 | 0.000115 | FIN | 375 | 0.000098 |
| 0.000323 | 0.000107 | ASJ | 375 | 0.000098 |
| 0.000321 | 0.000115 | AFR | 375 | 0.000098 |
| 0.000315 | 0.000111 | FEM | 375 | 0.000098 |
| 0.000315 | 0.000100 | EAS | 375 | 0.000098 |
| 0.000304 | 0.000095 | MAL | 375 | 0.000098 |
| 0.000293 | 0.000079 | OTH | 1025 | 0.000269 |
| 0.000278 | 0.000086 | ExA | 550 | 0.000144 |

| $\langle\langle P(cd\mid po)\rangle_N\rangle_{po}$ | SD | SD% | x | R |
|---|---|---|---|---|
| 0.000326 | 0.000037 | 11.431583 | 0.008531 | −0.818275 |

Benign cd → c8 QFD 5

| $\langle P(cd\mid po)\rangle_N$ | SD | po | W(Da/Si) 25 sol | $\langle P(cd\mid af)\rangle_N$ |
|---|---|---|---|---|
| 0.000826 | 0.000120 | YRI | 50 | 0.000945 |
| 0.000671 | 0.000163 | SAS | 275 | 0.000072 |
| 0.000544 | 0.000188 | NFE | 475 | 0.000124 |
| 0.000527 | 0.000160 | ASJ | 475 | 0.000124 |
| 0.000526 | 0.000162 | AFR | 575 | 0.000151 |
| 0.000522 | 0.000148 | EUR | 275 | 0.000072 |
| 0.000517 | 0.000154 | FIN | 475 | 0.000124 |
| 0.000501 | 0.000176 | EAS | 575 | 0.001083 |
| 0.000501 | 0.000177 | AMR | 575 | 0.001083 |
| 0.000499 | 0.000150 | FEM | 475 | 0.000124 |
| 0.000474 | 0.000150 | MAL | 475 | 0.000124 |
| 0.000436 | 0.000154 | OTH | 1525 | 0.000400 |
| 0.000406 | 0.000161 | ExA | 1075 | 0.000282 |

| $\langle\langle P(cd\mid po)\rangle_N\rangle_{po}$ | SD | SD% | x | R |
|---|---|---|---|---|
| 0.000535 | 0.000107 | 20.004932 | 0.015175 | −0.824628 |

Benign cd → cx QFD 6

| $\langle P(cd\mid po)\rangle_N$ | SD | po | W(Da/Si) 25 sol | $\langle P(cd\mid af)\rangle_N$ |
|---|---|---|---|---|
| 0.000582 | 0.000290 | SAS | 125 | 0.000033 |
| 0.000488 | 0.000230 | EUR | 125 | 0.000033 |
| 0.000474 | 0.000221 | ASJ | 450 | 0.000118 |
| 0.000471 | 0.000244 | NFE | 450 | 0.000118 |
| 0.000466 | 0.000203 | AFR | 475 | 0.000124 |
| 0.000465 | 0.000224 | FIN | 450 | 0.000118 |
| 0.000463 | 0.000223 | AMR | 475 | 0.000124 |
| 0.000462 | 0.000212 | FEM | 450 | 0.000118 |
| 0.000441 | 0.000219 | EAS | 475 | 0.000124 |
| 0.000440 | 0.000215 | MAL | 450 | 0.000118 |
| 0.000422 | 0.000180 | OTH | 1925 | 0.000505 |
| 0.000398 | 0.000170 | ExA | 775 | 0.000203 |

| $\langle\langle P(cd\mid po)\rangle_N\rangle_{po}$ | SD | SD% | x | R |
|---|---|---|---|---|
| 0.000464 | 0.000045 | 9.623709 | 0.012170 | −0.855669 |

Benign cd → f6

| $\langle P(cd\mid po)\rangle_N$ | SD | po | W(Da/Si) 25 sol | $\langle P(cd\mid af)\rangle_N$ |
|---|---|---|---|---|
| 0.000328 | 0.000178 | SAS | 175 | 0.000046 |
| 0.000226 | 0.000106 | EUR | 175 | 0.000046 |
| 0.000224 | 0.000099 | NFE | 600 | 0.000157 |
| 0.000215 | 0.000095 | FIN | 600 | 0.000157 |
| 0.000213 | 0.000089 | ASJ | 600 | 0.000157 |
| 0.000205 | 0.000081 | AFR | 675 | 0.001109 |
| 0.000203 | 0.000083 | AMR | 675 | 0.000177 |
| 0.000199 | 0.000079 | FEM | 600 | 0.000157 |
| 0.000195 | 0.000090 | EAS | 675 | 0.000177 |
| 0.000191 | 0.000086 | MAL | 600 | 0.000157 |
| 0.000171 | 0.000075 | OTH | 2450 | 0.000642 |
| 0.000164 | 0.000066 | ExA | 1500 | 0.000393 |

| $\langle\langle P(cd\mid po)\rangle_N\rangle_{po}$ | SD | SD% | x | R |
|---|---|---|---|---|
| 0.000211 | 0.000041 | 19.642693 | 0.005532 | −0.816704 |

Benign cd → f7 QFD 7

| $\langle P(cd\mid po)\rangle_N$ | SD | po | W(Da/Si) 25 sol | $\langle P(cd\mid af)\rangle_N$ |
|---|---|---|---|---|
| 0.001262 | 0.000127 | SAS | 300 | 0.000079 |
| 0.001164 | 0.000124 | EUR | 300 | 0.001010 |
| 0.001147 | 0.000192 | AFR | 600 | 0.001089 |
| 0.001136 | 0.000191 | ASJ | 550 | 0.000144 |
| 0.001132 | 0.000199 | FIN | 550 | 0.001076 |
| 0.001113 | 0.000235 | NFE | 550 | 0.001076 |
| 0.001110 | 0.000189 | FEM | 550 | 0.000144 |
| 0.001108 | 0.000204 | EAS | 600 | 0.001089 |
| 0.001103 | 0.000201 | AMR | 600 | 0.003982 |
| 0.001074 | 0.000198 | MAL | 550 | 0.000144 |
| 0.000990 | 0.000177 | OTH | 1525 | 0.001332 |
| 0.000987 | 0.000245 | ExA | 975 | 0.001187 |

| $\langle\langle P(cd\mid po)\rangle_N\rangle_{po}$ | SD | SD% | x | R |
|---|---|---|---|---|
| 0.001111 | 0.000073 | 6.616868 | 0.029100 | −0.916946 |

Benign cd → f9 QFD 8

| $\langle P(cd\mid po)\rangle_N$ | SD | po | W(Da/Si) 25 sol | $\langle P(cd\mid af)\rangle_N$ |
|---|---|---|---|---|
| 0.000531 | 0.000173 | SAS | 250 | 0.000066 |
| 0.000426 | 0.000153 | NFE | 525 | 0.000138 |
| 0.000422 | 0.000132 | EUR | 250 | 0.000066 |
| 0.000418 | 0.000151 | AFR | 700 | 0.000183 |
| 0.000415 | 0.000151 | ASJ | 525 | 0.000138 |
| 0.000415 | 0.000153 | AMR | 700 | 0.000183 |
| 0.000410 | 0.000155 | FIN | 525 | 0.000138 |
| 0.000410 | 0.000168 | FEM | 525 | 0.000138 |
| 0.000394 | 0.000162 | EAS | 700 | 0.000183 |
| 0.000371 | 0.000126 | MAL | 525 | 0.000138 |
| 0.000348 | 0.000126 | OTH | 1775 | 0.000465 |
| 0.000319 | 0.000127 | ExA | 950 | 0.000249 |

| $\langle\langle P(cd\mid po)\rangle_N\rangle_{po}$ | SD | SD% | x | R |
|---|---|---|---|---|
| 0.000407 | 0.000051 | 12.600463 | 0.010654 | −0.868032 |

Benign cd → l2 QFD 9

| $\langle P(cd\mid po)\rangle_N$ | SD | po | W(Da/Si) 25 sol | $\langle P(cd\mid af)\rangle_N$ |
|---|---|---|---|---|
| 0.001223 | 0.000191 | SAS | 200 | 0.000052 |
| 0.001126 | 0.000199 | EUR | 200 | 0.000052 |
| 0.001117 | 0.000248 | AFR | 450 | 0.000118 |
| 0.001106 | 0.000228 | NFE | 350 | 0.000092 |
| 0.001097 | 0.000221 | FIN | 350 | 0.001024 |
| 0.001095 | 0.000236 | AMR | 450 | 0.000118 |
| 0.001089 | 0.000249 | ASJ | 350 | 0.000092 |
| 0.001083 | 0.000246 | FEM | 350 | 0.000092 |
| 0.001078 | 0.000243 | EAS | 450 | 0.000118 |
| 0.001058 | 0.000252 | MAL | 350 | 0.000092 |
| 0.001012 | 0.000271 | OTH | 1175 | 0.001240 |
| 0.000974 | 0.000283 | ExA | 600 | 0.000157 |

| $\langle\langle P(cd\mid po)\rangle_N\rangle_{po}$ | SD | SD% | x | R |
|---|---|---|---|---|
| 0.001088 | 0.000061 | 5.600974 | 0.028516 | −0.904081 |

Benign cd → l5 QFD 10

| $\langle P(cd\mid po)\rangle_N$ | SD | po | W(Da/Si) 25 sol | $\langle P(cd\mid af)\rangle_N$ |
|---|---|---|---|---|
| 0.000628 | 0.000163 | AFR | 75 | 0.000020 |
| 0.000627 | 0.000155 | ASJ | 75 | 0.000020 |
| 0.000623 | 0.000143 | AMR | 75 | 0.000020 |
| 0.000618 | 0.000144 | FIN | 75 | 0.000020 |
| 0.000615 | 0.000170 | NFE | 75 | 0.000020 |
| 0.000604 | 0.000161 | EAS | 75 | 0.000020 |
| 0.000600 | 0.000162 | FEM | 75 | 0.000020 |
| 0.000568 | 0.000157 | MAL | 75 | 0.000020 |
| 0.000526 | 0.000143 | OTH | 150 | 0.000039 |
| 0.000509 | 0.000164 | ExA | 75 | 0.000020 |

| $\langle\langle P(cd\mid po)\rangle_N\rangle_{po}$ | SD | SD% | x | R |
|---|---|---|---|---|
| 0.000592 | 0.000043 | 7.290267 | 0.012924 | −0.901992 |

Benign cd → l6

| $\langle P(cd\mid po)\rangle_N$ | SD | po | W(Da/Si) 25 sol | $\langle P(cd\mid af)\rangle_N$ |
|---|---|---|---|---|
| 0.000241 | 0.000106 | SAS | 75 | 0.000020 |
| 0.000186 | 0.000069 | EUR | 75 | 0.000020 |
| 0.000182 | 0.000076 | AFR | 275 | 0.000072 |
| 0.000181 | 0.000073 | NFE | 275 | 0.000072 |
| 0.000179 | 0.000070 | ASJ | 275 | 0.000072 |
| 0.000174 | 0.000068 | FIN | 275 | 0.000072 |
| 0.000173 | 0.000062 | FEM | 275 | 0.000072 |
| 0.000172 | 0.000067 | OTH | 500 | 0.000131 |
| 0.000172 | 0.000069 | MAL | 275 | 0.000072 |
| 0.000165 | 0.000054 | AMR | 275 | 0.000072 |
| 0.000163 | 0.000072 | EAS | 275 | 0.000072 |
| 0.000151 | 0.000064 | ExA | 425 | 0.000111 |

| $\langle\langle P(cd\mid po)\rangle_N\rangle_{po}$ | SD | SD% | x | R |
|---|---|---|---|---|
| 0.000178 | 0.000022 | 12.287257 | 0.004673 | −0.796597 |

Benign cd → l8 QFD 11

| $\langle P(cd\mid po)\rangle_N$ | SD | po | W(Da/Si) 25 sol | $\langle P(cd\mid af)\rangle_N$ |
|---|---|---|---|---|
| 0.000542 | 0.000170 | NFE | 25 | 0.000007 |
| 0.000533 | 0.000168 | AFR | 25 | 0.000007 |
| 0.000532 | 0.000164 | ASJ | 25 | 0.000007 |
| 0.000527 | 0.000155 | FIN | 25 | 0.000007 |
| 0.000522 | 0.000161 | AMR | 25 | 0.000007 |
| 0.000513 | 0.000147 | FEM | 25 | 0.000007 |
| 0.000512 | 0.000148 | EAS | 25 | 0.000007 |
| 0.000485 | 0.000147 | MAL | 25 | 0.000007 |
| 0.000463 | 0.000144 | OTH | 25 | 0.000007 |
| 0.000423 | 0.000145 | ExA | 50 | 0.000013 |

| $\langle\langle P(cd\mid po)\rangle_N\rangle_{po}$ | SD | SD% | x | R |
|---|---|---|---|---|
| 0.000505 | 0.000038 | 7.429435 | 0.011031 | −0.908323 |

Benign cd → lx QFD 12

| $\langle P(cd\mid po)\rangle_N$ | SD | po | W(Da/Si) 25 sol | $\langle P(cd\mid af)\rangle_N$ |
|---|---|---|---|---|
| 0.000660 | 0.000159 | AFR | 100 | 0.000026 |
| 0.000656 | 0.000150 | AMR | 100 | 0.000026 |
| 0.000651 | 0.000154 | ASJ | 100 | 0.000026 |
| 0.000646 | 0.000183 | NFE | 100 | 0.000026 |
| 0.000636 | 0.000142 | FIN | 100 | 0.000026 |
| 0.000636 | 0.000150 | FEM | 100 | 0.000026 |
| 0.000627 | 0.000173 | EAS | 100 | 0.000026 |
| 0.000596 | 0.000151 | MAL | 100 | 0.000026 |
| 0.000544 | 0.000175 | OTH | 200 | 0.000052 |
| 0.000532 | 0.000152 | ExA | 75 | 0.000020 |

| $\langle\langle P(cd\mid po)\rangle_N\rangle_{po}$ | SD | SD% | x | R |
|---|---|---|---|---|
| 0.000619 | 0.000046 | 7.473512 | 0.013506 | −0.902909 |

Benign cd → pr

| $\langle P(cd\mid po)\rangle_N$ | SD | po | W(Da/Si) 25 sol | $\langle P(cd\mid af)\rangle_N$ |
|---|---|---|---|---|
| 0.000483 | 0.000200 | SAS | 100 | 0.000026 |
| 0.000379 | 0.000138 | NFE | 350 | 0.000092 |
| 0.000372 | 0.000142 | FIN | 350 | 0.000092 |
| 0.000368 | 0.000149 | AFR | 350 | 0.000092 |
| 0.000364 | 0.000126 | EUR | 100 | 0.000026 |
| 0.000360 | 0.000147 | ASJ | 350 | 0.000092 |
| 0.000354 | 0.000157 | EAS | 350 | 0.000092 |
| 0.000349 | 0.000133 | FEM | 350 | 0.000092 |
| 0.000334 | 0.000124 | OTH | 1025 | 0.000269 |
| 0.000332 | 0.000148 | MAL | 350 | 0.000092 |
| 0.000329 | 0.000105 | AMR | 350 | 0.000092 |
| 0.000299 | 0.000102 | ExA | 375 | 0.000098 |

| $\langle\langle P(cd\mid po)\rangle_N\rangle_{po}$ | SD | SD% | x | R |
|---|---|---|---|---|
| 0.000360 | 0.000045 | 12.451186 | 0.009442 | −0.838763 |

Benign cd → s1 QFD 13

| $\langle P(cd\mid po)\rangle_N$ | SD | po | W(Da/Si) 25 sol | $\langle P(cd\mid af)\rangle_N$ |
|---|---|---|---|---|
| 0.000736 | 0.000152 | SAS | 75 | 0.000020 |
| 0.000633 | 0.000142 | EUR | 75 | 0.000020 |
| 0.000627 | 0.000161 | AFR | 100 | 0.000026 |
| 0.000624 | 0.000139 | ASJ | 25 | 0.000007 |
| 0.000622 | 0.000190 | NFE | 25 | 0.000007 |
| 0.000612 | 0.000159 | FIN | 25 | 0.000007 |
| 0.000609 | 0.000158 | AMR | 100 | 0.000026 |
| 0.000603 | 0.000145 | FEM | 25 | 0.000007 |
| 0.000586 | 0.000177 | EAS | 100 | 0.000026 |
| 0.000574 | 0.000172 | MAL | 25 | 0.000007 |
| 0.000513 | 0.000173 | OTH | 100 | 0.000026 |
| 0.000488 | 0.000159 | ExA | 150 | 0.000039 |

| $\langle\langle P(cd\mid po)\rangle_N\rangle_{po}$ | SD | SD% | x | R |
|---|---|---|---|---|
| 0.000602 | 0.000062 | 10.308242 | 0.015783 | −0.898658 |

Benign cd → s2 QFD 14

| $\langle P(cd\mid po)\rangle_N$ | SD | po | W(Da/Si) 25 sol | $\langle P(cd\mid af)\rangle_N$ |
|---|---|---|---|---|
| 0.000768 | 0.000163 | SAS | 50 | 0.000013 |
| 0.000662 | 0.000136 | EUR | 50 | 0.000013 |
| 0.000653 | 0.000175 | AFR | 125 | 0.000965 |
| 0.000637 | 0.000186 | NFE | 100 | 0.000026 |
| 0.000635 | 0.000164 | ASJ | 100 | 0.000026 |
| 0.000633 | 0.000183 | AMR | 125 | 0.000033 |
| 0.000627 | 0.000164 | FIN | 100 | 0.000026 |
| 0.000623 | 0.000156 | EAS | 125 | 0.000033 |
| 0.000617 | 0.000171 | FEM | 100 | 0.000026 |
| 0.000597 | 0.000167 | MAL | 100 | 0.000026 |
| 0.000550 | 0.000163 | OTH | 150 | 0.000039 |
| 0.000512 | 0.000186 | ExA | 175 | 0.000046 |

| $\langle\langle P(cd\mid po)\rangle_N\rangle_{po}$ | SD | SD% | x | R |
|---|---|---|---|---|
| 0.000626 | 0.000062 | 9.885447 | 0.016407 | −0.886831 |

Benign cd → s3 QFD 15

| $\langle P(cd\mid po)\rangle_N$ | SD | po | W(Da/Si) 25 sol | $\langle P(cd\mid af)\rangle_N$ |
|---|---|---|---|---|
| 0.000596 | 0.000175 | AFR | 50 | 0.000013 |
| 0.000591 | 0.000155 | ASJ | 50 | 0.000013 |
| 0.000583 | 0.000177 | NFE | 50 | 0.000013 |
| 0.000580 | 0.000163 | FIN | 50 | 0.000013 |
| 0.000574 | 0.000175 | FEM | 50 | 0.000013 |
| 0.000574 | 0.000175 | EAS | 50 | 0.000013 |
| 0.000567 | 0.000181 | AMR | 50 | 0.000013 |
| 0.000549 | 0.000164 | MAL | 50 | 0.000013 |
| 0.000475 | 0.000138 | OTH | 200 | 0.000052 |
| 0.000464 | 0.000176 | ExA | 125 | 0.000033 |

| $\langle\langle P(cd\mid po)\rangle_N\rangle_{po}$ | SD | SD% | x | R |
|---|---|---|---|---|
| 0.000555 | 0.000047 | 8.496645 | 0.012126 | −0.861313 |

Benign cd → s4 QFD 16

| $\langle P(cd\mid po)\rangle_N$ | SD | po | W(Da/Si) 25 sol | $\langle P(cd\mid af)\rangle_N$ |
|---|---|---|---|---|
| 0.000642 | 0.000171 | AFR | 25 | 0.000007 |
| 0.000635 | 0.000174 | NFE | 25 | 0.000007 |
| 0.000631 | 0.000157 | ASJ | 25 | 0.000007 |
| 0.000631 | 0.000173 | FIN | 25 | 0.000007 |
| 0.000619 | 0.000188 | FEM | 25 | 0.000007 |
| 0.000617 | 0.000183 | AMR | 25 | 0.000007 |
| 0.000615 | 0.000174 | EAS | 25 | 0.000007 |
| 0.000590 | 0.000176 | MAL | 25 | 0.000007 |
| 0.000527 | 0.000183 | ExA | 50 | 0.000013 |
| 0.000518 | 0.000151 | OTH | 100 | 0.000026 |

| $\langle\langle P(cd\mid po)\rangle_N\rangle_{po}$ | SD | SD% | x | R |
|---|---|---|---|---|
| 0.000602 | 0.000045 | 7.386635 | 0.013157 | −0.875683 |

Benign cd → s5

| $\langle P(cd\mid po)\rangle_N$ | SD | po | W(Da/Si) 25 sol | $\langle P(cd\mid af)\rangle_N$ |
|---|---|---|---|---|
| 0.000512 | 0.000157 | AFR | 75 | 0.000020 |
| 0.000509 | 0.000156 | NFE | 75 | 0.000020 |
| 0.000497 | 0.000153 | ASJ | 75 | 0.000020 |
| 0.000496 | 0.000134 | FIN | 75 | 0.000020 |
| 0.000495 | 0.000167 | AMR | 75 | 0.000020 |
| 0.000490 | 0.000153 | FEM | 75 | 0.000020 |
| 0.000464 | 0.000134 | MAL | 75 | 0.000020 |
| 0.000462 | 0.000154 | EAS | 75 | 0.000020 |
| 0.000400 | 0.000124 | OTH | 175 | 0.000046 |

| $\langle\langle P(cd\mid po)\rangle_N\rangle_{po}$ | SD | SD% | x | R |
|---|---|---|---|---|
| 0.000481 | 0.000035 | 7.234359 | 0.009444 | −0.867346 |

Benign cd → s6 QFD 17

| $\langle P(cd\mid po)\rangle_N$ | SD | po | W(Da/Si) 25 sol | $\langle P(cd\mid af)\rangle_N$ |
|---|---|---|---|---|
| 0.000753 | 0.000172 | SAS | 25 | 0.000007 |
| 0.000631 | 0.000157 | AFR | 75 | 0.000020 |
| 0.000627 | 0.000155 | EUR | 25 | 0.000007 |
| 0.000623 | 0.000169 | ASJ | 75 | 0.000020 |
| 0.000623 | 0.000197 | NFE | 75 | 0.000020 |
| 0.000615 | 0.000171 | FIN | 75 | 0.000020 |
| 0.000612 | 0.000182 | AMR | 75 | 0.000020 |
| 0.000608 | 0.000165 | FEM | 75 | 0.000020 |
| 0.000596 | 0.000182 | EAS | 75 | 0.000020 |
| 0.000578 | 0.000152 | MAL | 75 | 0.000020 |
| 0.000527 | 0.000164 | OTH | 150 | 0.000039 |
| 0.000490 | 0.000153 | ExA | 75 | 0.000020 |

| $\langle\langle P(cd\mid po)\rangle_N\rangle_{po}$ | SD | SD% | x | R |
|---|---|---|---|---|
| 0.000607 | 0.000063 | 10.402641 | 0.015903 | −0.868606 |

Benign cd → s7 QFD 18

| $\langle P(cd\mid po)\rangle_N$ | SD | po | W(Da/Si) 25 sol | $\langle P(cd\mid af)\rangle_N$ |
|---|---|---|---|---|
| 0.000633 | 0.000146 | NFE | 25 | 0.000007 |
| 0.000631 | 0.000174 | AFR | 25 | 0.000007 |
| 0.000628 | 0.000165 | ASJ | 25 | 0.000007 |
| 0.000626 | 0.000137 | FIN | 25 | 0.000007 |
| 0.000626 | 0.000165 | AMR | 25 | 0.000007 |
| 0.000606 | 0.000179 | FEM | 25 | 0.000007 |
| 0.000604 | 0.000153 | EAS | 25 | 0.000007 |
| 0.000578 | 0.000164 | MAL | 25 | 0.000007 |
| 0.000529 | 0.000155 | OTH | 125 | 0.000033 |
| 0.000504 | 0.000170 | ExA | 25 | 0.000007 |

| $\langle\langle P(cd\mid po)\rangle_N\rangle_{po}$ | SD | SD% | x | R |
|---|---|---|---|---|
| 0.000597 | 0.000046 | 7.669611 | 0.013026 | −0.886841 |

Benign cd → s8 QFD 19

| $\langle P(cd\mid po)\rangle_N$ | SD | po | W(Da/Si) 25 sol | $\langle P(cd\mid af)\rangle_N$ |
|---|---|---|---|---|
| 0.000545 | 0.000233 | NFE | 100 | 0.000026 |
| 0.000518 | 0.000195 | ASJ | 100 | 0.000026 |
| 0.000513 | 0.000163 | FIN | 100 | 0.000026 |
| 0.000513 | 0.000183 | AFR | 100 | 0.000026 |
| 0.000504 | 0.000160 | AMR | 100 | 0.000026 |
| 0.000503 | 0.000198 | EAS | 100 | 0.000026 |
| 0.000492 | 0.000174 | FEM | 100 | 0.000026 |
| 0.000472 | 0.000167 | MAL | 100 | 0.000026 |
| 0.000441 | 0.000175 | OTH | 250 | 0.000066 |
| 0.000405 | 0.000153 | ExA | 275 | 0.000072 |

| $\langle\langle P(cd\mid po)\rangle_N\rangle_{po}$ | SD | SD% | x | R |
|---|---|---|---|---|
| 0.000491 | 0.000041 | 8.349245 | 0.010712 | −0.924135 |

Benign cd → z1 QFD 20

| $\langle P(cd\mid po)\rangle_N$ | SD | po | W(Da/Si) 25 sol | $\langle P(cd\mid af)\rangle_N$ |
|---|---|---|---|---|
| 0.000676 | 0.000138 | AFR | 25 | 0.000007 |
| 0.000668 | 0.000163 | ASJ | 25 | 0.000007 |
| 0.000664 | 0.000159 | AMR | 25 | 0.000007 |
| 0.000658 | 0.000151 | NFE | 25 | 0.000007 |
| 0.000644 | 0.000152 | EAS | 25 | 0.000007 |
| 0.000638 | 0.000139 | FIN | 25 | 0.000007 |
| 0.000633 | 0.000144 | FEM | 25 | 0.000007 |
| 0.000610 | 0.000133 | MAL | 25 | 0.000007 |
| 0.000566 | 0.000142 | OTH | 25 | 0.000007 |
| 0.000549 | 0.000148 | ExA | 50 | 0.000013 |

| $\langle\langle P(cd\mid po)\rangle_N\rangle_{po}$ | SD | SD% | x | R |
|---|---|---|---|---|
| 0.000631 | 0.000043 | 6.861804 | 0.013770 | −0.935326 |

{−0.935326, to , −0.715106}
