## Supplementary material for "Demographic Model for Inheritable Cardiac Disease": actc1summary

{ACTC1 Pathogenic Summary} {ACTC1 Benign Summary}

| Pathogenic | cd → d1 | | | |
|---|---|---|---|---|
| $\langle P(cd\|po)\rangle_N$ | SD | po | W(Da/Si) 25 sol | $\langle P(cd\|af)\rangle_N$ |
| 0.001153 | 0.000228 | OTH | 550 | 0.000045 |
| $\langle\langle P(cd\|po)\rangle_N\rangle_{po}$ | SD | SD% | x | Σx |
| 0.001153 | 0 | 0.000000 | 0.011689 | 0.014478 |

| Benign | cd → d1 | | | |
|---|---|---|---|---|
| $\langle P(cd\|po)\rangle_N$ | SD | po | W(Da/Si) 25 sol | $\langle P(cd\|af)\rangle_N$ |
| 0.000275 | 0.000228 | OTH | 550 | 0.000045 |
| $\langle\langle P(cd\|po)\rangle_N\rangle_{po}$ | SD | SD% | x | R |
| 0.000275 | 0 | 0.000000 | 0.002789 | 0 |

| Pathogenic | cd → d3 | | | |
|---|---|---|---|---|
| $\langle P(cd\|po)\rangle_N$ | SD | po | W(Da/Si) 25 sol | $\langle P(cd\|af)\rangle_N$ |
| 0.001166 | 0.000186 | FEM | 25 | 0.000002 |
| 0.000850 | 0.000200 | OTH | 550 | 0.000045 |
| $\langle\langle P(cd\|po)\rangle_N\rangle_{po}$ | SD | SD% | x | Σx |
| 0.001008 | 0.000223 | 22.121230 | 0.020433 | 0.028979 |

| Benign | cd → d3 | | | |
|---|---|---|---|---|
| $\langle P(cd\|po)\rangle_N$ | SD | po | W(Da/Si) 25 sol | $\langle P(cd\|af)\rangle_N$ |
| 0.000578 | 0.000200 | OTH | 550 | 0.000045 |
| 0.000265 | 0.000188 | FEM | 25 | 0.000002 |
| $\langle\langle P(cd\|po)\rangle_N\rangle_{po}$ | SD | SD% | x | R |
| 0.000422 | 0.000221 | 52.519892 | 0.008546 | 0 |

| Pathogenic | cd → h2 | | | |
|---|---|---|---|---|
| $\langle P(cd\|po)\rangle_N$ | SD | po | W(Da/Si) 25 sol | $\langle P(cd\|af)\rangle_N$ |
| 0.001207 | 0.000158 | OTH | 25 | 0.000002 |
| $\langle\langle P(cd\|po)\rangle_N\rangle_{po}$ | SD | SD% | x | Σx |
| 0.001207 | 0 | 0.000000 | 0.012234 | 0.014478 |

| Benign | cd → h2 | | | |
|---|---|---|---|---|
| $\langle P(cd\|po)\rangle_N$ | SD | po | W(Da/Si) 25 sol | $\langle P(cd\|af)\rangle_N$ |
| 0.000221 | 0.000158 | OTH | 25 | 0.000002 |
| $\langle\langle P(cd\|po)\rangle_N\rangle_{po}$ | SD | SD% | x | R |
| 0.000221 | 0 | 0.000000 | 0.002244 | 0 |

| Pathogenic | cd → h6 | | | |
|---|---|---|---|---|
| $\langle P(cd\|po)\rangle_N$ | SD | po | W(Da/Si) 25 sol | $\langle P(cd\|af)\rangle_N$ |
| 0.001287 | 0.000117 | AFR | 25 | 0.000002 |
| 0.001253 | 0.000140 | FEM | 25 | 0.000002 |
| 0.001247 | 0.000121 | AMR | 25 | 0.000002 |
| 0.001244 | 0.000135 | EAS | 25 | 0.000002 |
| $\langle\langle P(cd\|po)\rangle_N\rangle_{po}$ | SD | SD% | x | Σx |
| 0.001258 | 0.000020 | 1.583082 | 0.050985 | 0.057829 |

| Benign | cd → h6 | | | |
|---|---|---|---|---|
| $\langle P(cd\|po)\rangle_N$ | SD | po | W(Da/Si) 25 sol | $\langle P(cd\|af)\rangle_N$ |
| 0.000181 | 0.000136 | EAS | 25 | 0.000002 |
| 0.000179 | 0.000120 | AMR | 25 | 0.000002 |
| 0.000175 | 0.000142 | FEM | 25 | 0.000002 |
| 0.000140 | 0.000116 | AFR | 25 | 0.000002 |
| $\langle\langle P(cd\|po)\rangle_N\rangle_{po}$ | SD | SD% | x | R |
| 0.000169 | 0.000020 | 11.653823 | 0.006844 | −0.875782 |

| Pathogenic | cd → hb | | | |
|---|---|---|---|---|
| $\langle P(cd\|po)\rangle_N$ | SD | po | W(Da/Si) 25 sol | $\langle P(cd\|af)\rangle_N$ |
| 0.001179 | 0.000231 | OTH | 125 | 0.000010 |
| $\langle\langle P(cd\|po)\rangle_N\rangle_{po}$ | SD | SD% | x | Σx |
| 0.001179 | 0 | 0.000000 | 0.011944 | 0.014478 |

| Benign | cd → hb | | | |
|---|---|---|---|---|
| $\langle P(cd\|po)\rangle_N$ | SD | po | W(Da/Si) 25 sol | $\langle P(cd\|af)\rangle_N$ |
| 0.000250 | 0.000231 | OTH | 125 | 0.000010 |
| $\langle\langle P(cd\|po)\rangle_N\rangle_{po}$ | SD | SD% | x | R |
| 0.000250 | 0 | 0.000000 | 0.002534 | 0 |

| Pathogenic | cd → rr | | | |
|---|---|---|---|---|
| $\langle P(cd\|po)\rangle_N$ | SD | po | W(Da/Si) 25 sol | $\langle P(cd\|af)\rangle_N$ |
| 0.001293 | 0.000085 | AFR | 25 | 0.000002 |
| 0.001280 | 0.000095 | EAS | 25 | 0.000002 |
| 0.001278 | 0.000107 | SAS | 25 | 0.000002 |
| 0.001254 | 0.000106 | AMR | 25 | 0.000002 |
| 0.001253 | 0.000143 | EUR | 25 | 0.000002 |
| 0.001226 | 0.000161 | OTH | 50 | 0.000004 |
| $\langle\langle P(cd\|po)\rangle_N\rangle_{po}$ | SD | SD% | x | Σx |
| 0.001264 | 0.000024 | 1.927794 | 0.076856 | 0.086731 |

| Benign | cd → rr | | | |
|---|---|---|---|---|
| $\langle P(cd\|po)\rangle_N$ | SD | po | W(Da/Si) 25 sol | $\langle P(cd\|af)\rangle_N$ |
| 0.000202 | 0.000161 | OTH | 50 | 0.000004 |
| 0.000169 | 0.000105 | AMR | 25 | 0.000002 |
| 0.000168 | 0.000140 | EUR | 25 | 0.000002 |
| 0.000150 | 0.000094 | EAS | 25 | 0.000002 |
| 0.000147 | 0.000102 | SAS | 25 | 0.000002 |
| 0.000137 | 0.000087 | AFR | 25 | 0.000002 |
| $\langle\langle P(cd\|po)\rangle_N\rangle_{po}$ | SD | SD% | x | R |
| 0.000162 | 0.000023 | 14.327382 | 0.009875 | −0.968309 |

| Pathogenic | cd → s6 | | | |
|---|---|---|---|---|
| $\langle P(cd\|po)\rangle_N$ | SD | po | W(Da/Si) 25 sol | $\langle P(cd\|af)\rangle_N$ |
| 0.001208 | 0.000185 | OTH | 25 | 0.000002 |
| $\langle\langle P(cd\|po)\rangle_N\rangle_{po}$ | SD | SD% | x | Σx |
| 0.001208 | 0 | 0.000000 | 0.012246 | 0.014478 |

| Benign | cd → s6 | | | |
|---|---|---|---|---|
| $\langle P(cd\|po)\rangle_N$ | SD | po | W(Da/Si) 25 sol | $\langle P(cd\|af)\rangle_N$ |
| 0.000220 | 0.000185 | OTH | 25 | 0.000002 |
| $\langle\langle P(cd\|po)\rangle_N\rangle_{po}$ | SD | SD% | x | R |
| 0.000220 | 0 | 0.000000 | 0.002232 | 0 |

| Pathogenic | cd → sy | | | |
|---|---|---|---|---|
| $\langle P(cd\|po)\rangle_N$ | SD | po | W(Da/Si) 25 sol | $\langle P(cd\|af)\rangle_N$ |
| 0.001218 | 0.000172 | OTH | 25 | 0.000002 |
| $\langle\langle P(cd\|po)\rangle_N\rangle_{po}$ | SD | SD% | x | Σx |
| 0.001218 | 0 | 0.000000 | 0.012343 | 0.014478 |

| Benign | cd → sy | | | |
|---|---|---|---|---|
| $\langle P(cd\|po)\rangle_N$ | SD | po | W(Da/Si) 25 sol | $\langle P(cd\|af)\rangle_N$ |
| 0.000211 | 0.000172 | OTH | 25 | 0.000002 |
| $\langle\langle P(cd\|po)\rangle_N\rangle_{po}$ | SD | SD% | x | R |
| 0.000211 | 0 | 0.000000 | 0.002136 | 0 |

{ACTC1 R-correlation range over domains} {−0.968309, to , −0.875782}
